# Total Biosynthesis of Triacsin Featuring an *N*-hydroxytriazene Pharmacophore

**DOI:** 10.1101/2021.05.12.443849

**Authors:** Antonio Del Rio Flores, Frederick F. Twigg, Yongle Du, Wenlong Cai, Daniel Q. Aguirre, Michio Sato, Moriel J. Dror, Maanasa Narayanamoorthy, Jiaxin Geng, Nicholas A. Zill, Wenjun Zhang

## Abstract

Triacsins are an intriguing class of specialized metabolites possessing a conserved *N*-hydroxytriazene moiety not found in any other known natural products. Triacsins are notable as potent acyl-CoA synthetase inhibitors in lipid metabolism, yet their biosynthesis has remained elusive. Through extensive mutagenesis and biochemical studies, we here report all enzymes required to construct and install the *N*-hydroxytriazene pharmacophore of triacsins. Two distinct ATP-dependent enzymes were revealed to catalyze the two consecutive N-N bond formation reactions, including a glycine-utilizing hydrazine-forming enzyme, Tri28, and a nitrous acid-utilizing *N*-nitrosating enzyme, Tri17. This study paves the way for future mechanistic interrogation and biocatalytic application of enzymes for N-N bond formation.

## Introduction

The triacsins (**1-4**) are a unique class of natural products containing an 11-carbon alkyl chain and a terminal *N*-hydroxytriazene moiety (**Fig. 1**). Originally discovered from *Streptomyces aureofaciens* as vasodilators through an antibiotic screening program^1^, the triacsins are most notable for their use as acyl-CoA synthetase inhibitors by mimicking fatty acids to study lipid metabolism^2–11^. Recently, triacsin C has also shown promise to inhibit SARS-CoV-2-replication, where morphological effects were observed at viral replication centers^12^. Despite potent activities, the triacsin biosynthetic pathway, particularly the enzymatic machinery to install the *N*-hydroxytriazene pharmacophore, remain obscure.

**Figure 1.**
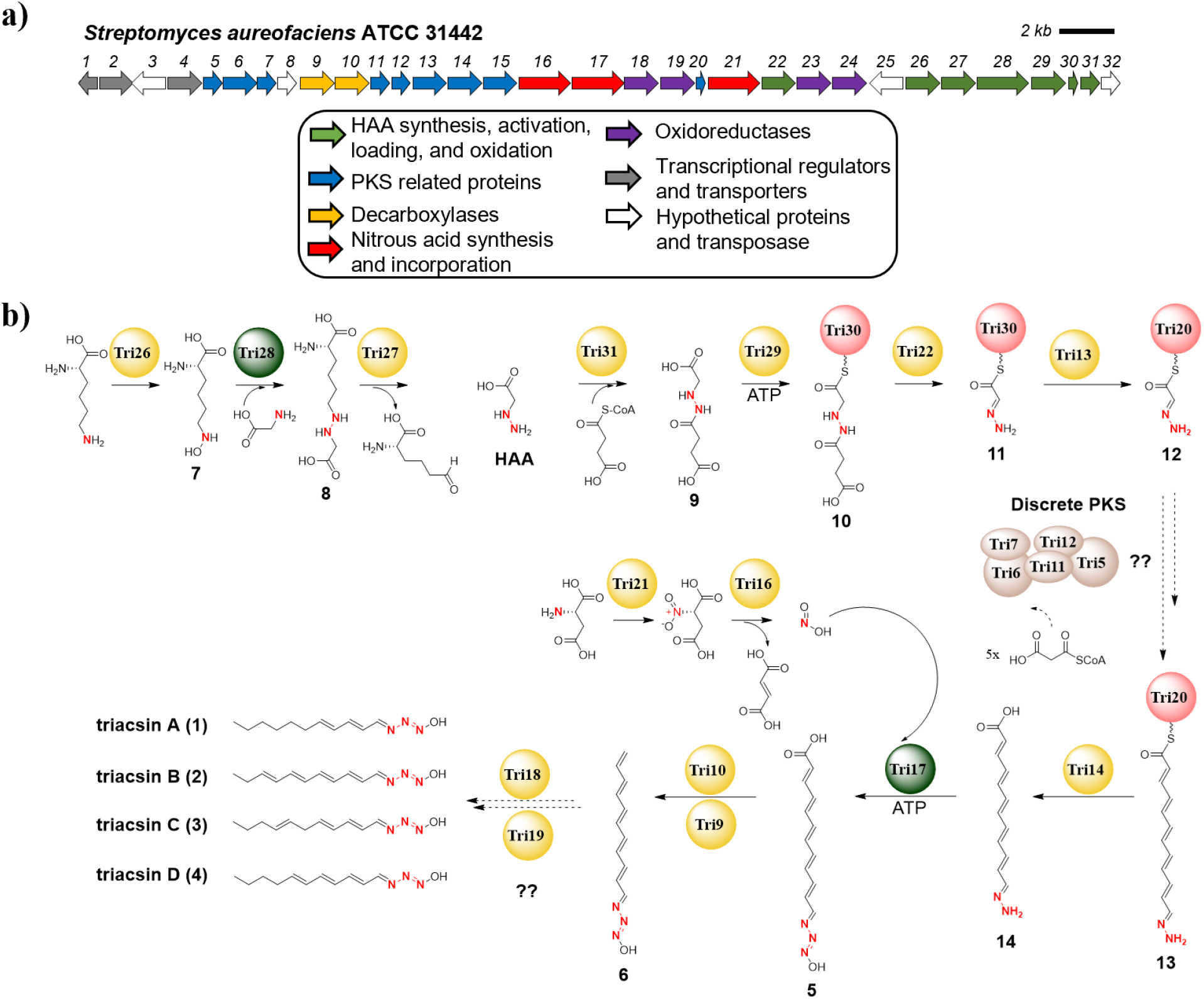
Biosynthesis of triacsins (1-4). a) Schematic of the triacsin BGC in *S. aureofaciens* ATCC 31442. b) The proposed biosynthetic pathway of triacsins based on this work. Tri28 and Tri17 (highlighted in green) are the two ATP-utilizing enzymes which catalyze formation of the two different N-N bonds found in triacsins.

The *N*-hydroxytriazene moiety, containing three consecutive nitrogen atoms, is rare in nature and has only been identified in the triacsin family of natural products^13^. In contrast to the prevalence of N-N linkages in synthetic drug libraries, the enzymes for N-N bond formation have only started to be revealed recently^14,15^. Examples include a heme enzyme (KtzT) in kutzneride biosynthesis^16^, a fusion protein (Spb40) consisting of a cupin and a methionyl-tRNA synthetase (metRS)-like domain in s56-p1 biosynthesis^17^, an *N*-nitrosating metalloenzyme (SznF) in streptozotocin biosynthesis^18^, and a transmembrane protein (AzpL) in 6-diazo-5-oxo-L-norleucine formation^19^ (**Supplementary Fig. 1**). Investigation of the biosynthesis of triacsins that contain two consecutive N-N bonds may determine if known mechanisms are used or if yet new enzymes for N-N bond formation exist.

To shed light on triacsin biosynthesis, we identified the triacsin biosynthetic gene cluster (*tri*1-32) in *S. aureofaciens* through genome mining and mutagenesis studies^20^. Based on bioinformatic analyses and labeled precursor feeding experiments, we further proposed a chemical logic for *N*-hydroxytriazene biosynthesis via a key intermediate, hydrazinoacetic acid (HAA) as well as nitrous acid dependent N-N bond formation^20^. In this present study, we scrutinized the activity of 15 enzymes for triacsin biosynthesis through biochemical analyses. We revealed all enzymes required to construct and install the unusual *N*-hydroxytriazene pharmacophore, highlighting a new ATP-dependent, nitrous acid-utilizing *N*-nitrosating enzyme.

## Results

### *N*-hydroxytriazene assembly starts before PKS extension

Although we inferred from bioinformatic analysis that the acyl portion of the triacsin scaffold is derived from discrete polyketide synthases (PKSs), it was unclear the reaction timing of how nitrous acid and HAA pathways are coordinated with the purported acyl intermediate to form the triacsin scaffold with the *N*-hydroxytriazene moiety^20^. One hypothesis is that HAA, once loaded onto an acyl carrier protein (ACP), is modified by nitrous acid to form an ACP-bound intermediate which resembles the final *N*-hydroxytriazene moiety that is further elongated by PKSs. Alternatively, the *N*-hydroxytriazene moiety may serve as a PKS chain terminating agent rather than a “starter unit”. The overall odd number of carbon chain length was also puzzling, which was likely due to a decarboxylation event on the PKS chain since the carboxylate carbon of glycine, a predicted precursor of HAA, was retained (**Extended Data Fig. 1a**). While mutants with each of the 25 *tri* genes individually disrupted did not accumulate any biosynthetic intermediates or shunt products, the *Δtri9-10* mutant strain, in which the two genes encoding a decarboxylase and its accessory enzyme were both deleted, abolished the production of triacsins and accumulated a new metabolite (**5**) (**Fig. 2a**). **5** exhibited a distinct UV profile and was predicted to have a molecular formula of C12H13N3O3 based on high-resolution mass spectrometry (HRMS) analysis (**Supplementary Fig. 2**, **Supplementary Note 1**). From a 12-L fermentation culture of *Δtri9-10*, 2.5 mg of **5** was obtained as a yellow and amorphous powder. Subsequent one-dimensional (1D) and two-dimensional (2D) nuclear magnetic resonance (NMR) analysis revealed **5** to be a new triacsin-like metabolite with a 12-carbon fully unsaturated alkyl chain terminated by a carboxylic acid moiety (**Supplementary Note 1**). **5** was unstable and >90% degraded after two days at room temperature (**Supplementary Fig. 2**). The presence of both the *N*-hydroxytriazene moiety and the terminal carboxylic acid in **5** argues against the possibility of the *N*-hydroxytriazene moiety as a PKS chain terminating agent. We thus propose that the *N*-hydroxytriazene assembly starts before PKS extension (**Fig. 1**).

**Figure 2.**
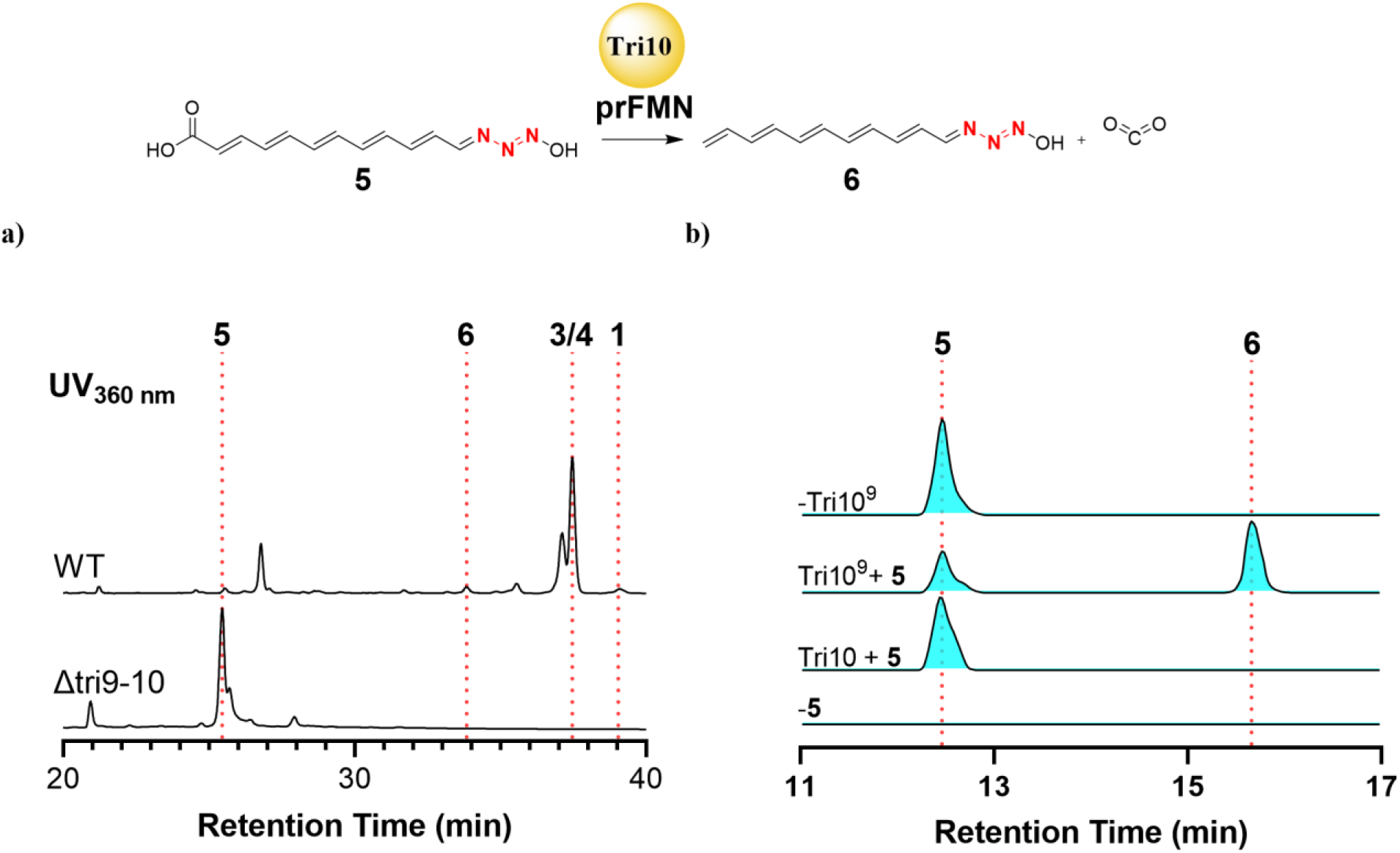
In vivo and in vitro analysis of decarboxylase. a) HPLC/UV trace (360 nm) demonstrating accumulation of a new metabolite (**5**) in the *Δtri9-10* mutant and abolishment of triacsins. b) Biochemical analysis of Tri10^9^. Extracted ion chromatograms (EICs) showing that Tri10 purified from *E. coli* converted **5** to **6** only when co-expressed with Tri9. The calculated masses for **5**: m/z=248.1030 ([M+H]^+^) and **6**: m/z=204.1132 ([M+H]^+^) are used for each trace with a 10-ppm mass error tolerance. At least three independent replicates were performed for each assay, and representative results are shown.

To confirm that **5** is a direct substrate for Tri10, a UbiD family of non-oxidative decarboxylase, Tri10 was overexpressed in *E. coli* and purified for biochemical analysis (**Supplementary Fig. 3**). Notably, Tri9 is homologous to UbiX, a flavin prenyltransferase that modifies FMN using an isoprenyl unit to generate prFMN, a required co-factor for the activity of UbiD^21–24^. As expected, the purified Tri10 alone showed no activity toward **5** presumably due to the absence of the required co-factor (**Fig. 2b**). In contrast, the purified Tri10 that was co-expressed with Tri9 in *E. coli* successfully decarboxylated **5** based on HRMS analysis with the concomitant production of CO_2_ (**Fig. 2b, Extended Data Fig. 1b**). While the full structural characterization of the decarboxylated product of **5** was not feasible due to instability of both substrate and product, the UV profile and HRMS data of the reaction product are consistent with a decarboxylated product **6** (**Supplementary Fig. 4**). In addition, **6** was also identified as a minor metabolite from the culture broth of the wild-type *S. aureofaciens* (**Fig. 2a**, **Supplementary Fig. 4**). Considering that only **5** with a fully unsaturated alkyl chain was accumulated in *Δtri9-10*, we propose that **6** is likely a precursor for all observed triacsin congeners varying in the unsaturation pattern of their alkyl chains (**Fig. 1**).

### Succinylation is required for HAA activation and loading onto ACP

We previously proposed that hydrazinoacetic acid (HAA), presumably generated through the activities of Tri26-28, is a key intermediate for triacsin biosynthesis^20^. Specifically, *N*^6^-hydroxy-lysine (**7**) that is generated by Tri26 (an *N*-hydroxylase homolog), reacts with glycine, promoted by Tri28 (a cupin and metRS didomain protein), to form *N*^6^- (carboxymethylamino)lysine (**8**), which then undergoes an oxidative cleavage to yield HAA and 2-aminoadipate 6-semialdehyde catalyzed by Tri27, an FAD-dependent d-amino acid oxidase homolog. The Tri26-28 coupled biochemical assay successfully produced HAA by comparing to an authentic standard through 2,4-dinitrofluorobenzene (DNFB) derivatization^17,25^, confirming the predicted activity of Tri26-28 (**Fig. 3a, Supplementary Fig. 5**). In addition, Tri26 was found to only hydroxylate lysine but not ornithine, and the coupled assay with Tri26/28 generated **8** as expected, further confirming the activity of each enzyme (**Extended Data Fig. 2, Supplementary Fig. 5**).

**Figure 3.**
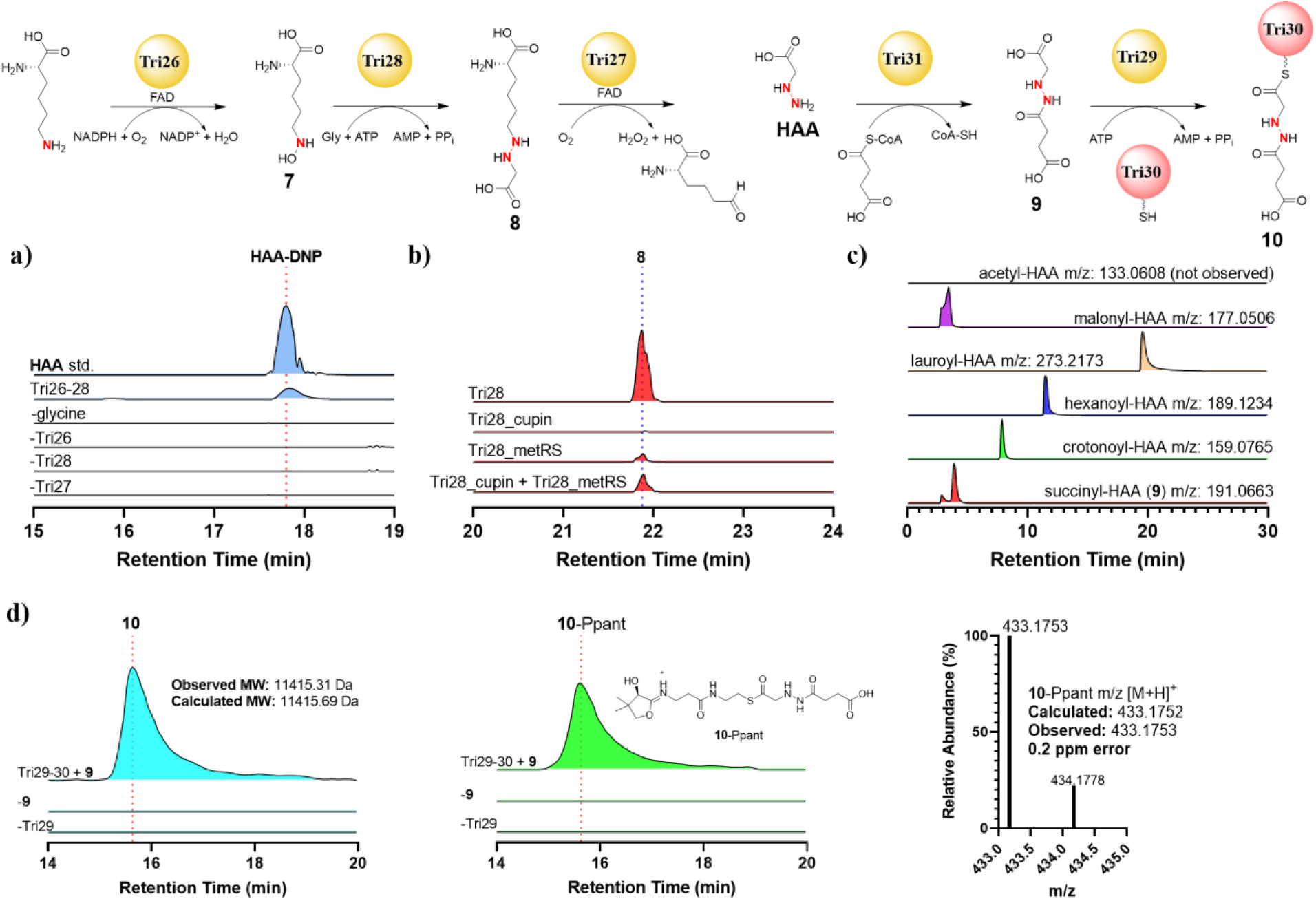
HAA synthesis, activation, and loading onto Tri30. a) EICs demonstrating production of a derivatized HAA from coupled Tri26-28 assays subjected to DNFB derivatization. Omission of any of the enzymes or glycine resulted in abolishment of HAA-DNP. The calculated mass for HAA-DNP: m/z=257.0517 ([M+H]^+^) is used for each trace. b) EICs showing production of **8** from coupled Tri26,28 assays. The cupin and metRS domains were individually dissected and tested in parallel. The calculated mass for **8**: m/z=220.1292 ([M+H]^+^) is used for each trace. c) EICs showing production of various acyl-HAA products generated by Tri31. Individual masses used for each trace are listed. d) EICs showing production of **10 (**+12 charge) along with its deconvoluted mass and Ppant fragment. Omission of Tri29 or **9** (generated in situ using Tri31) resulted in abolishment of **10**. A 10-ppm mass error tolerance was used for each trace. At least three independent replicates were performed for each assay, and representative results are shown.

We next dissected Tri28 to obtain insight into the roles of the cupin and metRS domains in forming the N-N bond between **7** and glycine. Each domain was individually purified from *E. coli* and re-combining each domain in the in vitro assay showed successful, albeit less efficient, **8** formation than the intact didomain enzyme, demonstrating the feasibility of domain dissection (**Fig. 3b**). Surprisingly, the metRS domain alone was sufficient in generating **8** without the cupin domain, although addition of the cupin domain increased the titer of the product (**Fig. 3b**).

After confirming the production of HAA, we next probed the fate of HAA in *N*-hydroxytriazene assembly. *Tri26-28* is a part of six-gene operon of *tri26-31* that are homologous to *spb38-43* in s56-p1 biosynthesis^17^, whereas the functions of *tri29-31*, putatively encoding an AMP-dependent synthetase, an ACP and an *N*-acetyltransferase, respectively, remained unknown for both triacsin and s56-p1 biosynthesis. We hypothesized that Tri29 activates HAA through adenylation and then loads it onto Tri30. Tri31 possibly acetylates the distal nitrogen of the purported hydrazine moiety as a protecting group, similar to the activity of FzmQ/KinN^26,27^. A biochemical assay with Tri29 and the holo-ACP, Tri30, yielded no HAA-*S*-Tri30 based on LC-HRMS analysis (**Supplementary Fig. 6**), suggesting that Tri31 acetylates HAA prior to the Tri29-catalyzed activation and loading onto ACP. A subsequent assay of Tri31 with HAA and acetyl-CoA, however, did not yield acetyl-HAA (**Fig. 3c**). Instead, a new product with mass consistent with succinyl-HAA (**9**), was detected in a trace amount (**Supplementary Fig. 7**). We hypothesized that **9** was generated from succinyl-CoA, which tightly bound to the Tri31 protein purified from *E. coli*. The LC-HRMS analysis of the purified Tri31 protein solution revealed a trace amount of succinyl-CoA as compared to the standard, and the addition of extra succinyl-CoA to the Tri31 assay increased the yield of **9**, consistent with our hypothesis (**Fig. 3c, Supplementary Fig. 7**).

The preference of Tri31 toward succinyl-CoA over acetyl-CoA was unexpected based on the prediction of Tri31 as an *N*-acetyltransferase. To exclude the possibility of an artifact due to co-purification of succinyl-CoA with Tri31, we next investigated the acyl-CoA substrate specificity of Tri31 using various CoA substrates. Tri31 recognized all tested acyl-CoAs except acetyl-CoA, and moderately preferred succinyl-CoA over others based on kinetic analysis (**Fig. 3c, Supplementary Fig. 8**). The promiscuity of Tri31 also made possible to confirm that Tri31 acylates the distal nitrogen of HAA, as products such as hexanoyl-HAA were more feasible to be purified from in vitro assays than **9** due to its increased hydrophobicity. The NMR analysis of hexanoyl-HAA showed a ^3^*J* ^1^*H*-^15^N HMBC cross-peak between the nitrogen bearing methylene (*δ*_H_ 3.69) and the amide nitrogen (*δ*_N_ 144.7), as well as a ^2^*J* ^1^H-^15^N HMBC cross-peak between the amide -NH- (*δ*_H_ 9.68) and the other nitrogen (*δ*_N_ 61.7), supporting that the distal nitrogen of HAA is connected to the carbonyl carbon of the CoA substrate (**Supplementary Note 2**).

The activation and loading of all acyl-HAAs generated by Tri31 were then tested using a coupled assay containing Tri29-31. The succinyl-HAA loaded Tri30 (**10**) was successfully generated based on both the intact protein HRMS and the phosphopantetheine (Ppant) ejection assay (**Fig. 3d**). Interestingly, none of the other acyl-HAAs were loaded onto Tri30 (**Supplementary Fig. 9**), indicating that Tri29 acts as a gatekeeper enzyme in the biosynthetic pathway and succinylation is required for HAA activation and loading onto ACP (**Fig. 1**).

### Hydrazone “starter unit” for PKS extension

Tri22, an acyl-CoA dehydrogenase homolog, was proposed to oxidize **10** as the next step in *N*-hydroxytriazene formation. Tri22 was recombinantly expressed in *E. coli* as a yellow protein containing FAD (**Supplementary Fig. 10**). An *in vitro* assay of Tri22 with **10** generated in situ by Tri29-31 resulted in conversion of **10** to a product with mass consistent of **11** (**Fig. 4a**). To confirm the proposed structure of **11**, the reaction product was subjected to base hydrolysis to release the free acid that was further derivatized by *ortho*-phthalaldehyde (OPTA) for LC-UV-HRMS analysis^28,29^. The expected acid product, 2-hydrazineylideneacetic acid (2-HYAA), was also chemical synthesized and derivatized by OPTA. By comparison, the structure of **11** was confirmed to be 2-HYAA attached to Tri30 (**Extended Data Fig. 3, Supplementary Note 3, Supplementary Fig. 11**). It is notable that Tri22 showed no activity toward the free acid **9** (**Supplementary Fig. 11**), demonstrating the necessity of a carrier protein for Tri22 recognition.

**Figure 4.**
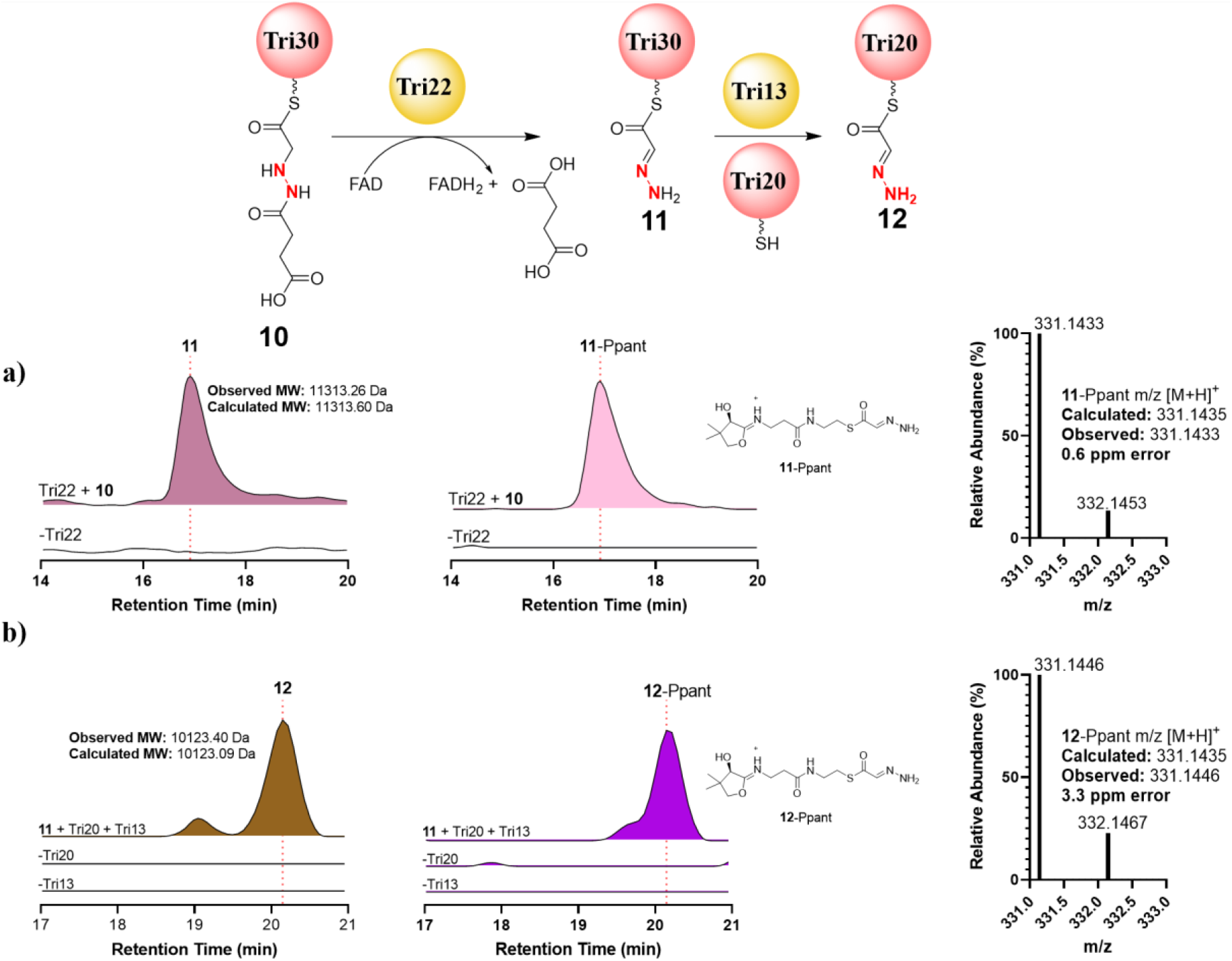
Hydrazone “starter unit” formation for PKS extension. a) Biochemical analysis of Tri22. EICs demonstrating production of **11 (**+12 charge) along with its deconvoluted mass and Ppant fragment. Omission of Tri22 results in abolishment of **11**. **10** was generated in situ using Tri29-31. b) Hydrazone translocation catalyzed by Tri13. EICs showing production of **12 (**+6 charge) along with its deconvoluted mass and Ppant fragment. Omission of Tri20 or Tri13 resulted in abolishment of **12**. A 10-ppm mass error tolerance was used for each trace. At least three independent replicates were performed for each assay, and representative results are shown.

**11** was proposed to be the first possible substrate to react with the presumed nitrous acid for *N*-hydroxytriazene formation during triacsin biosynthesis. We then biochemically reconstituted the activity of Tri16 (a lyase) and Tri21 (a flavin-dependent enzyme), the CreD and CreE homologs^30^, respectively, to confirm that nitrous acid can be generated by the pathway enzymes. After confirming the production of nitrous acid by Tri16/21 (**Supplementary Fig. 12**), we set to search for a nitrous acid-utilizing enzyme that modifies **11** to form *N*-hydroxytriazene. There was weak evidence that an acyl-CoA ligase homolog, CreM, promoted the reaction of an aryl amine with nitrous acid to form diazo in cremeomycin biosynthesis^31^, although CreM seemed not required for the diazo formation^30^. The triacsin biosynthetic gene cluster encodes a CreM homolog, Tri17 (40% sequence similarity), which was essential for triacsin biosynthesis based on the mutagenesis study^20^. However, an in vitro assay of Tri17 with nitrite, ATP, and **11** yielded no new product. Notably, AMP was formed in the presence of nitrite and Tri17, suggesting that Tri17 may activate nitrite using ATP (**Supplementary Fig. 13**).

Although enzyme(s) other than Tri17 were likely responsible for the third nitrogen addition, we reasoned that **11** might not be a correct substrate and the *N*-hydroxytriazene formation might take place at a later stage, after PKS extension and release of the polyketide intermediate. While the discrete nature and lack of some essential enzymes (likely shared with fatty acid biosynthesis in the native producer) made the in vitro reconstitution of the entire PKS machinery a formidable task, the biochemical assay containing Tri13, a ketosynthase homolog, and Tri20, a free-standing ACP, demonstrated that Tri13 promoted translocation of the 2-HYAA moiety from Tri30 to Tri20 to yield **12** (**Fig. 4b**). This observation suggests that the PKS extension likely takes place on Tri20 with a hydrazone “starter unit”. After five rounds of extension to yield **13**, the 12-carbon fully unsaturated alkyl chain would be released as a carboxylic acid to produce **14 (Fig. 1**). Tri14, a peptidase homolog, may play a thioesterase role to facilitate this hydrolytic release of the polyketide product. To probe the activity of Tri14, we purified a soluble version of Tri14 from *E. coli* upon cleavage of its predicted *N*-terminal recognition sequence for self-activation^32^. Due to the unavailability of **13**, surrogate substrates such as hexanoyl-*S*-Tri20 and lauroyl-*S*-Tri20 were generated using Sfp, a promiscuous Ppant transferase^33^, and used in the Tri14 assays (**Supplementary Fig. 14**). Tri14 exhibited a hydrolytic activity toward lauroyl-*S*-Tri20 but not hexanoyl-*S*-Tri20, demonstrating an acyl chain length preference (**Extended Data Fig. 4**). In addition, Tri14 showed no activity toward lauroyl-*S*-Tri30, consistent with the hypothesis that PKS extension occurs on Tri20 instead of Tri30 (**Extended Data Fig. 4, Supplementary Fig. 14**). These results strongly suggest that Tri14 is a plausible enzyme to catalyze the hydrolytic release of **14** from **13 (Fig. 1**).

### Tri17 completes *N*-hydroxytriazene assembly post PKS

Structural comparison of **14** and **5** suggests that the third nitrogen addition could occur on **14** post the PKS assembly line. Considering that **14** is likely more unstable than **5** as suggested by literature^34^, we turned to a surrogate substrate, **15**, to probe the activity of Tri17 (**Fig. 5a**). **15** was chemically synthesized (**Supplementary Note 4**) and used in the Tri17 assay together with ATP and nitrite. A new product was successfully produced which was identified to be triacsin A (**1**) by comparing to the authentic standard isolated from the wild-type culture of *S. aureofaciens* (**Fig. 5a, Supplementary Fig. 15, Supplementary Note 5**). This product was not observed in negative controls where each individual component such as Tri17, ATP, nitrite, or **15** was omitted (**Fig. 5a**). The utilization of ^15^N-nitrite also led to the expected mass spectral shift of the product (**Fig. 5a**), further confirming the utilization of nitrite by Tri17 in the N-N bond formation. The production of AMP was slightly increased in the presence of **15**, consistent with the proposed reaction mechanism of Tri17 in which **15** nucleophilically attacks the ATP-activated nitrite to yield **1** with the release of AMP (**Extended Data Fig. 5**).

**Figure 5.**
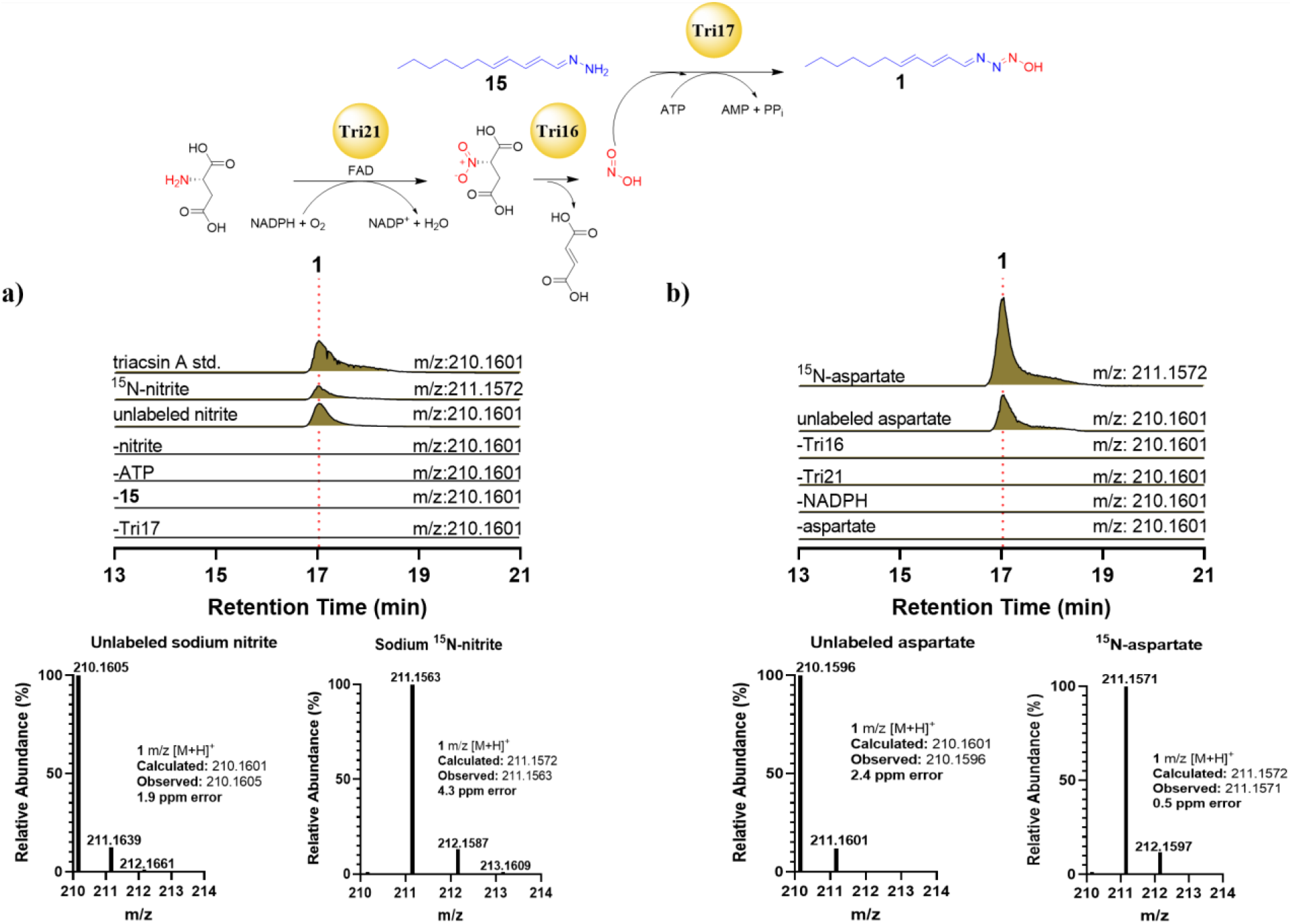
Biochemical analysis of Tri17. a) EICs showing the production of **1** from assays containing Tri17, ATP, nitrite, and **15**. Omission of any of these components led to abolishment of **1**. Utilization of ^15^N-nitrite resulted in the expected mass spectral shift. b) EICs demonstrating the production of **1** from a coupled Tri16, 21,17 assay containing aspartate, **15**, NADPH, and ATP. Omission of any of these components led to abolishment of **1**. Utilization of ^15^N-aspartate resulted in the expected mass spectral shift. A 10-ppm mass error tolerance was used for each trace. At least three independent replicates were performed for each assay, and representative results are shown.

We next probed whether nitrous acid, generated enzymatically by Tri16/21, could be utilized by Tri17. A biochemical assay containing Tri16, Tri21 and Tri17 with aspartic acid, NADPH, ATP and **15** resulted in **1** production, and the utilization of ^15^N-aspartic acid led to the expected mass spectral shift of **1** (**Fig. 5b**), confirming the utilization of nitrous acid generated by Tri16/21 in *N*-hydroxytriazene biosynthesis. Notably, no spontaneous formation of **1** was observed using either nitrite or enzymatically generated nitrous acid in the absence of Tri17 (**Fig. 5a and 5b**). We thus deduce that in triacsin biosynthesis, Tri17 takes nitrous acid and **14** as substrates, and catalyzes the production of **5** in an ATP-dependent manner (**Fig. 1**). Tri17 appeared to be promiscuous toward acyl chain modifications as both **14** and **15** could be recognized. However, neither 2-HYAA nor 12-aminododecanoic acid was recognized by Tri17 based on comparative metabolomics analysis (**Supplementary Fig. 16**), suggesting the activity of Tri17 is both chain length and “hydrazone” specific. Notably, Tri17 did not utilize nitrate to modify **15 (Supplementary Fig. 16**), indicating that Tri17 is a dedicated nitrous acid-utilizing *N*-nitrosating enzyme.

## Discussion

Our extensive bioinformatics, mutagenesis, and biochemical analysis allow a first proposal of the complex biosynthetic pathway for triacsins, particularly the *N*-hydroxytriazene moiety unique to this class of natural products (**Fig. 1**). The biosynthesis starts with the production of HAA from lysine and glycine catalyzed by Tri26-28, which is then succinylated by a succinyltransferase, Tri31, and activated and loaded onto Tri30 promoted by an acyl-ACP ligase, Tri29. Tri22, an acyl-ACP dehydrogenase, catalyzes desuccinylation and oxidation of HAA to a hydrazone moiety attached to Tri30, which is then translocated to another free-standing ACP, Tri20, facilitated by a ketosynthase, Tri13, for PKS extension. The PKS machinery likely include ketosynthases encoded by *tri6-7*, a ketoreductase encoded by *tri5*, two dehydratases encoded by *tri11-12*, and a malonyl-CoA ACP acyltransferase that is shared with the host fatty acid synthase machinery. Tri6/7 likely form heterodimers in which Tri6 performs the canonical Claisen-like decarboxylative condensation, while Tri7, lacking an active site cysteine, acts as a chain length factor that controls the number of iterative elongation steps^35–37^. Tri11/12 also likely dimerize to form a functional dehydratase. After five iterations of ketosynthase, reductase and dehydratase, a fully unsaturated polyketide chain is hydrolytically released from Tri20 promoted by a thioesterase, Tri14. Tri17, an ATP-dependent *N*-nitrosating enzyme, recognizes this highly unstable biosynthetic intermediate, **14**, and catalyzes the formation of **5** to finish *N*-hydroxytriazene assembly using nitrous acid generated by Tri16/21. **5** subsequently undergoes non-oxidative decarboxylation promoted by Tri9/10 and various reduction/oxidation reactions, likely catalyzed by Tri18/19, two remaining oxidoreductases encoded by the gene cluster, to yield triacsin congeners varying in the unsaturation pattern of their alkyl chains (**Fig. 1**). This proposed biosynthetic pathway is consistent with failure in detecting intermediate accumulation in most of the gene deletion mutants as most of the presumed biosynthetic intermediates are either attached to an ACP or highly unstable.

The biosynthesis of triacsins features two distinct mechanisms for the two consecutive N-N bond formations. One involves Tri28, a cupin and metRS didomain protein, which catalyzes the formation of *N6*-(carboxymethylamino)lysine (**8**) from *N6*-hydroxy-lysine (**7**) and glycine. Previous work established a proposed reaction scheme for Spb40, a homolog of Tri28 (75% sequence similarity), using filtered culture broth from *E. coli* expressing the recombinant Spb40 and isotope-labeled substrates^17^. However, the utilization of whole cell lysate instead of purified enzymes precluded the detailed biochemical characterization of this enzyme, leaving questions such as the tRNA requirement and the true functions of each of two domains unanswered. Using purified Tri28 and the dissected individual domains, we revealed that tRNA is not required for the activity of Tri28. Glycine is thus most likely activated by ATP to form glycyl-AMP which is subjected to a nucleophilic attack by **7** to form an ester intermediate (**Supplementary Fig. 17**). Surprisingly, the *N*-terminal cupin domain is also not required for **8** formation, which is inconsistent with prior mutagenesis results where *E. coli* producing the Spb40 mutant with the substitution of alanine for His45, one of the key residues for metal-binding of the cupin domain that is also conserved in Tri28, abolished the ability to produce **8**^17^. Considering that the cupin domain increased the catalytic efficiency in vitro (**Fig. 3b**), we hypothesize that this cupin domain promotes the rearrangement of the putative ester intermediate in N-N bond formation, and this reactivity might be important in vivo. Further biochemical and structural analysis will provide more insights into the catalytic mechanism of this intriguing family of enzymes.

This study also identifies a new *N*-nitrosating enzyme, Tri17, which catalyzes *N*-hydroxytriazene formation from nitrous acid and acyl hydrazone in an ATP-dependent way. Tri17 is homologous to an acyl-CoA ligase but CoA is not a required substrate, suggesting that an AMP derivative is a plausible biosynthetic intermediate (**Extended Data Fig. 5**). Although nitrous acid has been implicated to react with an amine for N-N bond formation in multiple natural product biosynthesis, such as cremeomycin^30,31^, fosfazinomycin^27^, kinamycin^26^, alanosine^38,39^, and alazopeptin^19^, the underlying enzymology often remains elusive, lacking a definitive biochemical characterization using purified enzymes. For example, CreM, a homolog of Tri17, was previously reported to catalyze diazotization in cremeomycin biosynthesis, although the evidence is weak because of its low solubility in *E. coli* and the high reactivity of nitrous acid to nonenzymatically react with an aryl amine to form a diazo moiety^27,31^. Similarly, non-enzymatic *N*-nitrosation using nitrous acid could occur in alanosine and fragin biosynthesis^39,40^, although other possibilities such as a fusion protein (AlnA) consisting of a cupin and an AraC-like DNA-binding domain, or a CreM homolog encoded elsewhere on the genome was also proposed to be an N-N bond forming enzyme without evidence^38,39^. AzpL, a transmembrane protein with no homology to any characterized protein, was recently revealed to catalyze diazotization in alazopeptin biosynthesis through mutagenesis and heterologous expression, but no in vitro biochemical evidence was obtained due to the membrane protein nature^19^. Other enzyme candidates that may catalyze N-N bond formation using nitrous acid were also proposed in fosfazinomycin and kinamycin biosynthesis (FzmP and KinJ, respectively), but these hypothetical proteins have not been studied^26,27^. The discovery of Tri17 provides the first solid example of a nitrous acid-utilizing enzyme required for N-N bond formation and provides the basis for future mechanistic understanding of this new family of dedicated *N*-nitrosating enzymes.

In conclusion, through identification of a highly unstable biosynthetic intermediate from mutagenesis studies and biochemical analysis of 15 enzymes required for triacsin biosynthesis, we have revealed the exquisite timing and mechanisms of the biosynthesis of this intriguing class of natural products. Two distinct ATP-dependent enzymes have been revealed to catalyze the two consecutive N-N bond formation of *N*-hydroxytriazene, including a glycine-utilizing hydrazine-forming enzyme and a nitrous acid-utilizing *N*-nitrosating enzyme. This study paves the way for future discovery, mechanistic understanding, and biocatalytic application of enzymes for N-N bond formation.

## Acknowledgements

We would like to thank Tzu-Yang Huang and Bryan McCloskey at UC Berkeley for their assistance with initial GC-MS experiments and consultation in designing experiments. We would like to also thank Dr. Miao Zhang from the UC Berkeley Catalysis Center for training with the GC-MS apparatus and aiding with data interpretation. The GC-MS work was made possible by the Catalysis Facility of Lawrence Berkeley National Laboratory, supported by the Director, Office of Science, of the U.S. Department of Energy (contract no. DE-AC02-05CH11231. This research was financially supported by grants to W.Z. from the NIH (R01GM136758 and DP2AT009148) and the Chan Zuckerberg Biohub Investigator Program. We would like to also thank Zhijuan Hu and Rui Zhai for helpful discussions regarding the complex biosynthesis of triacsins. Lastly, we would like to thank Yuanbo Shen for assisting in protein purification and kinetic assays.

## Author contributions

A.D.R.F designed the experiments, performed all *in vitro* experiments with purified enzymes, analyzed the data, and wrote the manuscript. F.F.T designed experiments to detect the biosynthetic intermediate, constructed plasmids for protein expression in *E. coli*, and analyzed data. Y.D. analyzed all the NMR data. W.C. performed the GC-MS experiments and helped analyze NMR data. D.Q.A helped purify the biosynthetic intermediate. M.J.D, M.N., and J.G. aided in protein purification, constructing plasmids, and aided in repeating biochemical assays for this study. N.A.Z performed the comparative metabolomics work. W.Z. designed the experiments, analyzed the data, and wrote the manuscript.

## Online Methods

### Materials

Phusion High-Fidelity PCR Master Mix (NEB) was used for PCR reactions. Restriction enzymes were purchased from Thermo Scientific. All chemicals used in this work were obtained from Alfa Aesar, Sigma-Aldrich or Fisher Scientific, unless otherwise noted. CoA substrates (purity >95%) were obtained from CoALA Biosciences. Hydrazinoacetic acid HCl salt (purity 95%) was purchased from BOC Sciences. ^15^N-aspartic acid (purity 98%) and ^15^N-sodium nitrite (purity >98%) were purchased from Cambridge Isotope Laboratories, Inc.

### Bacterial Strains and Growth Conditions

The wild-type and *Δtri9-10* gene disruption mutant of *Streptomyces aureofaciens* ATCC 31442 were cultured on ISP4 agar plates to allow sporulation. Individual spore colonies were inoculated into a 2 mL seed culture of R5 liquid media and grown at 30°C and 240 rpm for 48 hours. The seed culture was inoculated into 30 mL R5 liquid media and grown for 5 days at 30°C and 240 rpm before cell harvesting. Culturing was performed in 125 mL Erlenmeyer vessels with a wire coil placed at the bottom to encourage aeration. Large scale culturing of WT *S. aureofaciens* was conducted as previously reported^20^. For large scale culturing of the *Δtri9-10* mutant, the 2 mL seed culture would instead be inoculated in 1 L R5 liquid media and grown in the same conditions. *Escherichia coli* strains were cultivated on lysogeny broth (LB) agar plates or in LB liquid media. Growth media was supplied with antibiotics as required at the following concentrations: kanamycin (50 μg/mL), apramycin (50 μg/mL), ampicillin (100 μg/mL).

### LC-HRMS Analysis of Triacsins and 5

Streptomycete cultures were pelleted by centrifugation (4,000 x g for 15 min), and the spent media was extracted 1:1 by volume with ethyl acetate. Ethyl acetate was removed by rotary evaporation, and the dry extract was dissolved in 300 μL of methanol for liquid chromatography-mass spectrometry (LC-MS) analysis by 10 μL injection onto an Agilent Technologies 6120 Quadrupole LC-MS instrument with an Agilent Eclipse Plus C18 column (4.6 x 100 mm). For high resolution MS (HRMS) analysis, the extract in methanol could be diluted by a factor of 5 for LC-HRMS and HRMS/MS analysis by 10 μL injection onto an Agilent Technologies 6545 Q-TOF LC-MS instrument with the same column. Linear gradients of 5-95% acetonitrile (vol/vol) in water with 0.1% formic acid (vol/vol) at 0.5 mL/min were used.

### Production and Purification of 5 and 1

After the 5-day culture period for the *Δtri9-10* mutant of *S. aureofaciens*, the cultures were pelleted by centrifugation (4,000 x g for 15 min), and the spent media was extracted 1:1 by volume with ethyl acetate. The ethyl acetate was removed by rotary evaporation, and the dry extract was dissolved in a 1:1 mixture (vol/vol) of dichloromethane and methanol. The dissolved extract was loaded onto a size exclusion column packed with Sephadex LH-20 (Sigma-Aldrich) and manually fractionated in a 1:1 dichloromethane and methanol running solvent. The fractions were screened by LC-MS with a diode array detector (DAD). Fractions containing the compound determined to be **5** were subjected to high performance liquid chromatography (HPLC) using an Agilent 1260 HPLC and an Atlantis T3 OBD Prep column (100 Å pore size, 5 μm particle size, 10 x 150 mm). A water/acetonitrile mobile phase buffered to a pH of 7.2 with ammonium bicarbonate was used to prevent exposing the fractions to acidic conditions. Linear gradients of 5-50% acetonitrile (vol/vol) at 2.5 mL/min were used for initial purification. Final purification of **5** was performed using an isocratic gradient at 14% acetonitrile using the same mobile phase, column, and flow rate. All fractions containing the product were kept covered to avoid light exposure and stored under N_2_ gas whenever practically achievable to avoid chemical degradation. Purified products were dried and analyzed by LC-MS and NMR. All NMR spectra were recorded on a Bruker AVANCE at 900 MHz (1H NMR) and 226 MHz (13C NMR). The production and purification procedure of **1** was conducted as previously reported^20^.

### Labeled Precursor Feeding of 1-^13^C-glycine

*S. aureofaciens* was cultured as described in a previous section. 24 hours after the 30 mL inoculation, the culture was supplied with 10 mM 1-^13^C-glycine. Compound extraction and LC-HRMS analysis were performed as described above.

### Construction of Plasmids for Expression in *E. coli*

Three vectors were used for *E. coli* induced expression of recombinant proteins using polyhistidine tags. The plasmid pETDuet-1 was used for the dual expression of Tri10 and Tri9 (Tri10^9^). The *tri10* gene was amplified with HindIII and NotI sites and inserted into the first multiple cloning site (MCS) of pETDuet-1 which encodes an N-terminal polyhistidine tag. The *tri9* gene was amplified using NdeI and XhoI sites and inserted into the second MCS of pETDuet-1 that encodes a *C*-terminal S-tag derived from pancreatic ribonuclease A. The plasmid pET-24b(+) was used for the expression of Tri28, Tri28_Cupin, Tri28_metRS, Tri30, Tri20, Tri17, Tri13 and Tri14 with *C*-terminal polyhistidine tags. The NdeI and XhoI sites were used for the restriction digest and ligation-based installation of the corresponding gene. The plasmid pLATE52 was used for the expression of Tri16, Tri21, Tri26, Tri27, Tri22, Tri29, and Tri31 with *N*-terminal polyhistidine tags. The corresponding genes were amplified and inserted through ligation-independent cloning following the protocol from Thermo Fisher Scientific’s aLICator Expression System, #K1281. All genes were amplified from either *S. aurofaciens ATCC 31442* or *S. tsukubaensis NRRL 18488* as listed in **Table S2**. Oligonucleotides utilized in this study were purchased from Integrated DNA Technologies. All oligonucleotides used in this study are listed in **Table S1** and all strains and constructs are listed in **Tables S2** and **S3**.

### Expression and Purification of Recombinant Proteins

The expression and purification for all proteins used in this study followed the same general procedure for polyhistidine tag purification as detailed here. Expression strains were grown at 37°C in 1 L of LB in a shake flask supplemented with 50 μg/mL of kanamycin or 100 μg/mL of ampicillin to an OD_600_ of 0.5 at 250 rpm. The shake flask was then placed over ice for 10 mins and induced with 120 μM of isopropyl-β-d-thiogalactopyranoside (IPTG). The cells were then incubated for 16 hours at 16°C at 250 rpm to undergo protein expression. Subsequently, the cells were harvested by centrifugation (6,371 x g, 15 min, 4°C), and the supernatant was removed. The cell pellet was resuspended in 30 mL of lysis buffer (25 mM HEPES pH 8, 500 mM NaCl, 5 mM imidazole) and cells were lysed by sonication on ice. Cellular debris was removed by centrifugation (27,216 x g, 1 hour, 4°C) and the supernatant was filtered with a 0.45 μm filter before batch binding. Ni-NTA resin (Qiagen) was added to the filtrate at 2 mL/L of cell culture, and the samples were allowed to nutate for 1 hour at 4°C. The protein-resin mixture was loaded onto a gravity flow column. The flow through was discarded and the column was then washed with approximately 25 mL of wash buffer (25 mM HEPES pH 8, 100 mM NaCl, 20 mM imidazole) and tagged protein was eluted in approximately 20 mL of elution buffer (25 mM HEPES pH 8, 100 mM NaCl, 250 mM imidazole). The whole process was monitored using a Bradford assay. Purified proteins were concentrated and exchanged into exchange buffer (25 mM HEPES pH 8, 100 mM NaCl) using Amicon ultra filtration units. After two rounds of buffer exchange and concentration, the purified enzyme was removed, and glycerol was added to a final concentration of 10% (vol/vol). Enzymes were subsequently flash-frozen in liquid nitrogen and stored at −80 °C or used immediately for *in vitro* assays. The presence and purity of purified enzymes was assessed using SDS-PAGE and the concentration was determined using a NanoDrop UV-Vis spectrophotometer (Thermo Fisher Scientific).

All proteins reported in this study were purified using the HEPES pH 8 buffer system, except for Tri29 for which Tris pH 7.5 was used to avoid precipitation upon concentration. Purification buffers for Tri30 and Tri20 (ACPs) contained 1 mM TCEP. BL21 Star (DE3) was utilized for expression of all proteins. For the case of Tri30 and Tri20, BAP1 was utilized to obtain the holo-ACPs and BL21 Star (DE3) was utilized for purification of apo-ACPs.

The approximate molecular weight and yield for each protein are the following: Tri26 (52.4 kDa, 5.2 mg/L), Tri27 (40.9 kDa, 5.4 mg/L), Tri28 (74.7 kDa, 18.5 mg/L), Tri28-metRS (62.3 kDa, 6.8 mg/L), Tri28-cupin (14.1 kDa, 22.7 mg/L), Tri29 (57.5 kDa, 34.3 mg/L), Tri30 (11 kDa, 16.9 mg/L), Tri20 (9.8 kDa, 10.8 mg/L), Tri13 (34.5 kDa, 6 mg/L), Tri31 (25.1 kDa, 20.5 mg/L), Tri22 (45.9 kDa, 6.7 mg/L), Tri21 (71.8 kDa, 6.4 mg/L), Tri16 (59.6 kDa, 15.2 mg/L), Tri17 (61.5 kDa, 34 mg/L), Tri10^9^ (56.7 kDa, 4.6 mg/L), and Tri14 (27.7 kDa, 5 mg/L).

### LC-HRMS and GC-MS Analysis of Tri10^9^ *In Vitro* Reconstitution

Reactions were performed at room temperature for 30 minutes in 100 μL of 50 mM HEPES (pH 8.0) containing 20 μM Tri10 and 100 μM **5**. **5** was delivered in the form of partially purified extracts from the *tri9-10* gene disruption mutant hydrated in water with 0.5% methanol (vol/vol) to aid in solubility. The concentration of **5** in the extracts was estimated by HPLC/UV analysis at 400 nm. The final methanol concentration in the reaction volume was 0.1% (vol/vol). After the 30-minute incubation period, the reaction was quenched with two volumes of chilled methanol. The precipitated protein was removed by centrifugation (15,000 x g, 2 min) and the supernatant was used for analysis. LC-HRMS analysis was performed on an Agilent Technologies 6545 Q-TOF LC-MS equipped with an Agilent Eclipse Plus C18 column (4.6 x 100 mm). Using a water/acetonitrile mobile phase with 0.1% (vol/vol) formic acid, analysis was performed with a linear gradient of 5-95% acetonitrile at a flow rate of 0.5 mL/min. At least three independent replicates were performed for each assay, and representative results are shown.

The GC-MS procedure to detect CO_2_ was adapted from previous work^41^. GC-MS assays consisted of a 4.5 mL of reaction mixture containing 50 mM HEPES pH 8, 1 mM **5**, and 100 μM Tri10^9^. The reaction was performed in a 10-mL sealed headspace vial (Agilent) and initiated by adding the substrate. 1 mL of the headspace gas was acquired using a gastight syringe (Hamilton) and injected into Agilent 5977A GCMS system equipped with a HP-5ms column. Injector temperature was set at 120 °C, and helium was used as the carrier gas at a flowrate of 3 mL/min. The temperature gradient was as follows: 40-100°C in 5 minutes. The mass spectrometer was operated in electron ionization mode with automatically tuned parameters, and the acquired mass range was m/z = 15-100. The CO_2_ signal was confirmed using an authentic standard and by extracting m/z = 44.

### LC-HRMS Analysis of Tri26 and Tri28 *In Vitro* Reconstitution

Reactions were performed at room temperature for 30 minutes in 100 μL of 50 mM Tris pH 7.5 containing 1 mM lysine, 1 mM glycine, 2 mM NADPH, 5 mM ATP, 2 mM MgCl_2_, 20 μM Tri26, and 20 μM Tri28. After the 30-minute incubation period, the reaction was quenched with two volumes of chilled methanol. The precipitated protein was removed by centrifugation (15,000 x g, 5 min) and the supernatant was used for analysis. LC-HRMS analysis was performed on an Agilent Technologies 6545 Q-TOF LC-MS equipped with an Agilent InfinityLab Poroshell 120 HILIC-Z column (4.6 x 100 mm). A mobile phase of water/acetonitrile was buffered with 10 mM ammonium formate and titrated to a pH of 3.2 with formic acid. LC-HRMS analysis was performed with a decreasing linear gradient of 90-60% acetonitrile at a flow rate of 1.0 mL/min. The analogous assays were performed using the individual domains of Tri28. At least three independent replicates were performed for each assay, and representative results are shown.

### LC-HRMS Analysis of HAA Production by Tri26-28 with DNFB Derivatization

Reactions were performed at room temperature for 30 minutes in 100 μL of 50 mM Tris pH 7.5, 1 mM lysine, 1 mM glycine, 2 mM NADPH, 0.2 mM FAD, 5 mM ATP, 2 mM MgCl_2_, 20 μM Tri26, 20 μM Tri27, and 20 μM Tri20. The DNFB derivatization method was adapted from a previous work^25^. The reaction was then derivatized with 200 μL of 1 mM of DNFB dissolved in 200 mM borate pH 9.0 and incubated for 30 mins at 60 °C. The precipitated protein was removed by centrifugation (15,000 x g, 5 min) and the supernatant was used for analysis. LC-HRMS analysis was performed using an Agilent Technologies 6545 Q-TOF LC-MS equipped with an Agilent Eclipse Plus C18 column (4.6 x 100 mm). A water/acetonitrile mobile phase with 0.1% (vol/vol) formic acid with a linear gradient of 2-98% acetonitrile at a flow rate of 0.5 mL/min was utilized. The same derivatization method was utilized with an authentic HAA standard. At least two independent replicates were performed for each assay, and representative results are shown.

### LC-HRMS Analysis of Tri31 In Vitro Reconstitution

Reactions were performed at room temperature for 30 minutes in 100 μL of 50 mM HEPES pH 8.0 containing 1 mM succinyl-CoA, 2 mM HAA, and 50 μM Tri31. After the 30-minute incubation period, the reaction was quenched with two volumes of chilled methanol. The precipitated protein was removed by centrifugation (15,000 x g, 5 min) and the supernatant was used for analysis. LC-HRMS analysis was performed using an Agilent Technologies 6545 Q-TOF LC-MS equipped with an Agilent Eclipse Plus C18 column (4.6 x 100 mm). A water/acetonitrile mobile phase with 0.1% (vol/vol) formic acid with a linear gradient of 5-95% acetonitrile at a flow rate of 0.5 mL/min was utilized. The same methodology was performed using different CoA substrates. At least three independent replicates were performed for each assay, and representative results are shown.

### Identification of Co-Purified Succinyl-CoA in Tri31

50 μL of 1.7 mM Tri31 was quenched with 100 μL of chilled methanol. The precipitated protein was removed by centrifugation (15,000 x g, 5 min) and the supernatant was used for analysis. LC-HRMS analysis was performed on an Agilent Technologies 6545 Q-TOF LC-MS equipped with an Agilent InfinityLab Poroshell 120 HILIC-Z column (4.6 x 100 mm). A mobile phase of water/acetonitrile was buffered with 10 mM ammonium formate and titrated to a pH of 3.2 with formic acid. LC-HRMS analysis was performed with a decreasing linear gradient of 90-60% acetonitrile at a flow rate of 1.0 mL/min. An authentic succinyl-CoA standard was utilized to compare the retention time and mass spectrum to the succinyl-CoA co-purified from Tri31.

### Determination of Kinetic Parameters for CoA Substrates from Tri31 Reaction

The following procedure to obtain 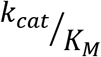 parameters for CoA substrates was adopted from previous work^19,28^. Reactions were performed at room temperature in 100 μL of 50 mM HEPES pH 8.0, 50 μM CoA substrate, 2 mM HAA, 0.1 mM 5,5’-dithio-bis(2-nitrobenzoic acid) (DTNB) and 20 μM Tri31. For each CoA substrate, a control reaction without enzyme was conducted. Activity was monitored continually over a 15-minute period by measuring the increase in absorbance at 412 nm from the reaction between the free thiol of CoASH and DNTB. Initial velocities were approximated using the extinction coefficient 14,150 *M*^−1^*cm*^−1^ and subtracting the control reactions from each substrate to correct for background activity. Each CoA substrate was tested in independent triplicate assays.

### Large scale Tri31 biochemical assay with HAA and Hexanoyl-CoA

A 1 mL reaction was performed at room temperature for 15 hours containing 50 mM HEPES pH 8.0, 5 mM HAA, 4 mM hexanoyl-CoA, and 100 μM Tri31. The reaction was extracted three times with equal volume of ethyl acetate. The organic fraction was dried over nitrogen and analyzed using NMR. **Supplementary Note 2** contains further structural characterization.

### LC-HRMS Analysis of Tri30 Loading Reaction and Ppant Ejection Assays with HAA and CoA substrates

Reactions were performed at room temperature for 20 minutes in 50 μL of 50 mM Tris pH 7.5 containing 0.2 mM succinyl-CoA, 1 mM HAA, 5 mM ATP, 2 mM MgCl_2_, 20 μM Tri31, 20 μM Tri29, and 50 μM Tri30. After the 20-minute incubation period, the reaction was diluted with 4 volumes of purified water and chilled on ice to slow further reaction progress. The diluted reaction mixture was immediately filtered by a 0.45 μm PVDF filter, and 15 μL was promptly injected onto an Agilent Technologies 6545 Q-TOF LC-MS equipped with a Phenomenex Aeris 3.6 μm Widepore XB-C18 column (2.1 x 100 mm). Using a water/acetonitrile mobile phase with 0.1% (vol/vol) formic acid, analysis was performed with a linear gradient of 30-50% acetonitrile at a flow rate of 0.25 mL/min. The molecular weights of proteins observed during electrospray ionization were determined using ESIprot to deconvolute the array of observed charge state spikes^42^. For phosphopantetheine (Ppant) ejection assays, the same conditions were used, but the source voltage was increased from 75 to 250 V to increase the fragmentation of the phosphodiester bond covalently linking the Ppant prosthetic to a holo-ACP. The same methodology was performed using different CoA substrates. To determine whether HAA can be directly loaded (without Tri31 and succinyl-CoA), the same methodology was used with the following changes: a linear gradient of 40-60% acetonitrile at a flow rate of 0.25 mL/min and the Phenomenex Aeris 3.6 μm Widepore XB-C18 column (2.1 x 250 mm) were utilized. At least three independent replicates were performed for each assay, and representative results are shown.

### LC-HRMS Analysis of Tri22 Activity on 10

Reactions were performed at room temperature for 20 minutes in 50 μL of 50 mM Tris pH 7.5 containing 0.2 mM succinyl-CoA, 1 mM HAA, 5 mM ATP, 0.2 mM FAD, 2 mM MgCl2, 20 μM Tri31, 20 μM Tri29, 20 μM Tri22 and 50 μM Tri30. After the 20-minute incubation period, the reaction was diluted with 4 volumes of purified water and chilled on ice to slow further reaction progress. The diluted reaction mixture was immediately filtered by a 0.45 μm PVDF filter, and 15 μL was promptly injected and analyzed using the same methodology in the previous section. At least three independent replicates were performed for each assay, and representative results are shown.

### Identification of FAD Tri22 Cofactor

Purified Tri22 (yellow color) was boiled for 10 minutes and precipitated protein was removed by centrifugation (15,000 x g, 5 min). The supernatant was further analyzed using LC-UV-MS with an Agilent Technologies 6545 Q-TOF LC-MS equipped with an Agilent Eclipse Plus C18 column (4.6 x 100 mm). A water/acetonitrile mobile phase with 0.1% (vol/vol) formic acid with a linear gradient of 2-98% acetonitrile at a flow rate of 0.5 mL/min was utilized. The UV was monitored at 420 nm and the mass spectrum/retention time was compared to an authentic FAD standard.

### Determination of 11 from Tri22 Reaction

The biochemical assay consisting of HAA, succinyl-CoA, HAA, Tri29-31, and Tri22 was performed as described in a previous section except for 50 μM of each enzyme was utilized and the assay was performed at a 100 μL scale. After the 20-minute incubation period, the reaction was quenched with two volumes of chilled methanol. The precipitated protein was isolated by centrifugation (15,000 x g, 5 min) and the supernatant was discarded. The protein pellet was then dissolved in 100 μL of 100 mM KOH for 10 minutes at room temperature, followed by neutralization with 100 μL of 100 mM HCl. The solution was spin filtered using a 2 kDa Amicon spin filter to remove protein residues and isolate the flowthrough. The flowthrough was subjected to OPTA derivatization^28^. The derivatization agent was prepared by mixing 1 mL of 80 mM OPTA with 98 mM of 2-ME (both dissolved in 200 mM NaOH). 100 μL of the flowthrough was combined with 100 μL of the derivatization agent. The reaction mixture was vortexed and allowed to incubate at room temperature for 2 minutes. The mixture was analyzed using LC-UV-MS with an Agilent Technologies 6545 Q-TOF LC-MS equipped with an Agilent Eclipse Plus C18 column (4.6 x 100 mm). A water/acetonitrile mobile phase with 0.1% (vol/vol) formic acid with a linear gradient of 2-98% acetonitrile at a flow rate 0.5 mL/min was utilized. 2-HYAA was chemically synthesized and subjected to the same derivatization procedure with OPTA. The mass spectrum, retention time, and UV at were compared to that of the standard. At least three independent replicates were performed for each assay, and representative results are shown.

### LC-HRMS Analysis of 2-HYAA Translocation to Tri20 with the Aid of Tri13

Reactions were performed at room temperature for 30 minutes in 50 μL of 50 mM Tris pH 7.5, 1 mM succinyl-CoA, 2 mM HAA, 5 mM ATP, 1 mM FAD, 2 mM MgCl2, 20 μM Tri31, 20 μM Tri29, 20 μM Tri22, 30 μM Tri30, 20 μM Tri20, and 10 μM Tri13. After the incubation period, the reaction was diluted with 4 volumes of purified water and chilled on ice to slow further reaction progress. The diluted reaction mixture was immediately filtered by a 0.45 μm PVDF filter, and 15 μL was promptly injected and analyzed using the same methodology in a previous section with the exception that a Phenomenex Aeris 3.6 μm Widepore XB-C18 column (2.1 x 250 mm) was used. At least two independent replicates were performed for each assay, and representative results are shown.

### Lauroyl-CoA and Hexanoyl-CoA Loading Assays on Tri30 and Tri20 using Sfp

Reactions were performed at room temperature for 30 minutes in 50 μL of 50 mM HEPES pH 8.0, 2.5 mM CoA substrate, 2 mM MgCl_2_, 100 μM ACP, and 50 μM Sfp. After the 30-minute incubation period, the reaction was diluted with 4 volumes of purified water and chilled on ice to slow further reaction progress. The diluted reaction mixture was immediately filtered by a 0.45 μm PVDF filter, and 10 μL was promptly injected onto an Agilent Technologies 6545 Q-TOF LC-MS equipped with a Phenomenex Aeris 3.6 μm Widepore XB-C18 column (2.1 x 250 mm) and Phenomenex Aeris 3.6 μm Widepore XB-C18 column (2.1 x 100 mm) for lauroyl-CoA and hexanoyl-CoA assays, respectively. Using a water/acetonitrile mobile phase with 0.1% (vol/vol) formic acid, analysis was performed with a linear gradient of 30-98% and 30-50% acetonitrile for lauroyl-CoA and hexanoyl-CoA assays, respectively at a flow rate of 0.25 mL/min. The analysis to deconvolute the protein spikes and obtain the Ppant fragment are analogous to a previous section. Hexanoyl-CoA and lauroyl-CoA were loaded to Tri20, while only lauroyl-CoA was loaded to Tri30. At least three independent replicates were performed for each assay, and representative results are shown.

### Tri14 Hydrolysis Activity Assays on Fatty Acyl-ACP Substrates

Reactions were performed at room temperature for 1 hour in 100 μL of 50 mM HEPES pH 8.0, 2.5 mM CoA substrate, 2 mM MgCl_2_, 100 μM ACP, 50 μM Tri14 and 50 μM Sfp. After the 1-hour incubation period, the reaction was quenched with two volumes of chilled methanol. The precipitated protein was removed by centrifugation (15,000 x g, 5 min) and the supernatant was used for analysis. LC-HRMS analysis was performed using an Agilent Technologies 6545 Q-TOF LC-MS equipped with an Agilent Eclipse Plus C18 column (4.6 x 100 mm). A water/acetonitrile mobile phase with 0.1% (vol/vol) formic acid with a linear gradient of 2-98% acetonitrile at a flow rate of 0.5 mL/min was utilized. Hexanoic and lauric acid standards prepared and analyzed to test for hydrolytic activity of Tri14. Assays with hexanoyl-CoA and lauroyl-CoA were utilized with Tri20, while lauroyl-CoA was utilized solely for Tri30. At least three independent replicates were performed for each assay, and representative results are shown.

### Griess Test for Nitrous Acid Production by Tri21 and Tri16

Reactions were performed at room temperature in 100 μL of 50 mM Tris (pH 7.5) containing 0.1 mM FAD, 1 mM NADPH or NADH, 1 mM aspartate, 20 μM Tri21, and 20 μM Tri16. The reaction was prepared with all reagents except NAD(P)H and aliquoted identically into 12 wells. Reactions were initiated simultaneously by the addition of NAD(P)H with a multichannel pipette. Starting with an initial time point, one reaction well was quenched every 4 minutes by the addition of one volume of Griess reagent purchased from Cell Signaling Technology.

### Tri17 Activity Assay on 15

Reactions were performed at room temperature for 30 minutes in 100 μ of 50 mM Tris pH 7.5, 1 mM **15**, 4 mM sodium nitrite (or ^15^N-sodium nitrite), 5 mM ATP, 4 mM MgCl_2_, and 100 μM Tri17. Chemically synthesized **15** was dissolved in DMSO to ensure full solubility and assays were maintained at a final concentration of 2% DMSO (vol/vol). After the incubation period, the reaction was quenched with two volumes of chilled methanol. The precipitated protein was removed by centrifugation (15,000 x g, 5 min) and the supernatant was used for analysis. LC-HRMS analysis was performed using an Agilent Technologies 6545 Q-TOF LC-MS equipped with an Agilent Eclipse Plus C18 column (4.6 x 100 mm). A water/acetonitrile mobile phase with 0.1% (vol/vol) formic acid with a linear gradient of 2-98% acetonitrile at a flow rate of 0.5 mL/min was utilized. **1** isolated from WT *S. aureofaciens* was utilized as a standard to compare the retention time, mass spectrum, and UV profiles between the biochemical assays. At least three independent replicates were performed for each assay, and representative results are shown. Analogous assays were performed with nitrate instead of nitrite and 2-HYAA or 12-aminododecanoic acid instead of **15**, respectively.

### Coupled Tri16, Tri21, and Tri17 Activity Assay

Reactions were performed at room temperature for 1 hour in 100 μL of 50 mM Tris pH 7.5, 0.1 mM FAD, 1 mM NADPH, 1 mM aspartate (or 2 mM ^15^N-aspartate), 20 μM Tri16, and 20 μM Tri21. After the incubation period, 100 μL of 50 mM Tris pH 7.5, 0.5 mM **15**, 5 mM ATP, 2 mM MgCl_2_, and 100 μM Tri17 was added to the first reaction. The coupled assay was incubated at room temperature for 16 hours. After the incubation period, the reaction was quenched with two volumes of chilled methanol. The precipitated protein was removed by centrifugation (15,000 x g, 5 min) and the supernatant was used for analysis. LC-HRMS analysis was performed using an Agilent Technologies 6545 Q-TOF LC-MS equipped with an Agilent Eclipse Plus C18 column (4.6 x 100 mm). A water/acetonitrile mobile phase with 0.1% (vol/vol) formic acid with a linear gradient of 2-98% acetonitrile at a flow rate of 0.5 mL/min was utilized. **1** isolated from WT *S. aureofaciens* was utilized as a standard to compare the retention time, mass spectrum, and UV profiles between the biochemical assays. At least three independent replicates were performed for each assay, and representative results are shown.

### Comparative Metabolomics

Tri17 assays utilizing nitrate, 2-HYAA, or 12-aminododecanoic acid were analyzed via LC-HRMS using an Agilent Technologies 6545 Q-TOF LC-MS equipped with an Agilent Eclipse Plus C18 column (4.6 x 100 mm). A water/acetonitrile mobile phase with 0.1% (vol/vol) formic acid with a linear gradient of 2-98% acetonitrile at a flow rate of 0.5 mL/min was utilized. Peak picking and comparative metabolomics were performed using MSDial with peak lists exported to Microsoft Excel.

### AMP Detection from Tri17 Assay

Reactions were performed at room temperature for 30 minutes as described in the Tri17 Activity Assay on **15** section. After the incubation period, the reaction was quenched with two volumes of chilled methanol. A similar assay was performed using **11** (generated enzymatically as described in a previous section) as a substrate instead of **15**. The precipitated protein was removed by centrifugation (15,000 x g, 5 min) and the supernatant was used for analysis. LC-HRMS analysis was performed on an Agilent Technologies 6545 Q-TOF LC-MS equipped with an Agilent InfinityLab Poroshell 120 HILIC-Z column (4.6 x 100 mm). A mobile phase of 90% acetonitrile and water were buffered with 10 mM ammonium acetate and titrated to a pH of 9.0 with ammonium hydroxide. LC-HRMS analysis was performed with a decreasing linear gradient of 81-43% acetonitrile at a flow rate of 1.0 mL/min. ATP, ADP, and AMP were also detected by monitoring the UV at 260 nm.

---

**Extended Data Figure 1.**
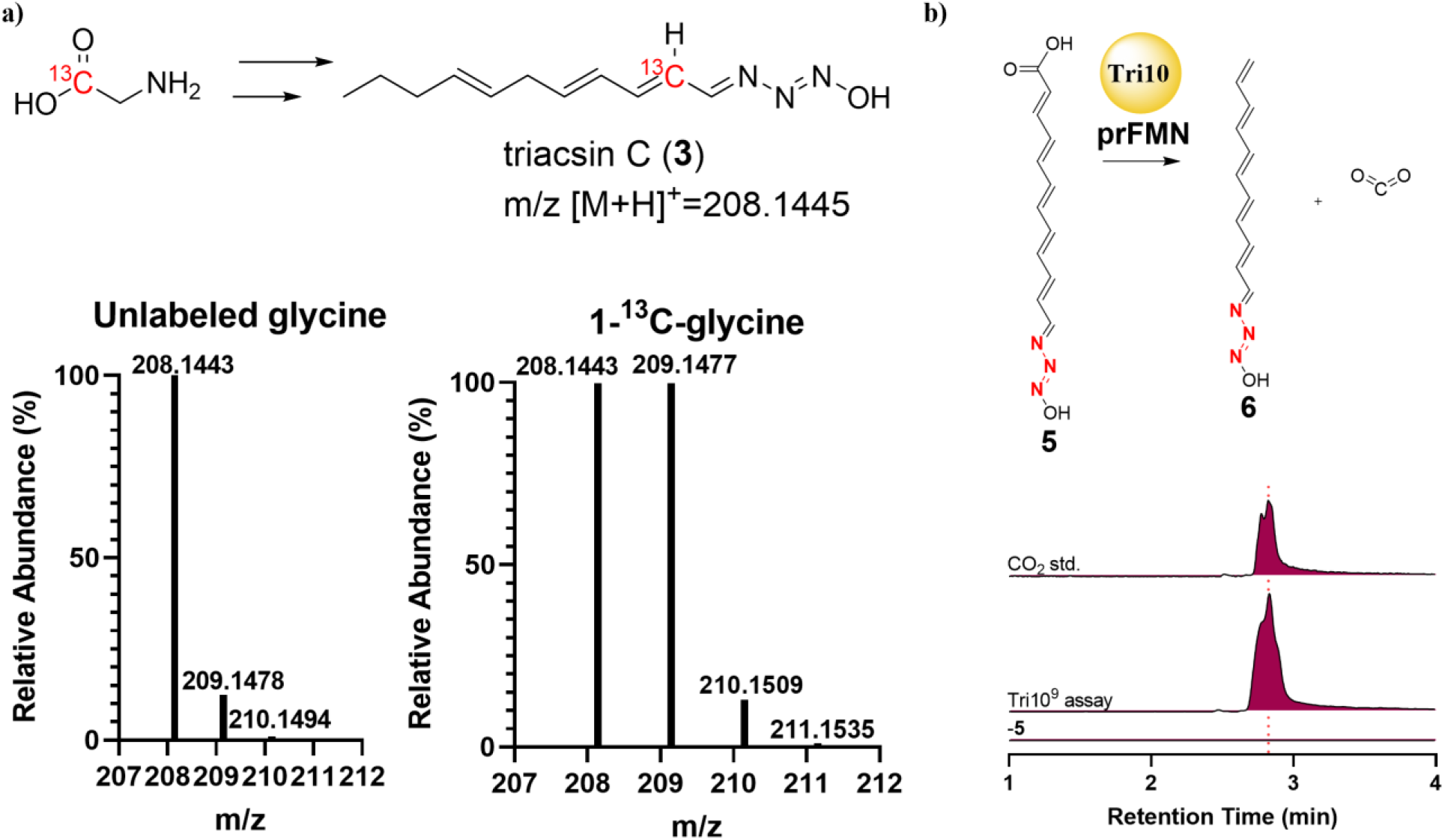
The decarboxylation event in triacsin biosynthesis. a) Feeding of unlabeled and labeled glycine to the triacsin producing cultures. HRMS analysis of **3** from a *S. aureofaciens* culture provided with 10 mM 1-^13^C-glycine demonstrated that the carboxylate carbon of glycine was retained in triacsin C. b) Detection of CO_2_ production from the Tri10^9^ biochemical assay. GC-MS chromatograms extracted with m/z=44, demonstrating CO_2_ production from a Tri10^9^ assay containing **5** as compared with an authentic standard. A 10-ppm mass error tolerance was used for each trace. At least two independent replicates were performed for each assay, and representative results are shown.

**Extended Data Figure 2.**
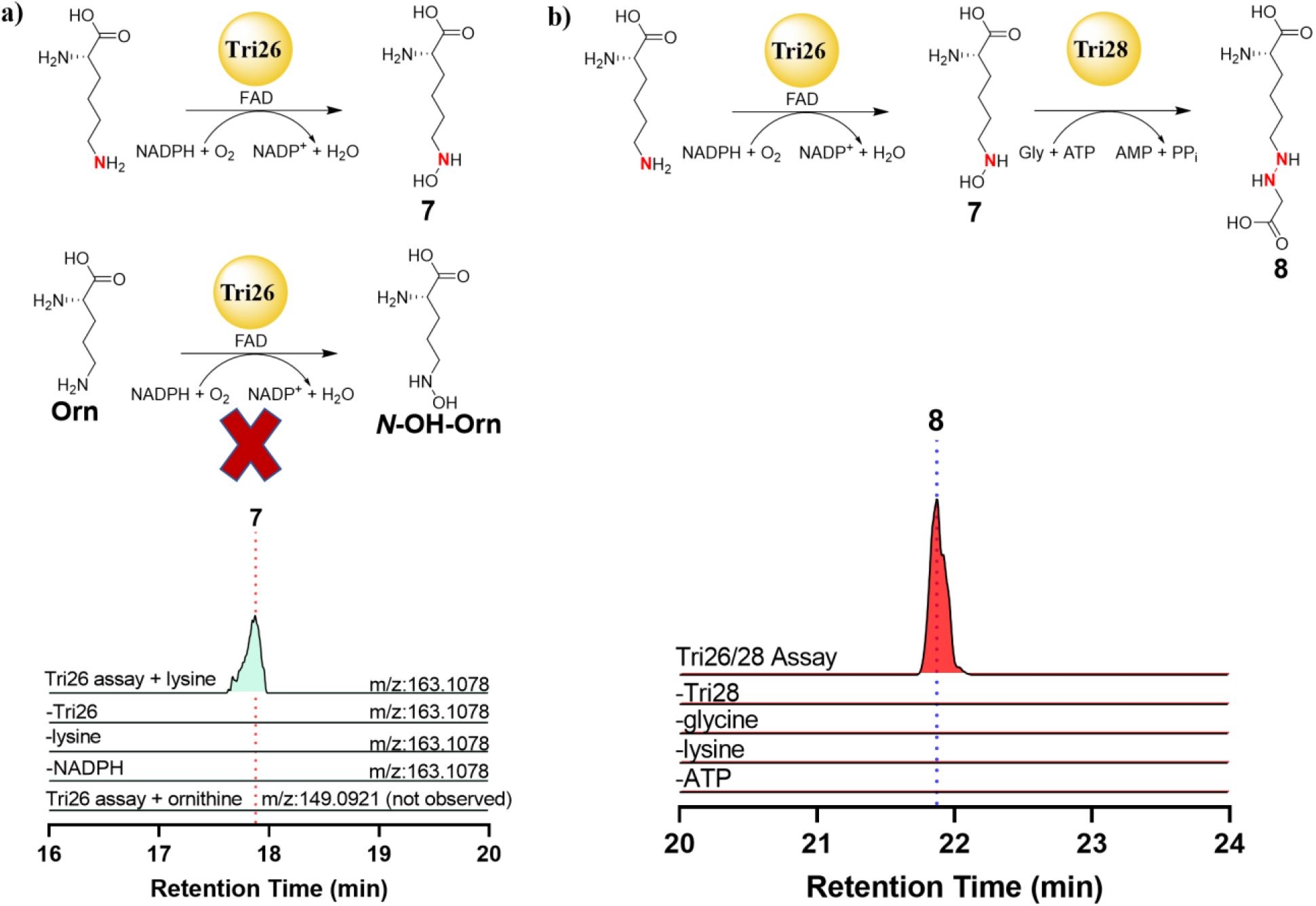
*In vitro* reconstitution of Tri26 and Tri28. a) EICs showing the production of **7** in an assay containing Tri26, lysine, and NADPH. Tri26 selectively hydroxylates lysine and not ornithine. The calculated mass for **7**: m/z=163.1078 ([M+H]^+^) is used for each trace, except for the ornithine assay in which the calculated mass for ***N*-OH-Orn** was used (m/z=149.0921 ([M+H]^+^)). b) EICs demonstrating that the production of **8** is dependent on Tri28, glycine, lysine, and ATP, along with the Tri26 assay components. The calculated mass for **8**: m/z=220.1292 ([M+H]^+^) is used for each trace. A 10-ppm mass error tolerance was used for each trace. At least three independent replicates were performed for each assay, and representative results are shown.

**Extended Data Figure 3.**
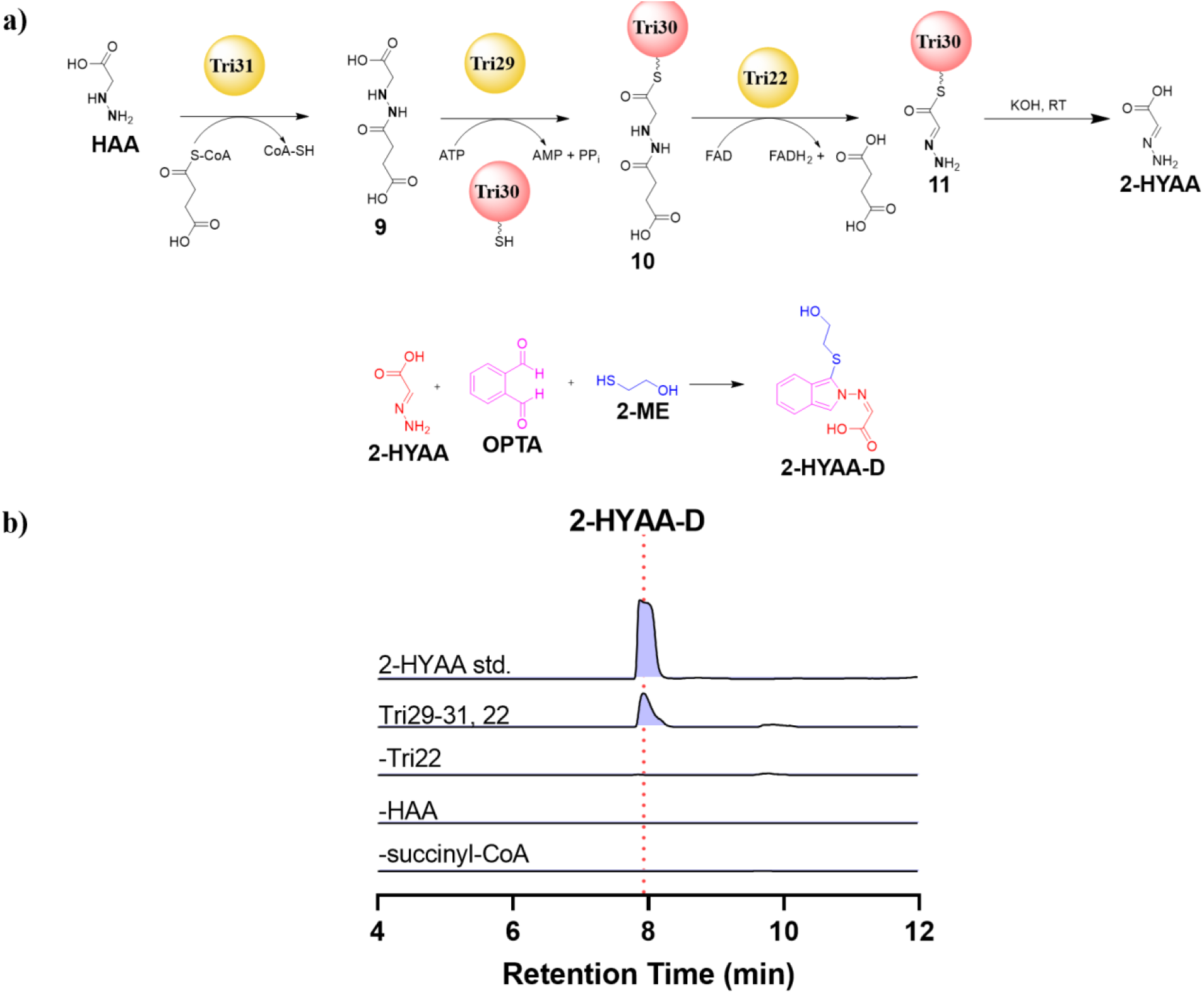
Characterization of the Tri22-catalyzed reaction product, 11. a) Schematic of a biochemical assay performed containing HAA, succinyl-CoA, Tri29-31, and Tri22 that is subjected to base hydrolysis, followed by OPTA derivatization to yield 2-HYAA-D. b) EICs showing the generation of 2-HYAA-D from the biochemical assay previously described in comparison with a derivatized 2-HYAA synthesized standard. Omission of Tri22, HAA, or succinyl-CoA from the couple enzymatic reaction resulted in abolishment of 2-HYAA-D. The calculated mass for 2-HYAA-D: m/z=265.0642 ([M+H]^+^) is used for each trace. A 10-ppm mass error tolerance was used for each trace. At least three independent replicates were performed for each assay, and representative results are shown.

**Extended Data Figure 4.**
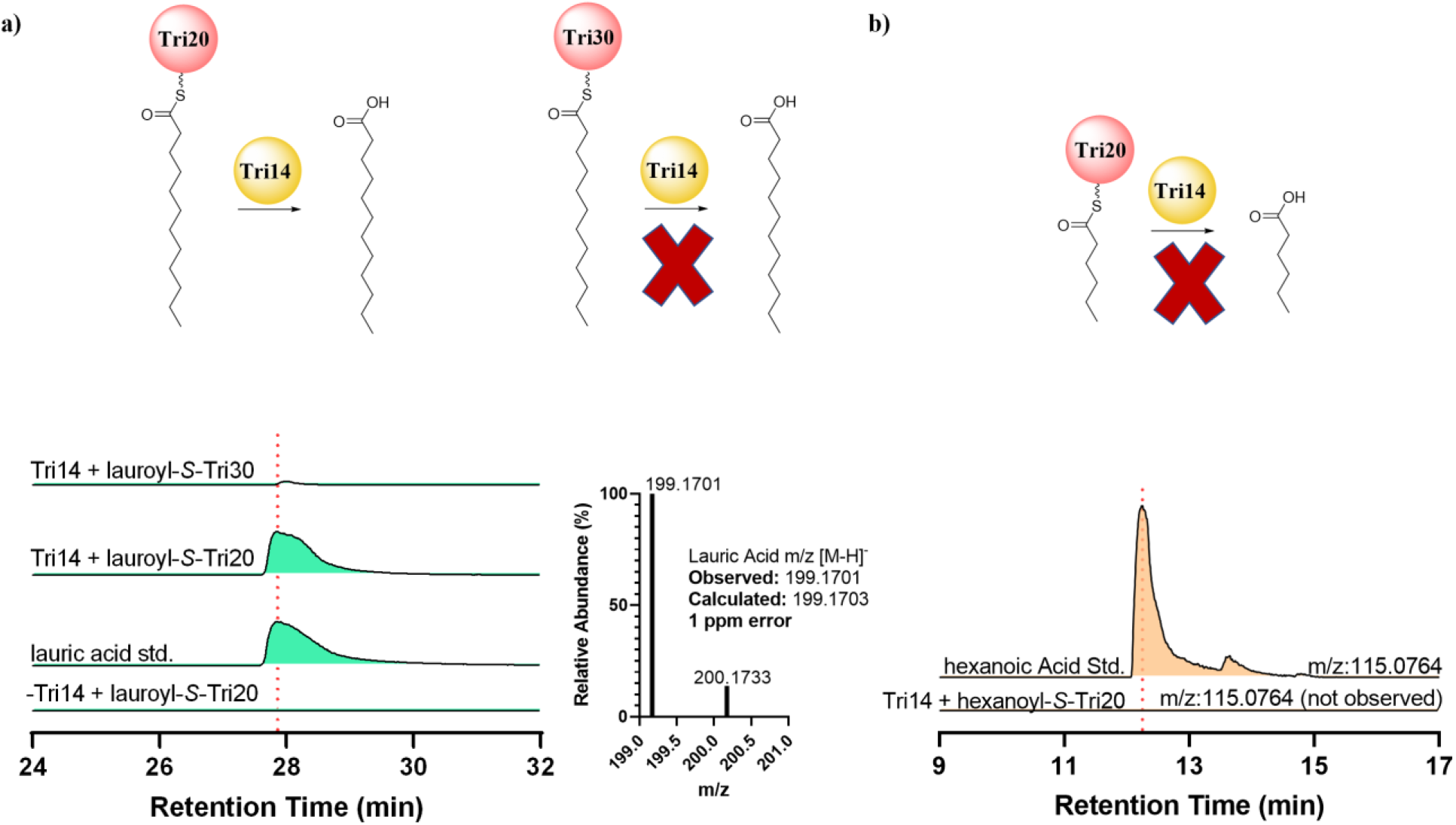
*In vitro* analysis of Tri14, a putative thioesterase. a) EICs demonstrating hydrolysis of lauroyl-*S*-Tri20 catalyzed by Tri14, resulting in production of lauric acid when compared to an authentic standard. Tri14 was shown to be Tri20 specific and imparted no activity on lauroyl-*S*-Tri30. The calculated mass for lauric acid: m/z=199.1701 ([M-H]^−^) is used for each trace. b) EIC demonstrating inability of Tri14 to hydrolyze hexanoyl-*S*-Tri20, thus suggesting that Tri14 activity is chain length specific. The calculated mass for hexanoic acid: m/z=115.0764 ([M-H]^−^) is used for each trace. A 10-ppm mass error tolerance was used for each trace. At least three independent replicates were performed for each assay, and representative results are shown.

**Extended Data Figure 5.**
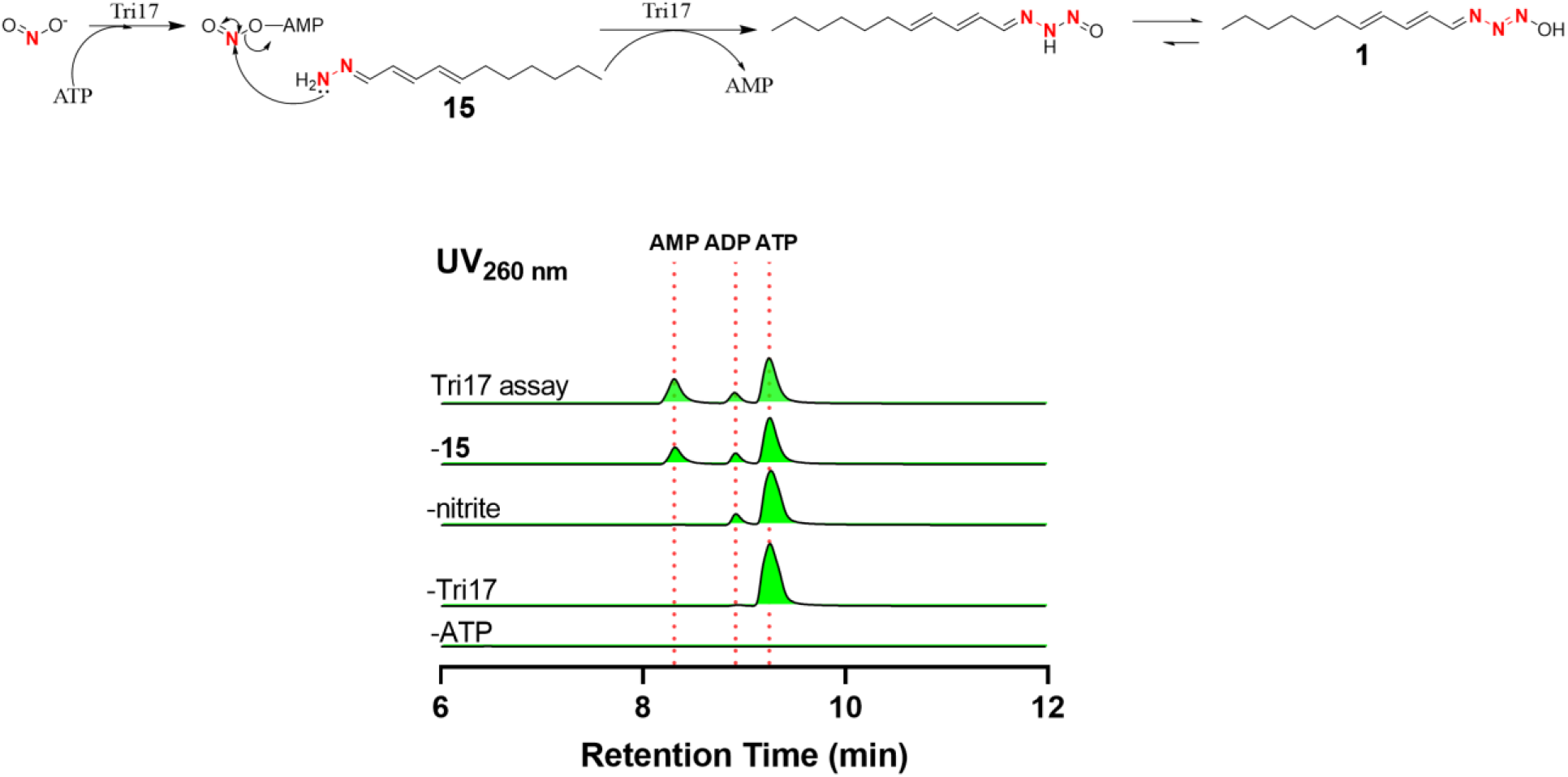
AMP Formation from Tri17 assay and proposed reaction mechanism. LC-UV detection of AMP at 260 nm from an assay containing **15**, nitrite, ATP, and Tri17. Controls lacking enzyme or nitrite result in undetectable amounts of AMP, while the production of AMP was slightly increased in the presence of **15**. ATP is proposed to activate nitrite to form a nitrite-AMP intermediate that undergoes a subsequent nucleophilic attack by **15** and tautomerization to yield **1**.

