## Supplementary Information for "Total Biosynthesis of Triacsin Featuring an *N*-hydroxytriazene Pharmacophore"

‡ Department of Chemical and Biomolecular Engineering, University of California Berkeley, Berkeley, California 94720 United States. [#]Department of Pharmaceutical Sciences, University of Shizuoka, Shizuoka 422-8526, Japan. [§]Chan Zuckerberg Biohub, San Francisco, California, 94158, United States. [♦] Department of Chemistry, University of California Berkeley, Berkeley, California 94720 United States. [✚] Department of Plant and Microbial Biology, University of California Berkeley, Berkeley, California 94720 United States.

[†]Co-First Authors

*

**Table of contents** **Pages**

**I. Supplementary Tables** 3-5

**Supplementary Table 1.** Oligonucleotides used in this study. 3-4

**Supplementary Table 2.** Plasmids used in this study. 5

**Supplementary Table 3.** Strains used in this study. 5

**II. Chemical Synthesis Methods** 6

Synthesis of 2-hydrazineylideneacetic acid. 6

Synthesis of ((2E,4E)-undeca-2,4-dien-1-ylidene)hydrazine. 6

**III. Supplementary Figures** 7-32

**Supplementary Figure 1.** Recently characterized N-N bond forming enzymes. 7

**Supplementary Figure 2.** UV time course of **5**. 8

**Supplementary Figure 3.** SDS-PAGE analysis of recombinant proteins from *E. coli*. 9

**Supplementary Figure 4.** HRMS and UV profile of **6**. 10
**Supplementary Figure 5.** HRMS of **7**, **8,** and HAA-DNP. 11-12
**Supplementary Figure 6. HAA** loading on Tri30. 13
**Supplementary Figure 7.** *In vitro* reconstitution of Tri31 and evidence of co-purified succinyl-CoA. 14-16
**Supplementary Figure 8.** Kinetic and HRMS analysis of acyl-CoA substrates from Tri31 assays. 17-18
**Supplementary Figure 9.** Activation and loading of acyl-HAAs on Tri30. 19-21
**Supplementary Figure 10.** Characterization of Tri22 as a FAD-containing enzyme. 22
**Supplementary Figure 11.** Tri22 *in vitro* biochemical assay with **9**. 23-24
**Supplementary Figure 12.** *In vitro* generation of nitrite by Tri16 and Tri21. *25*
**Supplementary Figure 13.** LC-UV analysis of AMP formation from Tri17 assay with nitrite and **11**. 26
**Supplementary Figure 14.** Acyl-*S*-Tri20 and acyl-*S*-Tri30 formation using Sfp. 27-29
**Supplementary Figure 15.** UV profile of **1**. 30
**Supplementary Figure 16.** Substrate specificity of Tri17. 31
**Supplementary Figure 17.** Proposed mechanism for N-N bond formation by Tri28. 32

**IV. Supplementary Notes** 33-61

**Supplementary Note 1.** Structural elucidation of purified **5**. 33-39
**Supplementary Note 2.** NMR characterization of Tri31 reaction**.** 40-45
**Supplementary Note 3.** Structural confirmation of synthetic 2-HYAA. 46-51
**Supplementary Note 4.** Structural confirmation of synthetic **15**. 52-56
**Supplementary Note 5.** Structural confirmation of purified triacsin A (**1**). 57-61

**References** 62

### I. Supplementary Tables

**Supplementary Table 1:** Oligonucleotides used in this study.

| Primer | Sequence (5' -> 3') | Description |
|---|---|---|
| tri10-Forward | attaagcttATGACACATCTGGC | Expression of Tri10 from S. *aureofaciens* |
| tri10-Reverse | aagcggccgcTCAGTCGTCCCAG | |
| tri9-Forward | aacatATGCCGACGGAACG | Expression of Tri9 from S. *aureofaciens* |
| tri9-Reverse | aactcgagTGAGGTCGCCCCTC | |
| tri16-Forward | ggttgggaattgcaaACGACCACCCCCGTAC | Expression of Tri16 from S. *aureofaciens* |
| tri16-Reverse | ggagatgggaagtcaTTATGGGGTTTTCTCCG | |
| tri21-Forward | ggttgggaattgcaaTCACAGCCGCCTCGGA | Expression of Tri21 from S. *aureofaciens* |
| tri21-Reverse | ggagatgggaagtcaTTACCGGCCGAGCACCG | |
| tri26-Forward | ggttgggaattgcaaATGGCGCATGAGCCCATA | Expression of Tri26 from S. *tsukubaensis* |
| tri26-Reverse | ggagatgggaagtcattACGATGCGCCTCCCG | |
| tri27-Forward | ggttgggaattgcaaGTGGCCGTGGTGGGC | Expression of Tri27 from S. tsukubaensis |
| tri27-Reverse | ggagatgggaagtcattATGATGGCCCCTGGTTCC | |
| tri28-Forward | aacatATGATCATCAGCCGT | Expression of Tri28 from S. *tsukubaensis* |
| tri28-Reverse | aactcgagTCGGTCACCCTCC | |
| tri28metRS-Forward | gaaggagatatacatATGGACCGGCCGGTGTTCG | Expression of Tri28 metRS domain from S. *tsukubaensis* |
| tri28metRS-Reverse | tgcggccgcaagcttTCGGTCACCCTCCCCGGG | |
| tri28Cupin-Forward | aacatATGATCATCAGCCGT | Expression of Tri28 Cupin domain from S. *tsukubaensis* |
| tri28Cupin-Reverse | aatctcgagCCCGAAGGCCGT | |
| tri29-Forward | ggttgggaattgcaaATGACGGAGGGAACCC | Expression of Tri29 from S. *tsukubaensis* |
| tri29-Reverse | ggagatgggaagtcattAACGGGCATGGATTCC | |
| tri30-Forward | aaggATCCATGCCCGTTGAC | Expression of Tri30 from S. *tsukubaensis* |

| | | |
|---|---|---|
| tri30-Reverse | aactcgagTGCCTGTGGATTCCC | |
| tri31-Forward | ggttgggaattgcaaATGAGCTGGGCGGAAC | Expression of Tri31 from S. *tsukubaensis* |
| tri31-Reverse | ggagatgggaagtcattAGTCCGCCGCGGT | |
| tri22-Forward | ggttgggaattgcaAGTGGAGACCATCACCA | Expression of Tri22 from S. *aureofaciens* |
| tri22-Reverse | ggagatgggaagtcattaTCATCCGGCGAGGG | |
| tri20-Forward | attcatATGTCCTCTGACCTGC | Expression of Tri20 from S. *tsukubaensis* |
| tri20-Reverse | aatctcgagTCCGGCCGAACG | |
| tri17-Forward | aacatATGATGATCAGTGAAGACC | Expression of Tri17 from S. *tsukubaensis* |
| tri17-Reverse | aactcgagGCAGCCGCCGATCA | |
| tri19-Forward | aacatATGCCACCGTCCGT | Expression of Tri19 from S. *tsukubaensis* |
| tri19-Reverse | aactcgagGAAGTCGAGTTCCACGTC | |
| tri13-Forward | gaaggagatatacatATGAGCACTTCCCCGGTGTC | Expression of Tri13 from S. *tsukubaensis* |
| tri13-Reverse | tgcggccgcaagcttGGGGCACTTGCGGAGCAGTA | |
| tri14-Forward | taacatatgTGCACGACCTACGCCC | Expression of Tri14 from S. *tsukubaensis* |
| tri14-Reverse | aactcgagCGACCCGGCCGTCC | |
| tri18-Forward | ggttgggaattgcaAATGAGCGTCGACTACG | Expression of Tri18 from S. *tsukubaensis* |
| tri18-Reverse | ggagatgggaagtcattaTCAGAGGACGGCCAG | |

**Supplementary Table 2.** Plasmids used in this study.

| Plasmid | Origin | Function |
|---|---|---|
| pCC2FOS-triBGC | pCC2FOS | Fosmid containing tri BGC from S*. aurofaciens* |
| pCC2FOS-tri9-tri10 | pCC2FOS | Gene disruption of tri9-10 in S*. aurofaciens* |
| pCR-Blunt-tri26-28 | pCR-Blunt II-TOPO | Plasmid containing tri26-28 from *S. tsukubaensis* |
| pCR-Blunt-tri29-32 | pCR-Blunt II-TOPO | Plasmid containing tri29-32 from *S. tsukubaensis* |
| pETDuet-1::tri10 | pCR-Blunt II-TOPO | Expression vector of tri10 from S*. aurofaciens* |
| pETDuet-1::tri9-10 | pCR-Blunt II-TOPO | Dual Expression vector of tri9-10 from S*. aurofaciens* |
| pLATE52::tri16 | pCR-Blunt II-TOPO | Expression vector of tri16 from S*. aurofaciens* |
| pLATE52::tri21 | pCR-Blunt II-TOPO | Expression vector of tri21 from S*. aurofaciens* |
| pLATE52::tri26 | pCR-Blunt II-TOPO | Expression vector of tri26 from *S. tsukubaensis* |
| pLATE52::tri29 | pCR-Blunt II-TOPO | Expression vector of tri29 from *S. tsukubaensis* |
| pLATE52::tri31 | pCR-Blunt II-TOPO | Expression vector of tri31 from *S. tsukubaensis* |
| pLATE52::tri22 | pCR-Blunt II-TOPO | Expression vector of tri22 from S*. aurofaciens* |
| pET-24b(+)::tri28 | pCR-Blunt II-TOPO | Expression vector of tri28 from *S. tsukubaensis* |
| pET-24b(+)::tri30 | pCR-Blunt II-TOPO | Expression vector of tri30 from *S. tsukubaensis* |

**Supplementary Table 3.** Strains used in this study.

| Strain | Description |
|---|---|
| *Streptomyces aureofaciens ATCC 31442* | Triacsin producer and genetic source for Tri9-10,16, 21, and 22 |
| *Streptomyces aureofaciens Δtri9-10 ATCC 31442* | Mutant strain accumulating carboxytriacsin |
| *Streptomyces tsukubaensis NRRL 18488* | Triacsin producer and genetic source for remaining proteins |
| *Escherichia coli XL1-Blue* | General cloning host |
| *Escherichia coli BL21 Star (DE3)* | Host for expression of recombinant enzymes |
| *Escherichia coli BAP1* | Host for expression of holo forms of acyl carrier proteins |

### II. Chemical Synthesis Methods

**Synthesis of 2-hydrazineylideneacetic acid.** Glyoxylic acid monohyrate (184 mg, 2 mmol) and hydrazine monohydrate (500 mg,10 mmol) were dissolved in 20 mL and 50 mL of methanol, respectively. Glyoxylic acid was added to the hydrazine monohydrate solution over the course of 1 hour under stirring conditions. Once the glyoxylic acid was transferred, the reaction was stirred and heated at 70 ºC for 4 hours. The reaction was then allowed to cool to room temperature and the methanol was further removed under vacuum to yield an oily, yellow liquid with solids. The oily liquid and solids were washed with isopropanol repeatedly, followed by removal of isopropanol under vacuum. The overall yield was 85%. Direct detection of this compound with LC-MS was unattainable since the m/z is less than 100, lower than the detection limit of our mass spectrometer. The m/z of the OPTA derivatized product: $[M+H]^+$= 265.0645 (observed), $[M+H]^+$= 265.0642 (calculated), 1.1 ppm error. $^1$H NMR (900 MHz, DMSO-$d_6$): $\delta_H$ 6.96 (s, 1H), 6.62 (s, 1H); $^{13}$C NMR ($^1$H-$^{13}$C HSQC and $^1$H-$^{13}$C HMBC at 900 MHz, DMSO-$d_6$): $\delta_C$ 129.1, 132.4, 168.1, 169.0.

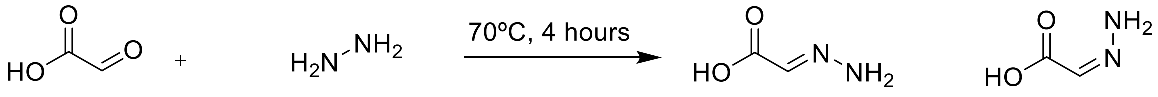


**glyoxylic acid** **hydrazine monohydrate** **2-HYAA**

**Synthesis of ((2*E*,4*E*)-undeca-2,4-dien-1-ylidene)hydrazine.** *2E,4E*-Undecadienal (332.5 mg, 2 mmol) and hydrazine monohydrate (500 mg,10 mmol) were dissolved in 20 mL and 50 mL of methanol, respectively. *2E,4E*-Undecadienal was added to the hydrazine monohydrate solution over the course of 1 hour under stirring conditions. Once the *2E,4E*-Undecadienal was transferred, the reaction was stirred and heated at 70 ºC for 4 hours. The reaction was then allowed to cool to room temperature and the methanol was further removed under vacuum to yield an oily, yellow liquid with solids. The oily liquid and solids were washed with isopropanol repeatedly, followed by removal of isopropanol under vacuum vacuum. The overall yield was 90%. The m/z of the compound: $[M+H]^+$= 181.1699 (observed), $[M+H]^+$= 181.1700 (calculated), 0.6 ppm error. $^1$H NMR (900 MHz, DMSO-$d_6$): $\delta_H$ 7.34 (d, $J$ = 9.4 Hz, 1H), 6.57 (s, 2H), 6.21 (dd, $J_1$ = 15.1 Hz, $J_2$ = 10.9 Hz, 1H), 6.11 (m, 2H), 5.72 (m 1H), 2.06 (dt, $J_1$ = 6.7 Hz, $J_2$ = 6.7 Hz, 2H), 1.35 (dd, $J_1$ = 7.4 Hz, $J_2$ = 6.9 Hz, 2H), 1.26 (m, 6H), 0.85 (t, $J$ = 7.0 Hz, 3H); $^{13}$C NMR (225 MHz, DMSO-$d_6$): $\delta_C$ 140.7, 134.9, 131.9, 130.4, 128.7, 32.2, 31.1, 28.7, 28.3, 22.1, 14.0.

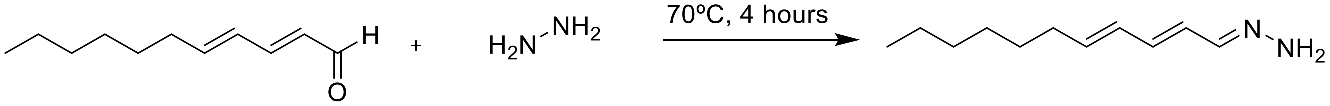


**2*E*,4*E*-undecadienal** **hydrazine monohydrate** **15**

### III. Supplementary Figures

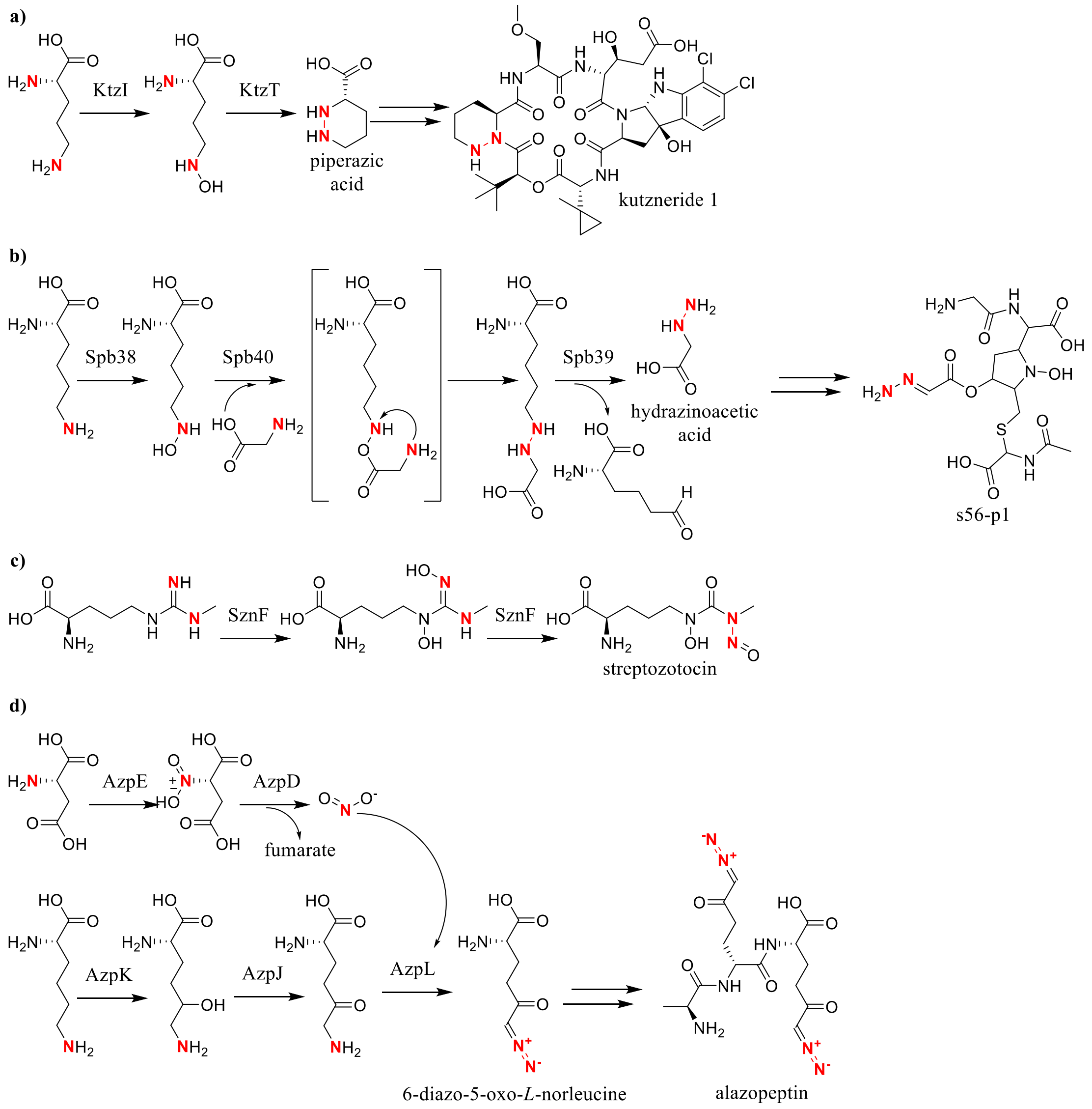


**Supplementary Figure 1.** Recently characterized N-N bond forming enzymes. a) The flavin-dependent KtzI promotes hydroxylation of the primary amine of the L-ornithine side chain, followed by the heme-dependent KtzT that catalyzes the cyclization of the amino acid to form piperazic acid. b) Spb38 firstly catalyzes hydroxylation of the primary amine of the L-lysine side chain. Spb40, a fusion protein consisting of a cupin and metRS domain, promotes the formation of $N^6$-(carboxymethylamino)lysine by utilizing $N^6$-hydroxylysine and glycine. Lastly, Spb39 lyses the product to yield a semialdehyde and hydrazinoacetic acid. c) The iron-dependent enzyme SznF utilizes molecular oxygen and NADPH to hydroxylate $N^{\omega}$-methylarginine twice before a subsequent intramolecular rearrangement. d) Nitrous acid generated by AzpE and AzpD is hypothesized to be incorporated into 5-oxolysine by the transmembrane protein, AzpL.

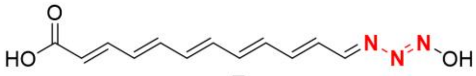


**5**

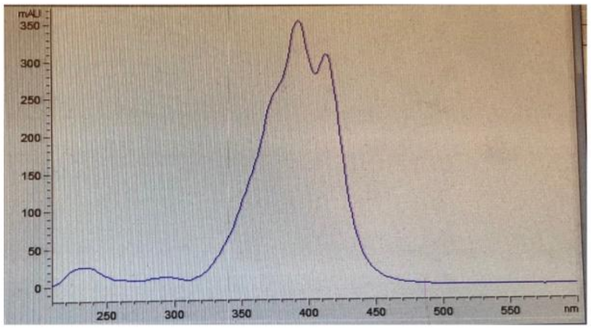


Purified **5**

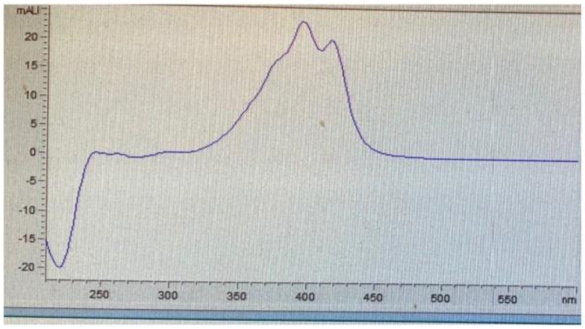


Purified **5** after two days at room temperature

**Supplementary Figure 2.** UV time course of **5**. UV profiles of freshly purified **5** and its degradation after 48 hours at room temperature.

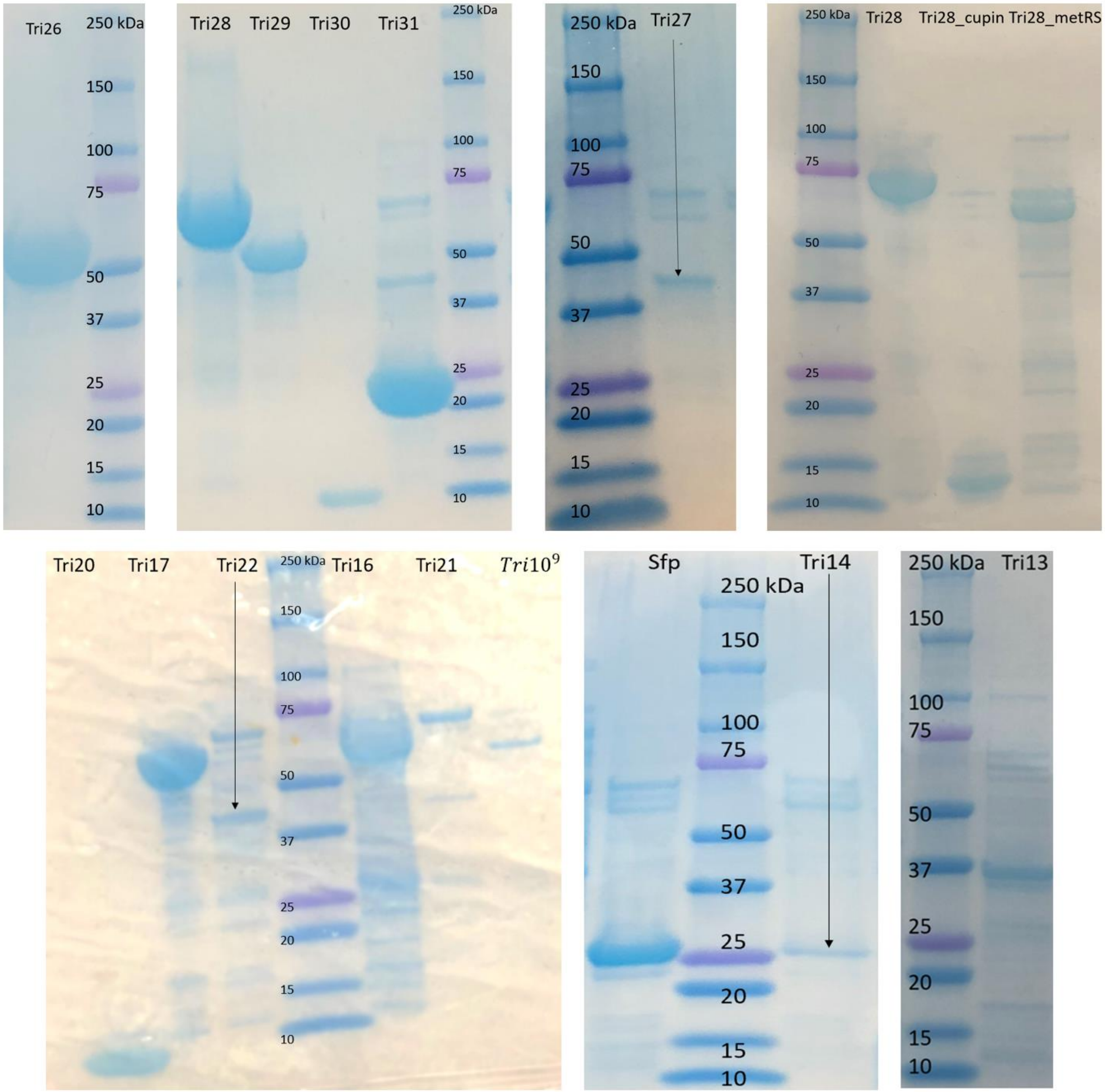


**Supplementary Figure 3.** SDS-PAGE analysis of recombinant proteins from *E. coli*. BioRad Mini-PROTEAN TGX gels (4-15% precast, 12 wells) were used to visualize proteins. The approximate molecular weight and yield for each protein are the following: Tri26 (52.4 kDa, 5.2 mg/L), Tri27 (40.9 kDa, 5.4 mg/L), Tri28 (74.7 kDa, 18.5 mg/L), Tri28_metRS (62.3 kDa, 6.8 mg/L), Tri28_cupin (14.1 kDa, 22.7 mg/L), Tri29 (57.5 kDa, 34.3 mg/L), Tri30 (11 kDa, 16.9 mg/L), Tri20 (9.8 kDa, 10.8 mg/L), Tri13 (34.5 kDa, 6 mg/L), Tri31 (25.1 kDa, 20.5 mg/L), Tri22 (45.9 kDa, 6.7 mg/L), Tri21 (71.8 kDa, 6.4 mg/L), Tri16 (59.6 kDa, 15.2 mg/L), Tri17 (61.5 kDa, 34 mg/L), Tri10[9] (56.7 kDa, 4.6 mg/L), and Tri14 (27.7 kDa, 5 mg/L).

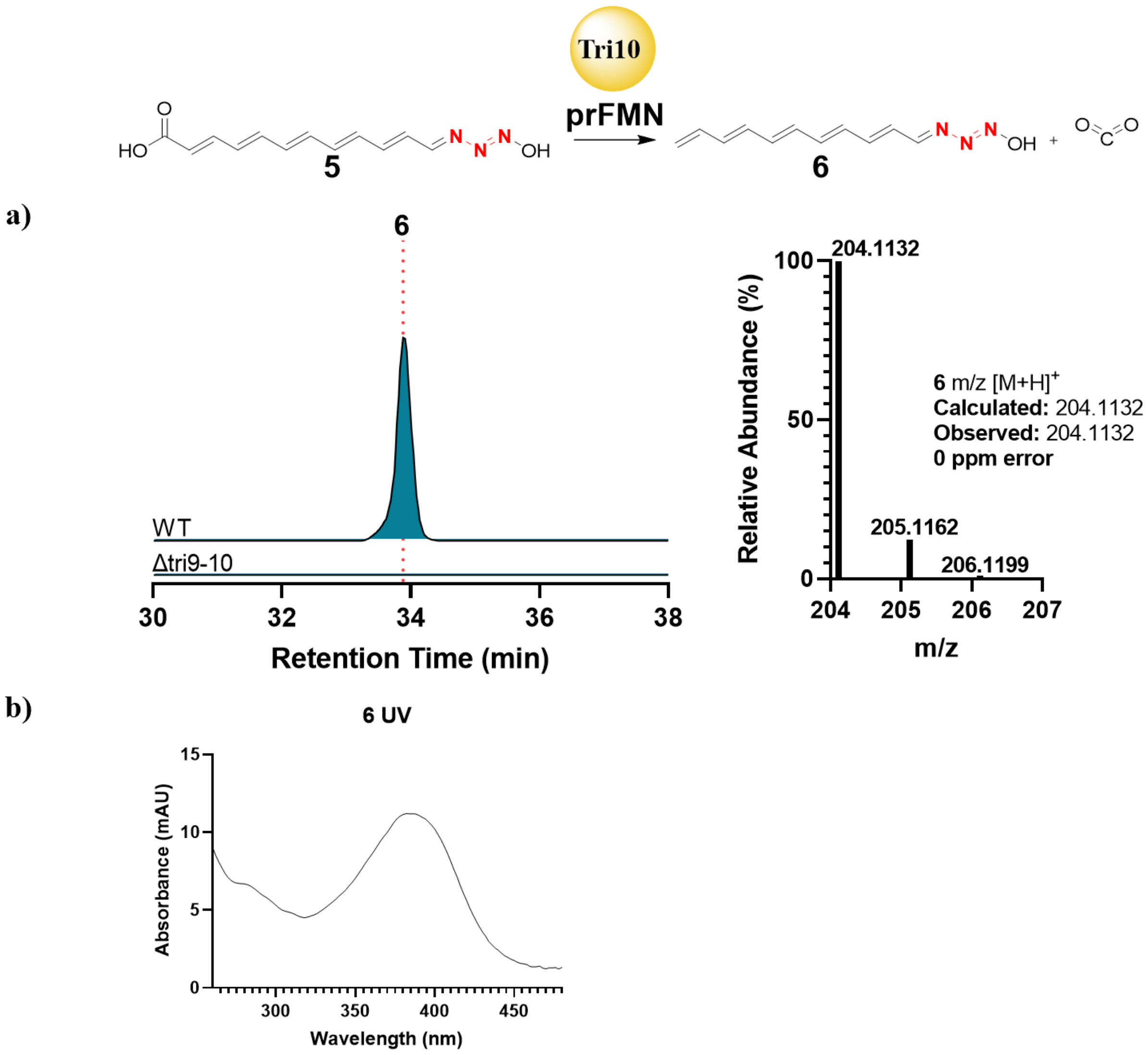


**Supplementary Figure 4.** HRMS and UV profile of **6**. a) HRMS of **6** detected in wild type *S. aureofaciens* and absent in the *Δtri9-10* mutant. b) UV profile of **6** detected in the wild type *S. aureofaciens*.

a)

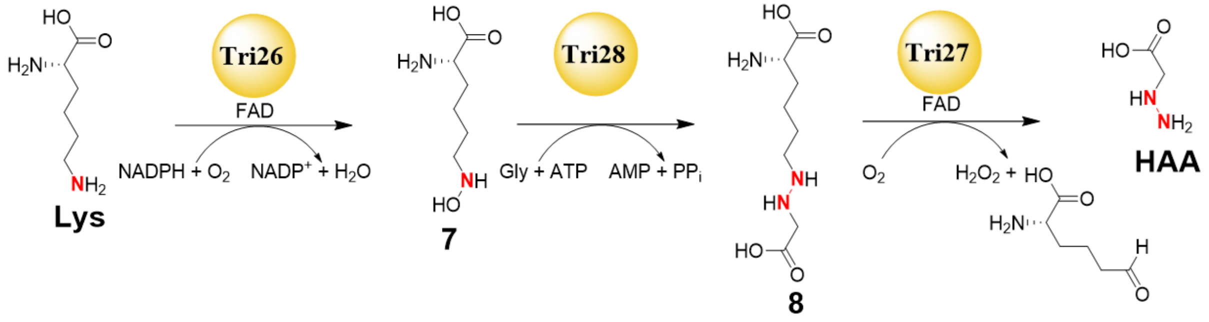
Tri26
FAD
NADPH + $O_2$
$NADP^+$ + $H_2O$
Tri28
Gly + ATP
AMP + $PP_i$
Tri27
FAD
$O_2$
$H_2O_2$ +
Lys
7
8
HAA

b)

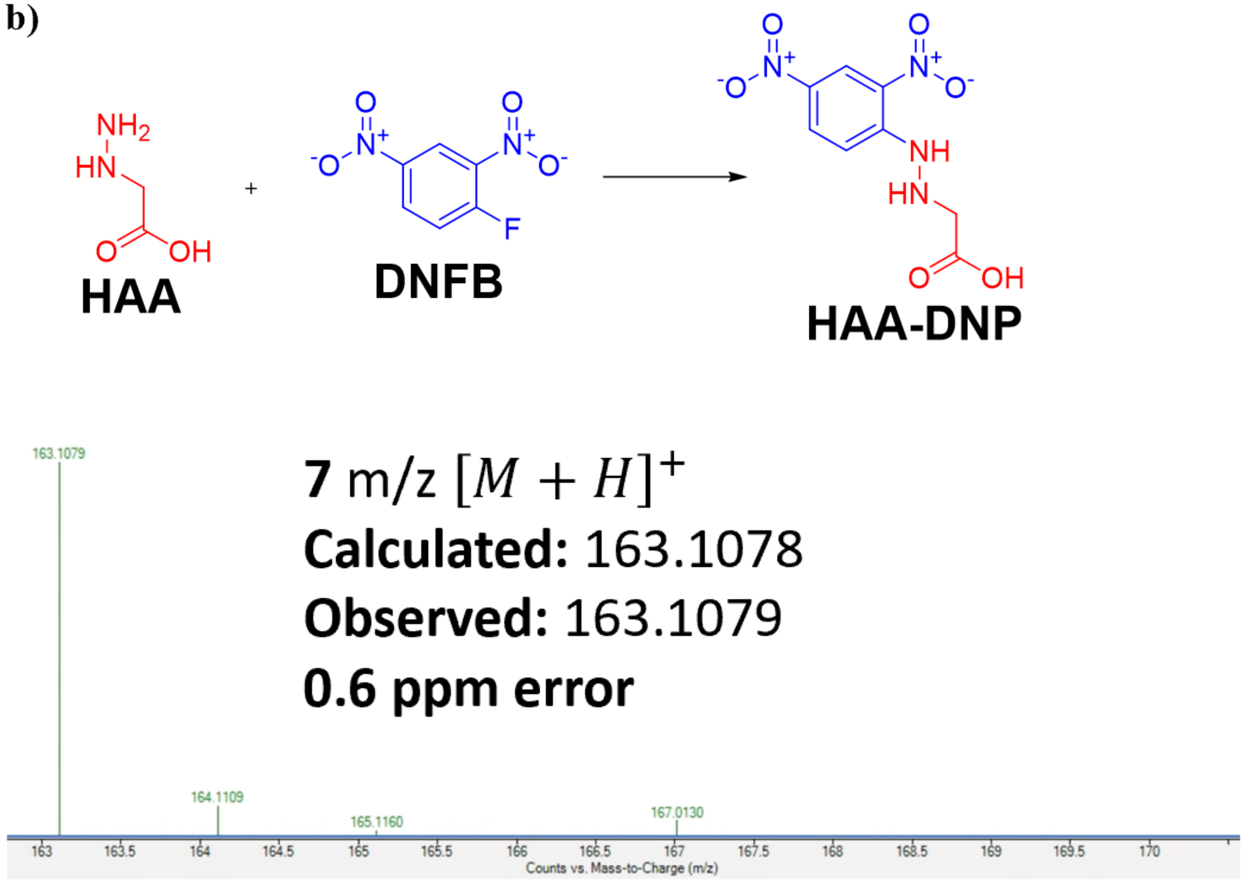
HAA
DNFB
HAA-DNP
7 m/z $[M + H]^+$
Calculated: 163.1078
Observed: 163.1079
0.6 ppm error
163.1079
164.1109
165.1160
167.0130
Counts vs. Mass-to-Charge (m/z)

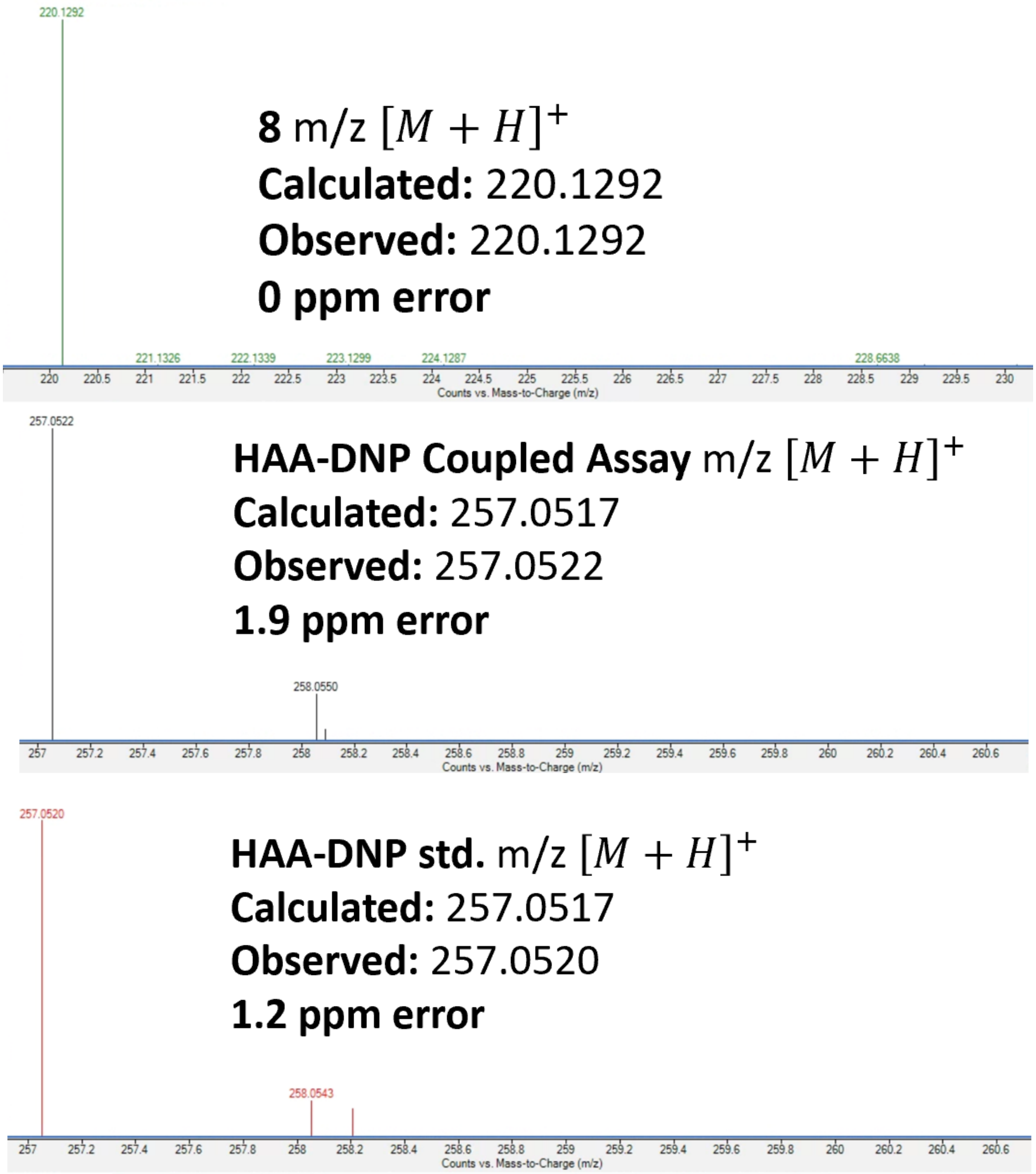


**Supplementary Figure 5.** HRMS of **7**, **8,** and HAA-DNP. a) HRMS of **7** and **8** afforded from *in vitro* reconstitution of Tri26 and Tri28. b) HRMS of HAA-DNP afforded from DNFB derivatization of a coupled Tri26-28 assay and an authentic standard, respectively.

**a)**

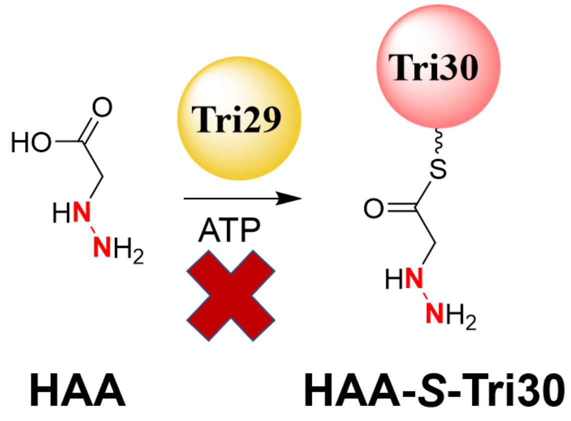


**b)**

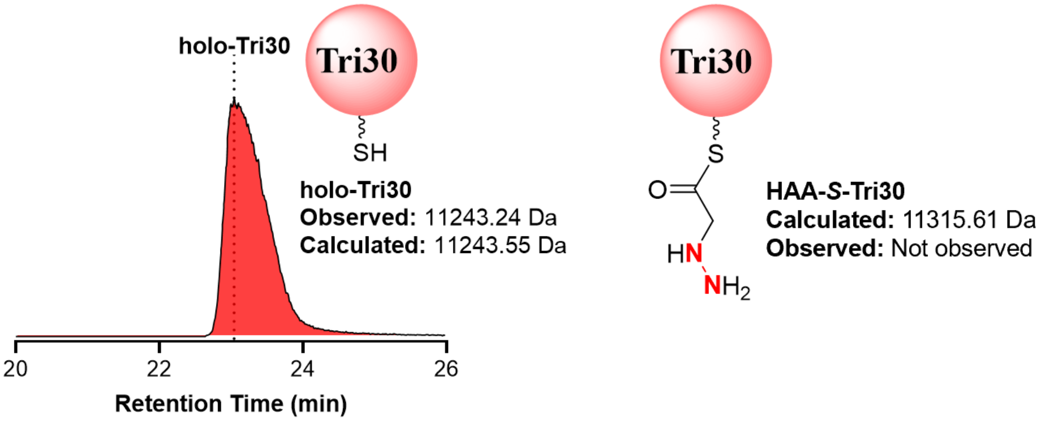


**Supplementary Figure 6.** HAA loading on Tri30. a) Schematic of attempted loading of HAA to Tri30 by Tri29. b) EIC of +12 charge state of holo-Tri30 showing failed conversion of holo-Tri30 to HAA-*S*-Tri30. The calculated intact protein mass for HAA-*S*-Tri30 was not observed.

a)

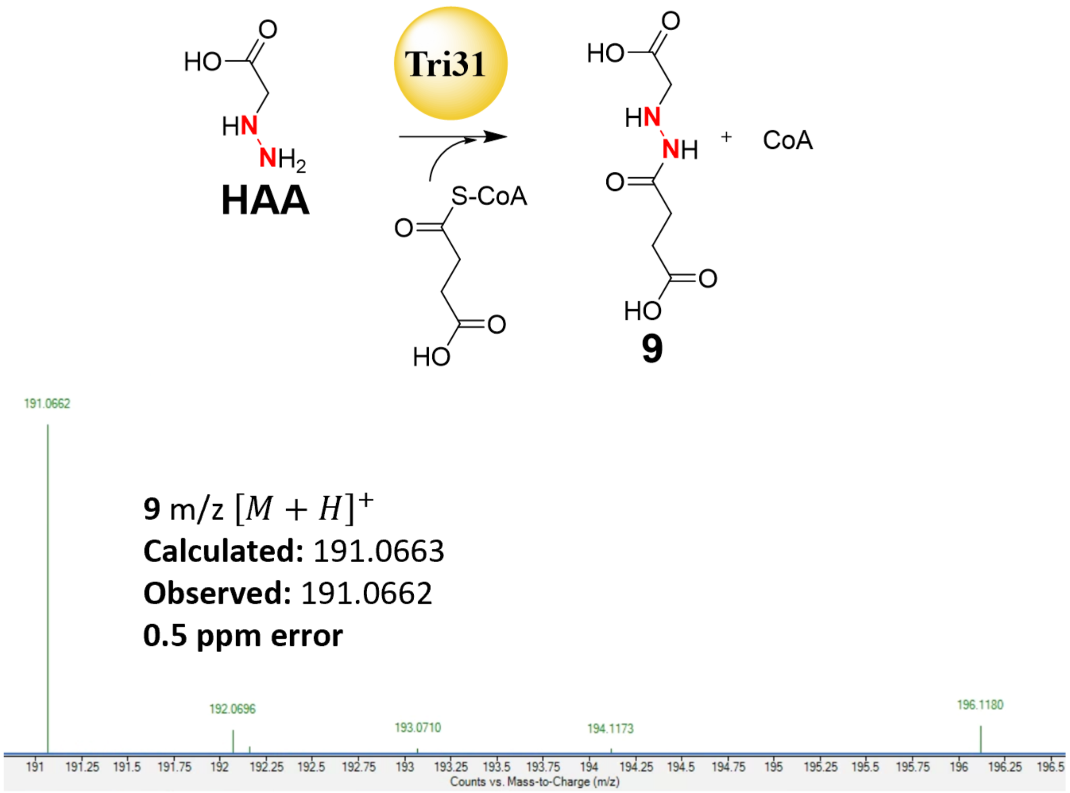

HO
O
HN
NH2
HAA
Tri31
S-CoA
O
HO
O
HO
O
HN
NH
O
HO
O
+ CoA
9
191.0662
9 m/z [M + H]+
Calculated: 191.0663
Observed: 191.0662
0.5 ppm error
192.0696
193.0710
194.1173
196.1180
191 191.25 191.5 191.75 192 192.25 192.5 192.75 193 193.25 193.5 193.75 194 194.25 194.5 194.75 195 195.25 195.5 195.75 196 196.25 196.5
Counts vs. Mass-to-Charge (m/z)


b)

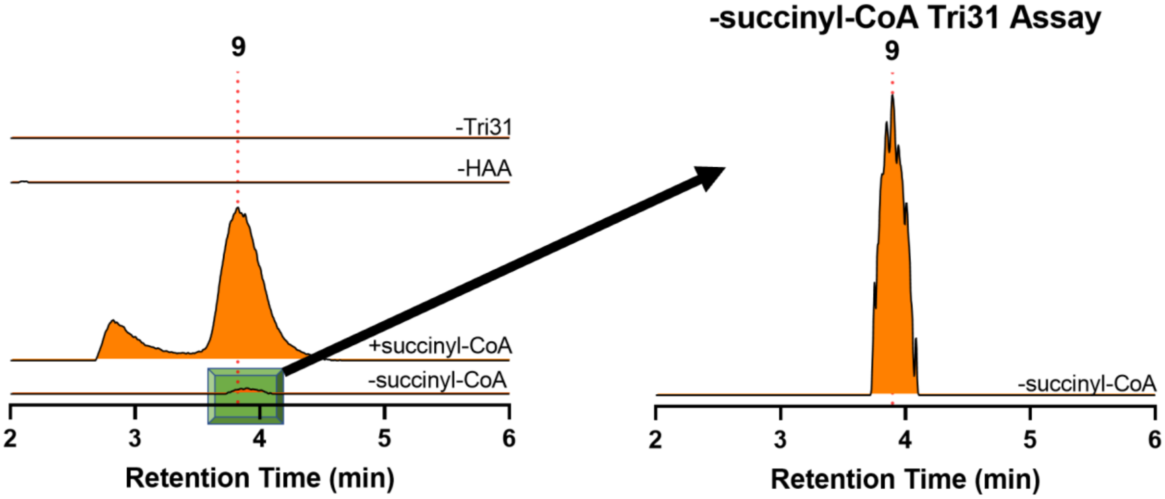

9
-Tri31
-HAA
+succinyl-CoA
-succinyl-CoA
2 3 4 5 6
Retention Time (min)
-succinyl-CoA Tri31 Assay
9
-succinyl-CoA
2 3 4 5 6
Retention Time (min)

c)

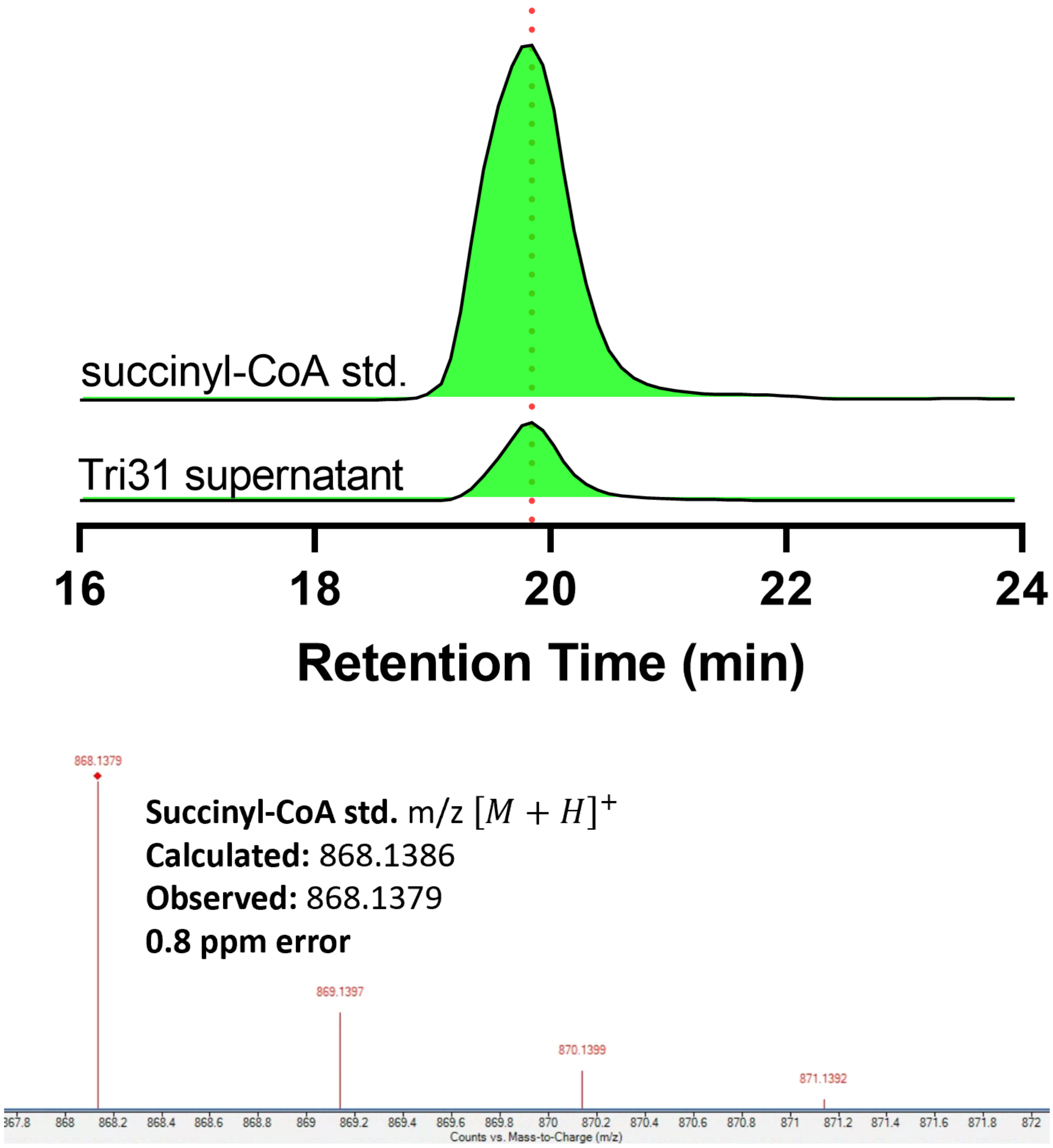
succinyl-CoA std.
Tri31 supernatant
16
18
20
22
24
Retention Time (min)
868.1379
Succinyl-CoA std. m/z [M + H]+
Calculated: 868.1386
Observed: 868.1379
0.8 ppm error
869.1397
870.1399
871.1392
Counts vs. Mass-to-Charge (m/z)

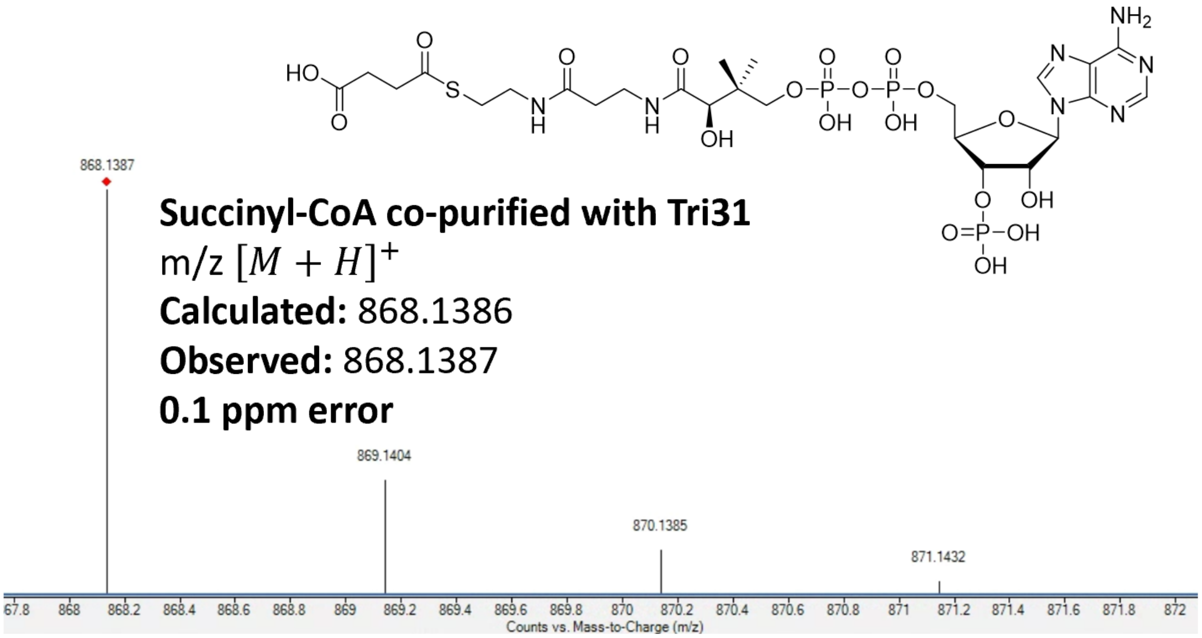


**Supplementary Figure 7.** *In vitro* reconstitution of Tri31 and evidence of co-purified succinyl-CoA. a) Schematic of Tri31 reaction with HRMS of a product consistent with succinylation of HAA (**9**). b) LC-HRMS chromatograms showing that trace amounts of **9** are detected without adding succinyl-CoA and subsequent addition leads to higher yield of **9**. c) LC-HRMS chromatograms showing trace amounts of succinyl-CoA present in purified Tri31 in comparison with an authentic standard.

**a)**

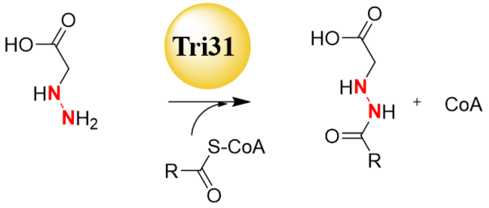


| Substrate | $\frac{k_{cat}}{K_m}$ $(mM^{-1}min^{-1})$ |
|---|---|
| succinyl-CoA | 14.7 ± 2.1 |
| hexanoyl-CoA | 5.4 ± 0.1 |
| lauroyl-CoA | 4.2 ± 0.9 |
| malonyl-CoA | 2.4 ± 0.3 |
| crotonoyl-CoA | 1.5 ± 0.1 |

**b)**

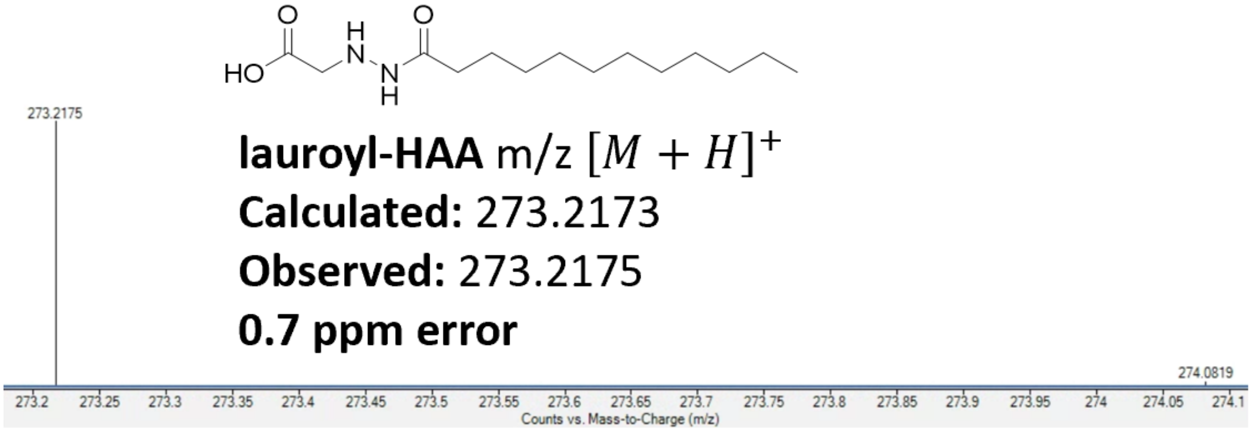


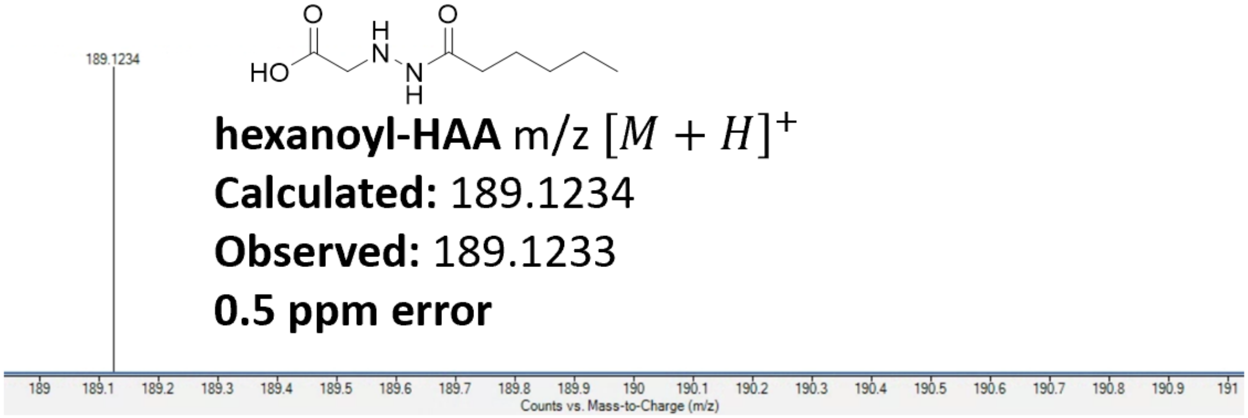

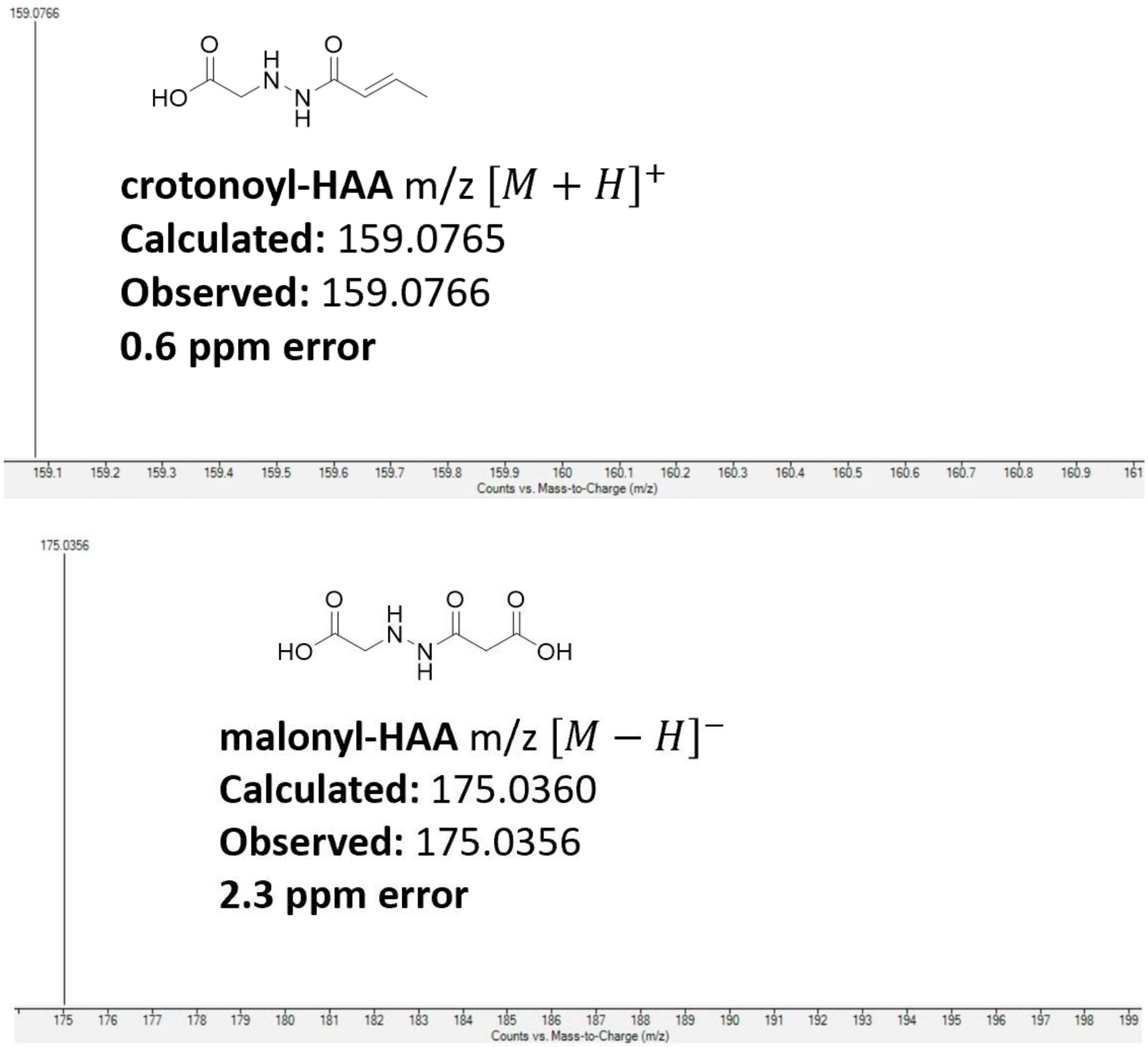


**Supplementary Figure 8.** Kinetic and HRMS analysis of acyl-CoA substrates from Tri31 assays. a) Schematic of Tri31 reaction between acyl-CoA substrates and **HAA** yielding acyl-HAA products. The $\frac{k_{cat}}{K_M}$ parameters demonstrate that Tri31 moderately prefers succinyl-CoA over the other substrates. Kinetic assays were performed in triplicate. The values correspond to the average and standard deviation from the triplicate trials, respectively. b) HRMS of acyl-HAA products.

a)

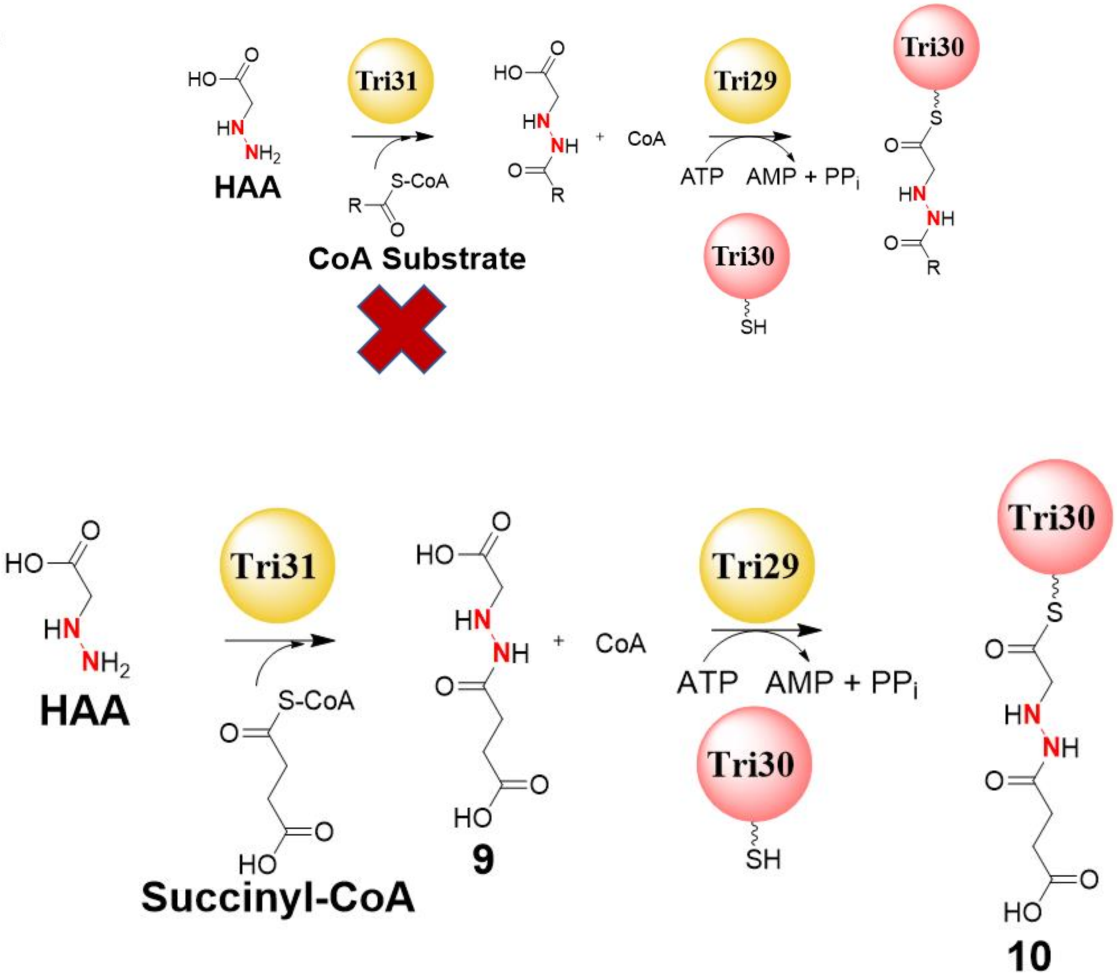

HAA
Tri31
CoA Substrate
+ CoA
Tri29
ATP AMP + PP$_i$
Tri30
HAA
Tri31
Succinyl-CoA
9
+ CoA
Tri29
ATP AMP + PP$_i$
Tri30
10

**b)**

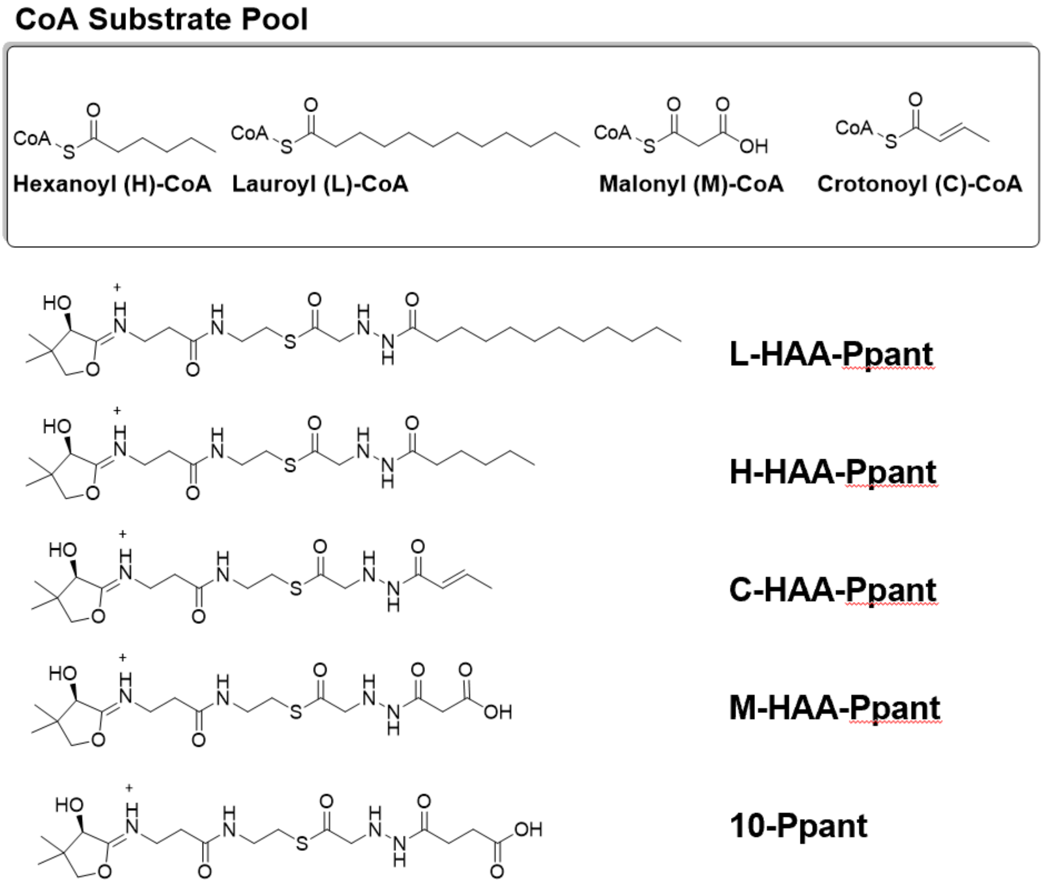
CoA Substrate Pool
Hexanoyl (H)-CoA
Lauroyl (L)-CoA
Malonyl (M)-CoA
Crotonoyl (C)-CoA
L-HAA-Ppant
H-HAA-Ppant
C-HAA-Ppant
M-HAA-Ppant
10-Ppant

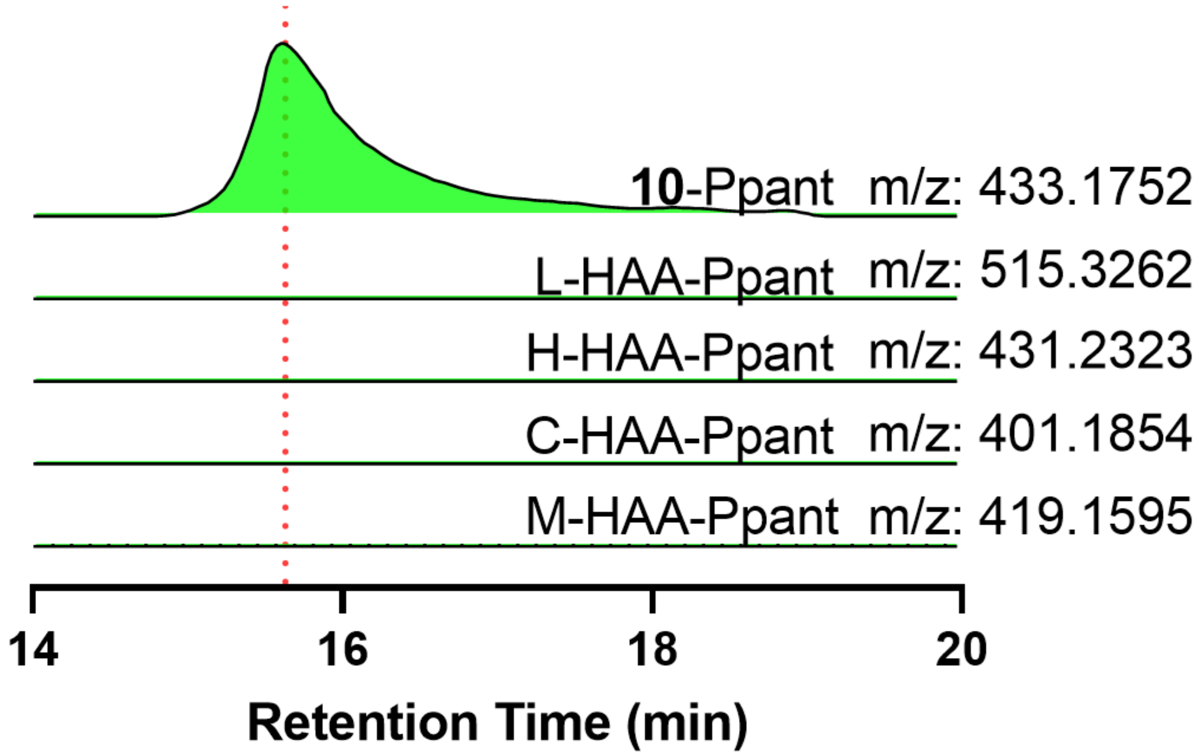


**Supplementary Figure 9.** Activation and loading of acyl-HAAs on Tri30. a) Schematic showing a coupled Tri29-31 assay consisting of acylation of HAA, activation, and loading on Tri30. b) LC-HRMS chromatograms showing that only succinyl-HAA was successfully activated and loaded on Tri30 to yield **10**. The Ppant fragments from other acyl-HAAs were not detected.

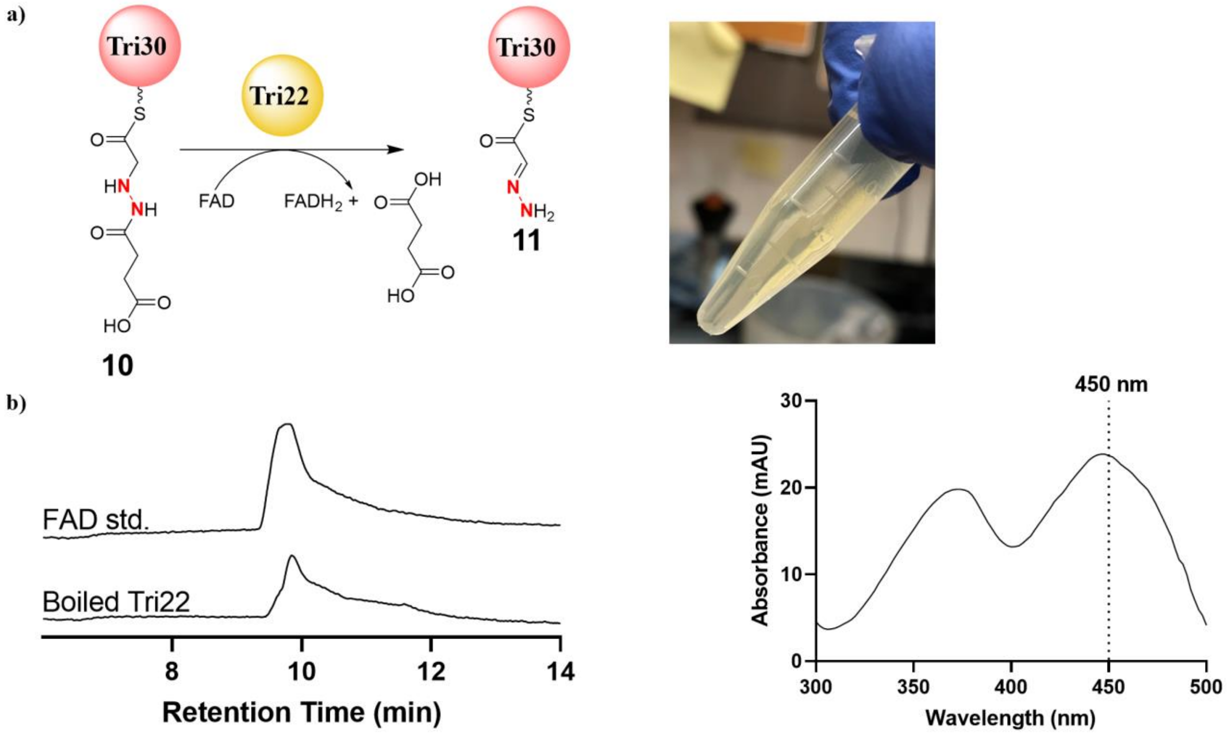


**Supplementary Figure 10.** Characterization of Tri22 as a FAD-containing enzyme. a) Schematic of the Tri22 catalyzed reaction and a picture of the yellow protein purified from *E. coli*. b) LC-HRMS and UV analysis of boiled Tri22 in comparison with an authentic FAD standard. The peak at 450 nm is indicative of a trace amount of bound-FAD. Photo by A. Del Rio Flores.

a)

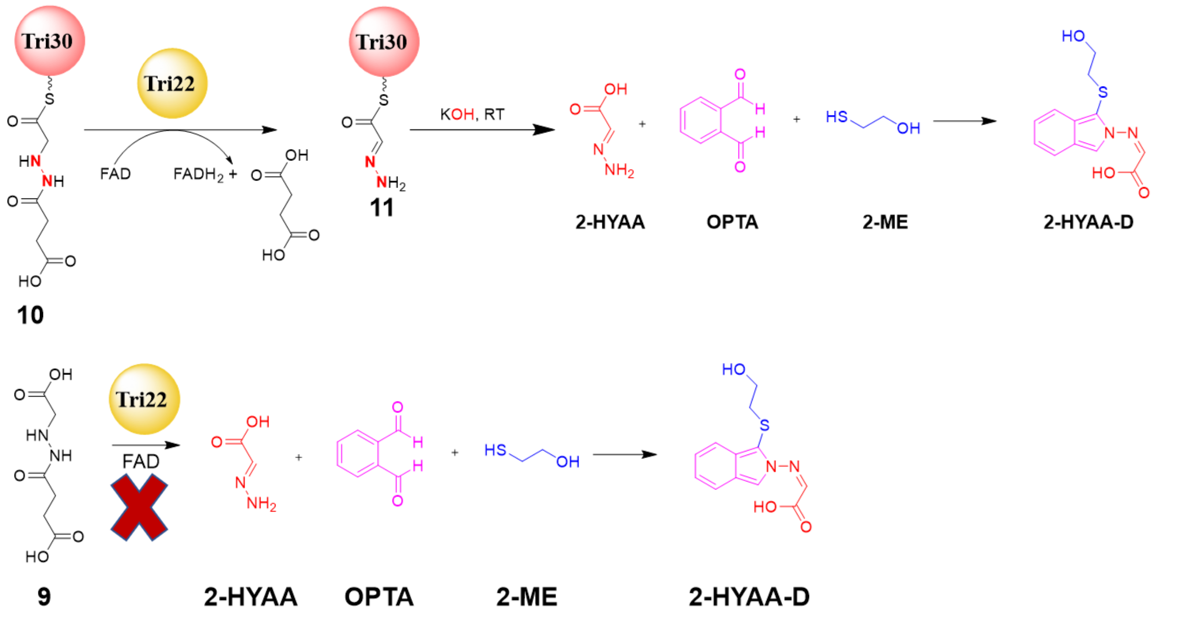
Tri30
Tri22
FAD
FADH2 +
KOH, RT
10
11
2-HYAA
OPTA
2-ME
2-HYAA-D
9
2-HYAA
OPTA
2-ME
2-HYAA-D

b)

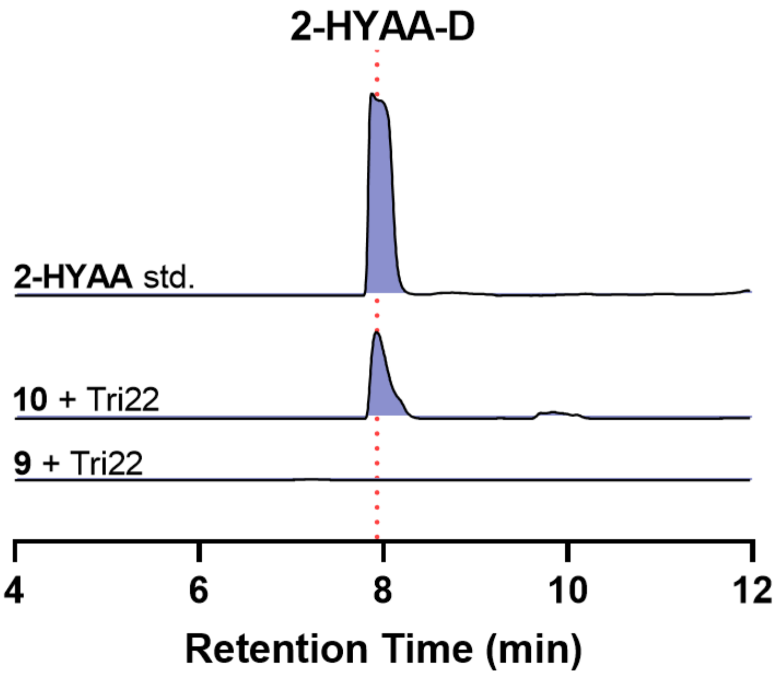
2-HYAA-D
2-HYAA std.
10 + Tri22
9 + Tri22
4
6
8
10
12
Retention Time (min)

**c)**

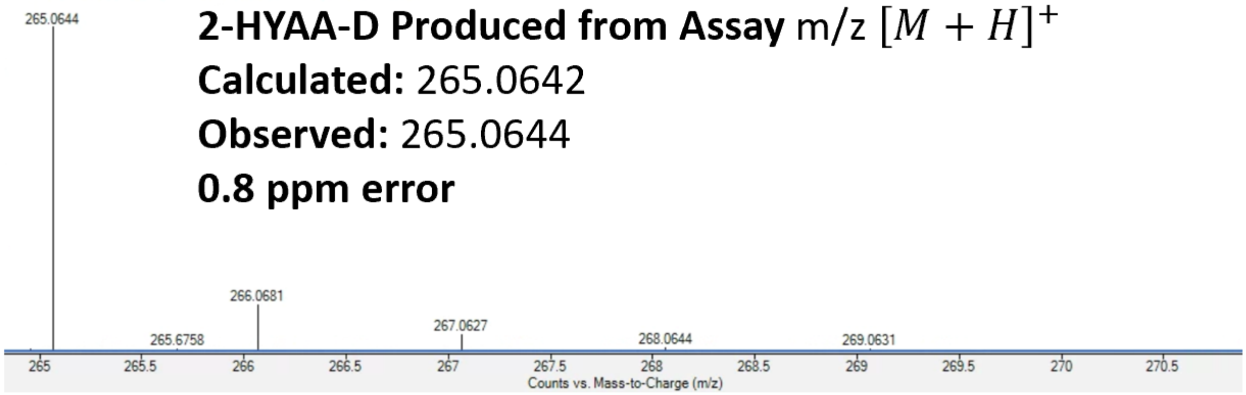


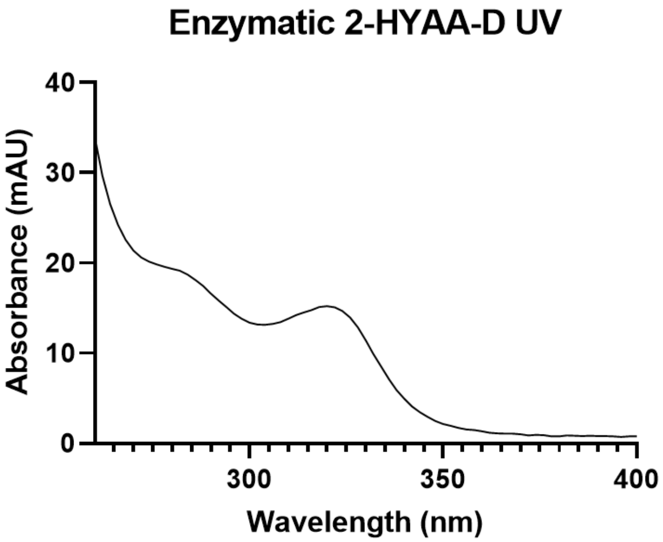


**Supplementary Figure 11.** Tri22 *in vitro* biochemical assay with **9**. a) Schematics of attempted Tri22 assays using **10** or **9**, followed by OPTA derivatization. b) EICs demonstrating production of 2-HYAA-D. c) HRMS and UV of 2-HYAA-D generated from the assay with **10** and OPTA derivatization.

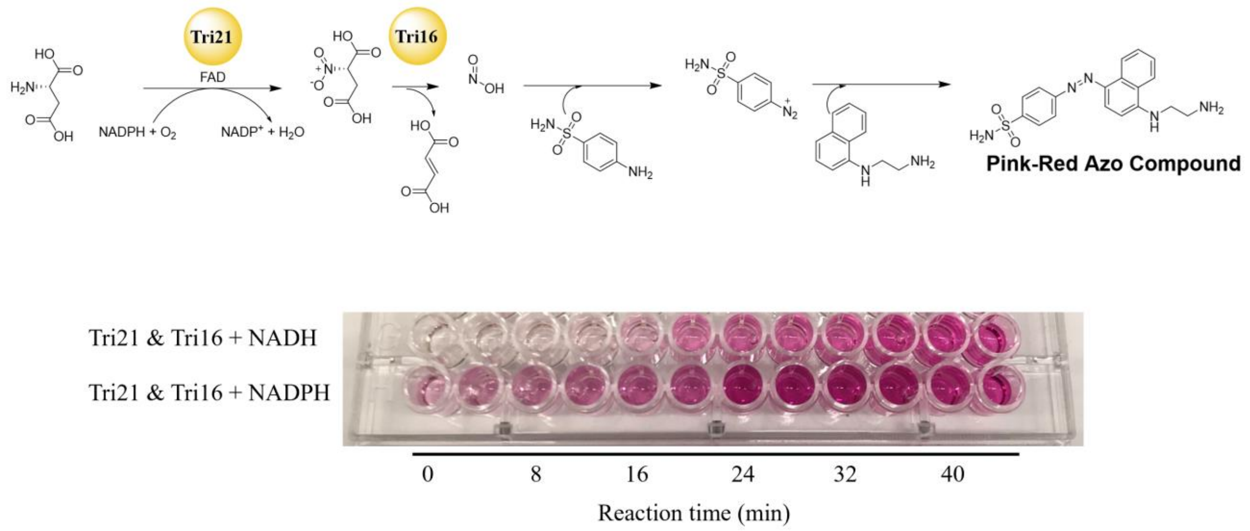


**Supplementary Figure 12.** *In vitro* generation of nitrite by Tri16 and Tri21. Nitrite produced by Tri16 and Tri21 was detected by utilizing the Griess test, a colorimetric assay generating a pink-red diazo compound with a unique absorbance at 550 nm. Negative controls lacking Tri21, Tri16, NADPH/NADH, or aspartate resulted in no color change. Based on the relative intensity from the time course, NADPH appears to be the preferred substrate for Tri21.

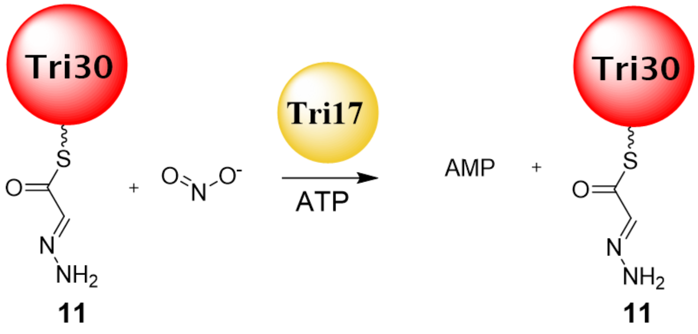


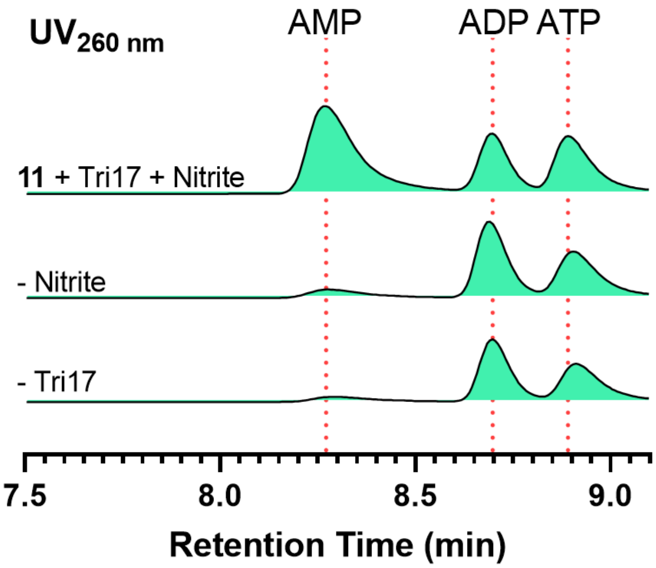


**Supplementary Figure 13.** LC-UV analysis of AMP formation from Tri17 assay with nitrite and **11**. **11** was generated in situ using Tri29-31. Tri22. While **11** was not modified in the Tri17 assay, AMP was demonstrated to be generated which was dependent on both nitrite and Tri17.

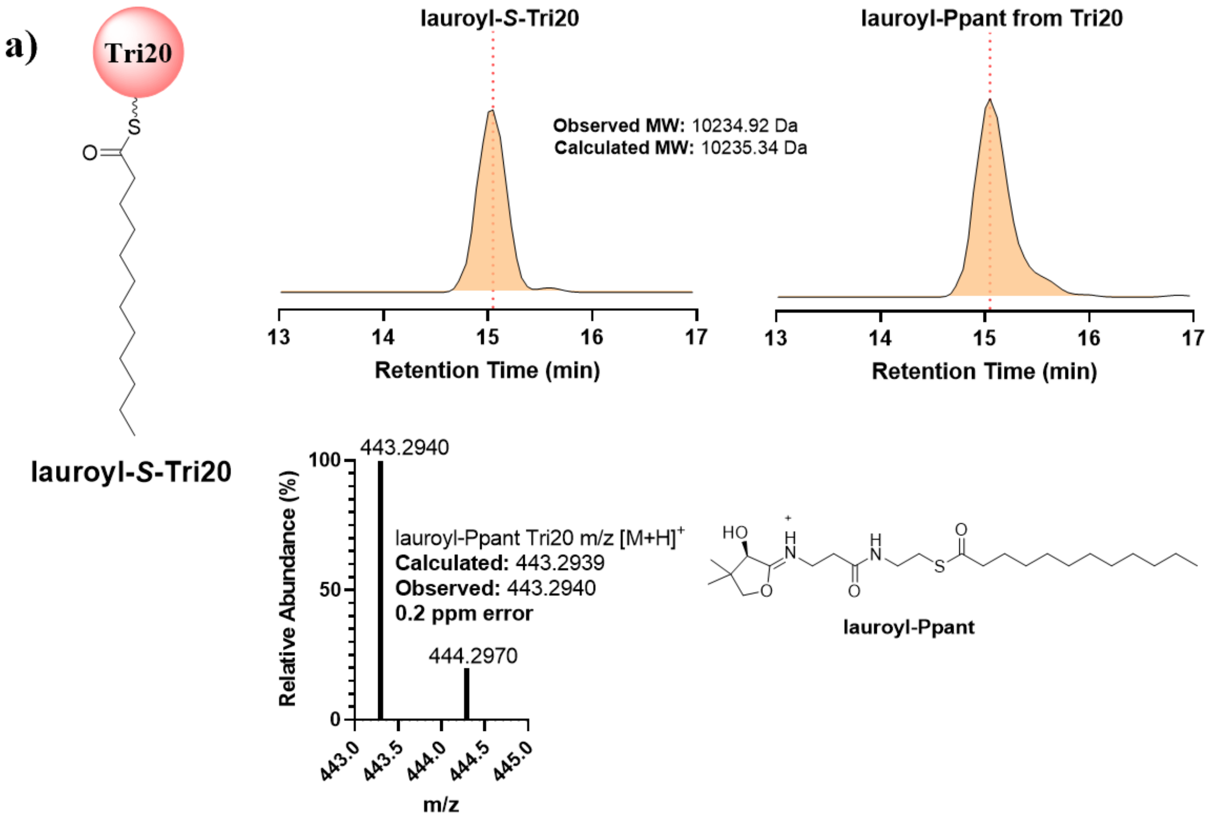
a)
Tri20
S
O
lauroyl-S-Tri20
lauroyl-S-Tri20
Observed MW: 10234.92 Da
Calculated MW: 10235.34 Da
13
14
15
16
17
Retention Time (min)
lauroyl-Ppant from Tri20
13
14
15
16
17
Retention Time (min)
443.2940
100
50
0
Relative Abundance (%)
lauroyl-Ppant Tri20 m/z [M+H]+
Calculated: 443.2939
Observed: 443.2940
0.2 ppm error
444.2970
443.0
443.5
444.0
444.5
445.0
m/z
HO
H
N
H
N
O
O
S
O
lauroyl-Ppant

b)

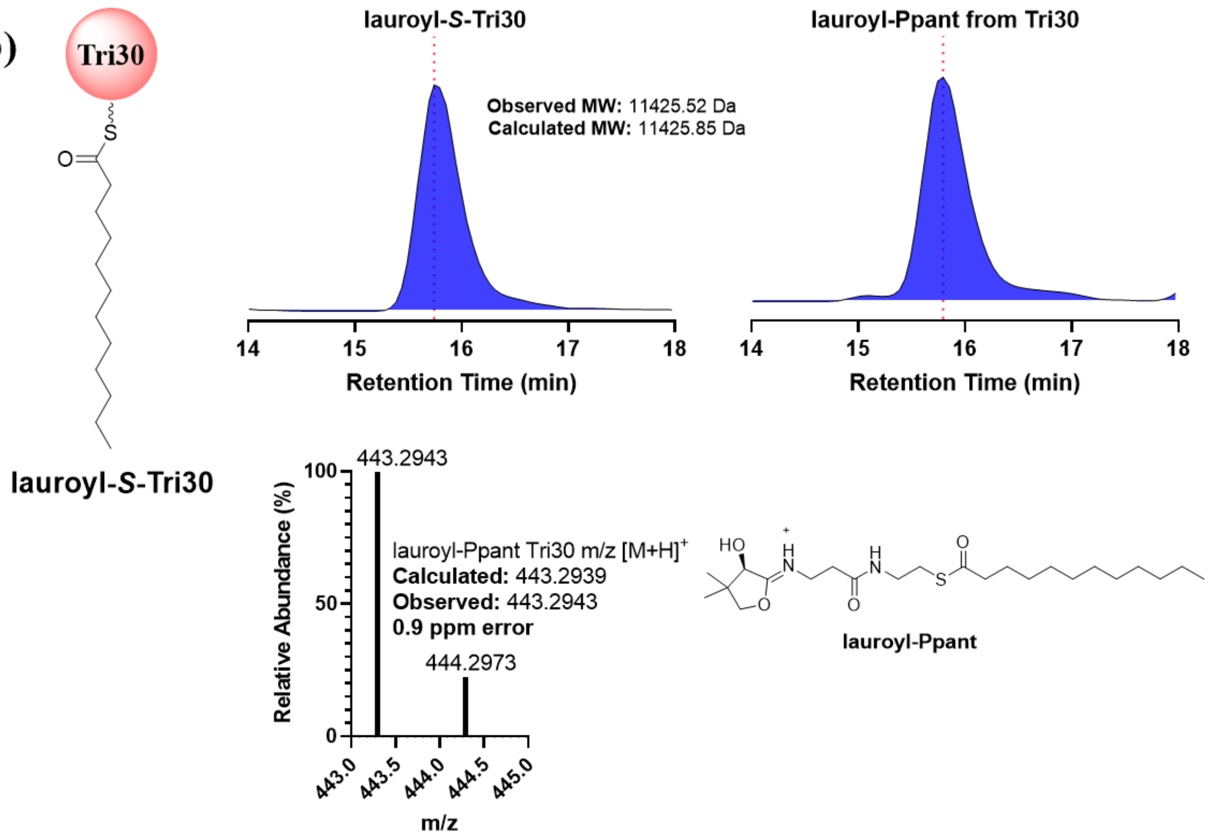

Tri30
S
O
lauroyl-S-Tri30
lauroyl-S-Tri30
Observed MW: 11425.52 Da
Calculated MW: 11425.85 Da
14
15
16
17
18
Retention Time (min)
lauroyl-Ppant from Tri30
14
15
16
17
18
Retention Time (min)
443.2943
100
50
0
Relative Abundance (%)
lauroyl-Ppant Tri30 m/z [M+H]+
Calculated: 443.2939
Observed: 443.2943
0.9 ppm error
444.2973
443.0
443.5
444.0
444.5
445.0
m/z
HO
H
N
H
N
O
S
O
O
lauroyl-Ppant

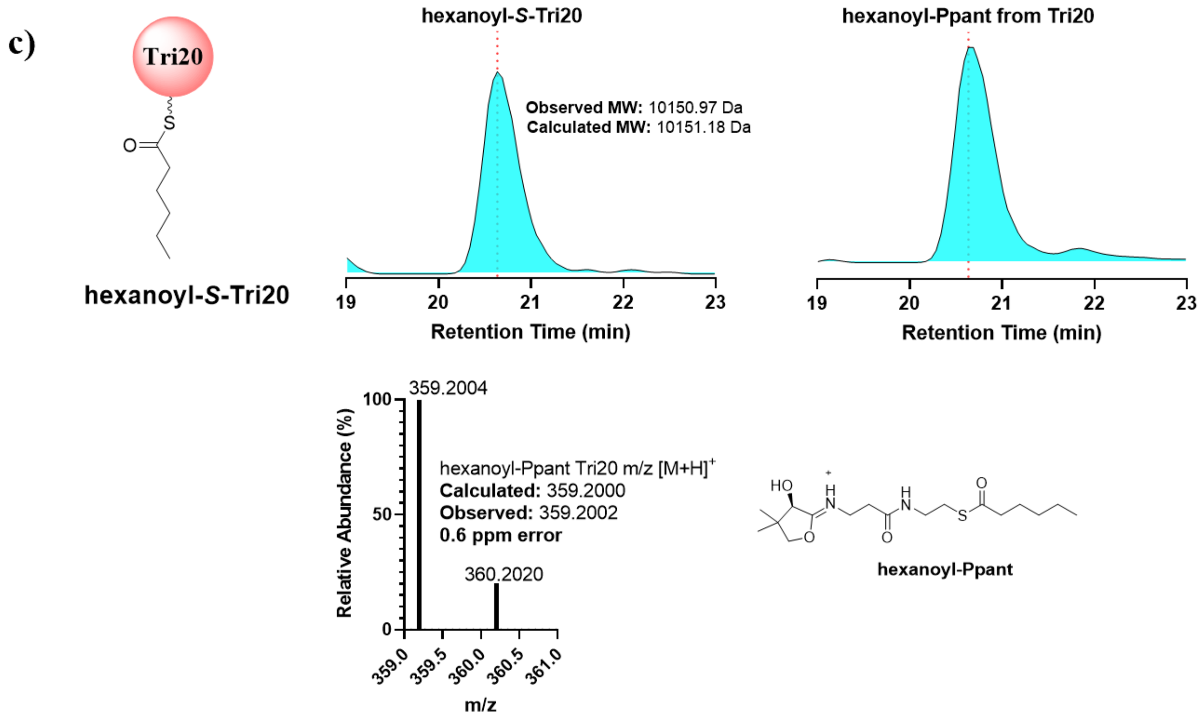


**Supplementary Figure 14.** Acyl-*S*-Tri20 and acyl-*S*-Tri30 formation using Sfp. a) EIC of +7 charge state and Ppant fragment of lauroyl-*S*-Tri20, respectively. b) EIC of +13 charge state and Ppant fragment of lauroyl-*S*-Tri30, respectively. c) EIC of +8 charge state and Ppant fragment of hexanoyl-*S*-Tri20, respectively.

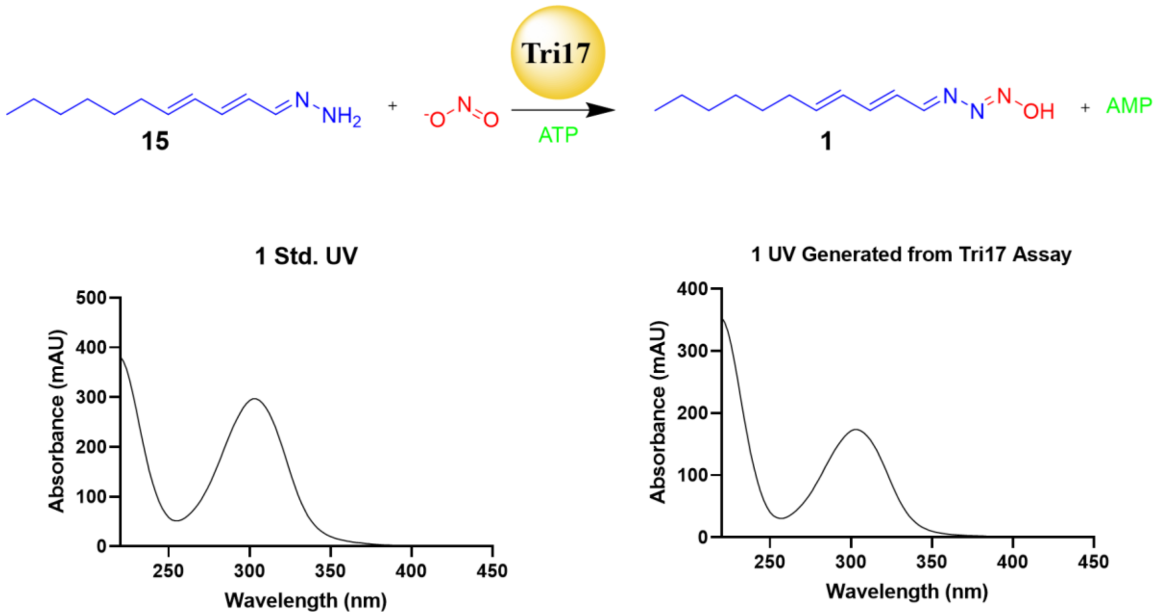


**Supplementary Figure 15.** UV profile of **1**. UV spectra of **1** isolated from WT *S. aureofaciens* and the Tri17 *in vitro* biochemical assay, respectively.

a)

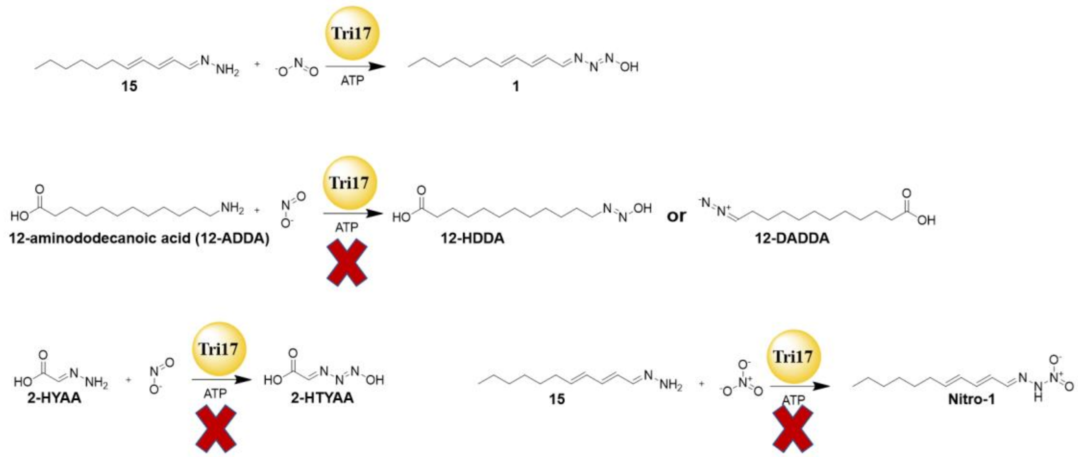


b)

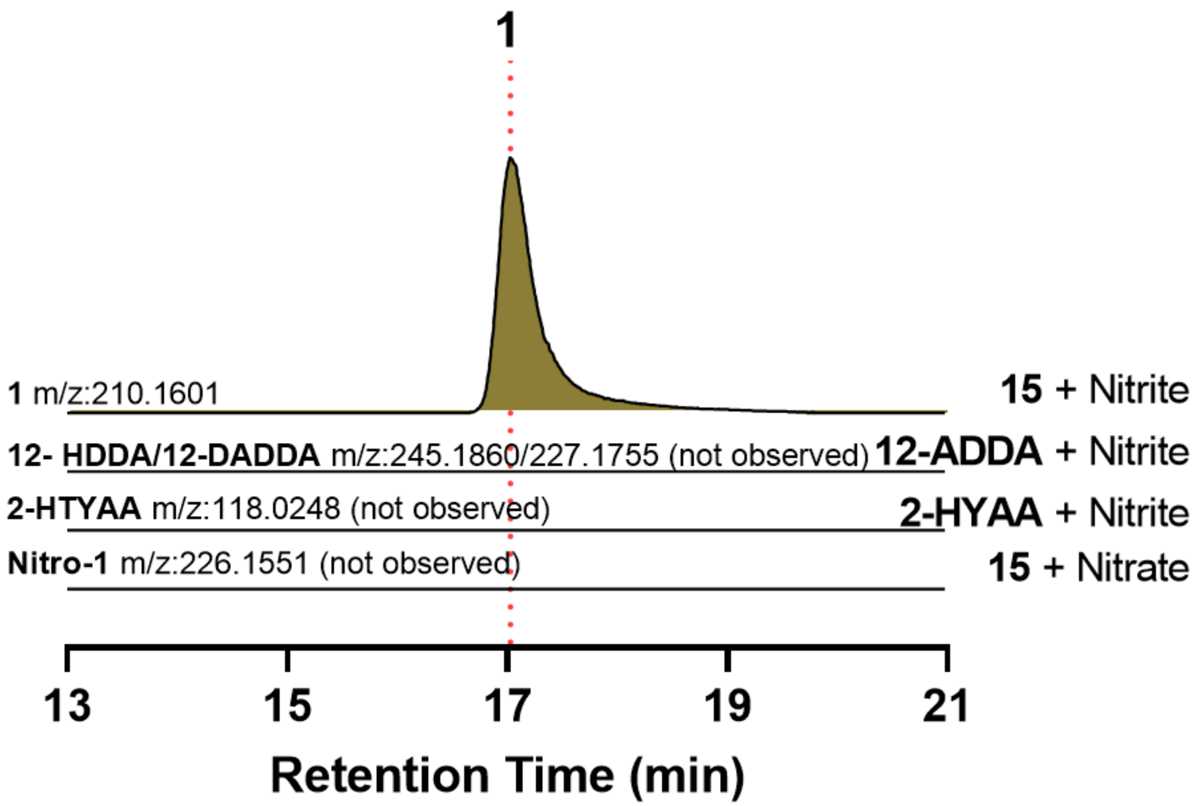


**Supplementary Figure 16.** Substrate specificity of Tri17. a) Reaction schemes of different substrates tested in Tri17 assays. **15** and nitrite were used as positive controls. b) EICs of expected products from each reaction. Tri17 was shown to be "hydrazone", chain length, and nitrite specific. Untargeted comparative metabolomics were also performed for each reaction, showing no new products from the reactions not including **15** and nitrite.

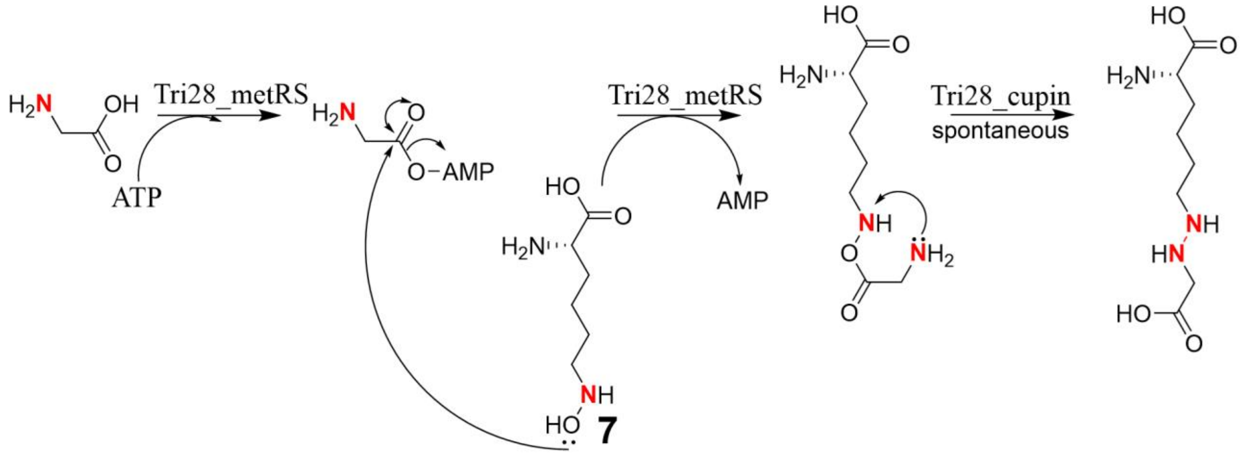


**Supplementary Figure 17.** Proposed mechanism for N-N bond formation by Tri28. Tri28_metRS adenylates glycine to form a glycyl-AMP intermediate, which undergoes a nucleophilic attack by **7** to form an ester intermediate. The cupin domain is proposed to promote the rearrangement of the ester intermediate to generate the hydrazine linkage, although this step may also occur spontaneously based on domain dissection results.

### IV. Supplementary Notes

**Supplementary Note 1.** Structural elucidation of purified **5**.

**Structural elucidation of 5.** The intermediate compound **5** was isolated as a yellow amorphous solid. It had the molecular formula $C_{12}H_{13}N_3O_3$ based on its positive mode HRMS, which showed an intense peak at m/z 248.1032 $[M + H]^+$, corresponding to $C_{12}H_{14}N_3O_3^+$ (calcd 248.1030). Inspection of 1D and 2D NMR spectroscopic data of **5** in comparison with those of triacsin C suggested that intermediate **5** was also a N-hydroxytriazene compound, with a similar structure to triacsin C. However, four $sp^3$ carbon signals were deshieldingly shifted from $\delta_C$ 35.4 (C-9), 34.1 (C-12), 22.0 (C-13), and 13.5 (C-14) *ppm* in triacsin C to $\delta_C$ 139.6 (C-9), 132.9 (C-12), 142.6 (C-13), and 125.0 (C-14) *ppm* in **5**, indicating that these four carbons were not $sp^3$ hybridized, but $sp^2$ hybridized in **5**. The unsaturated long chain of **5** was proposed by analysis of MS2 spectrum with a relative peak at m/z 158.0962, corresponding to $C_{11}H_{12}N^+$ (calcd 158.0965), moreover corroborated by COSY and HMBC cross-peaks. The addition of one carbon and two oxygen atoms in the formula of **5**, together with $^2J$ $^1H$-$^{13}C$ HMBC correlation between H-14 ($\delta_H$ 5.94) and C-15 ($\delta_C$ 167.8), indicated the presence of carboxylic acid group as the other terminal of the unsaturated long chain. Furthermore, the configurations of carbon carbon double bonds were assigned as *E* based on the coupling constants of protons.

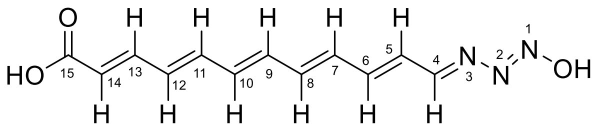


**5**

| Table 2. The NMR Data of **5** in DMSO-$d_6$ | | |
|---|---|---|
| Position | $\delta_H$ (*J* in Hz) | $\delta_C$ (C type) |
| 4 | 8.41 d 10.0 | 167.4, CH |
| 5 | 6.59 m | 128.2, CH |
| 6 | 7.16 dd 15.0, 11.2 | 146.6, CH |
| 7 | 6.68 m | 133.0, CH |
| 8 | 6.66 m | 133.4, CH |
| 9 | 6.75 dd 14.7, 9.2 | 139.6, CH |
| 10 | 6.62 m | 136.1, CH |
| 11 | 6.74 dd 14.7, 9.3 | 139.4, CH |
| 12 | 6.56 dd 14.7, 11.0 | 132.9, CH |
| 13 | 7.14 dd 15.2, 11.0 | 142.6, CH |
| 14 | 5.94 d 15.2 | 125.0, CH |
| 15 | | 167.8, C |
| N-OH | 8.36 s | |

.

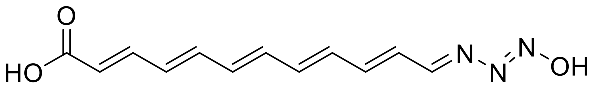

**5** ( [1]H NMR, DMSO-$d_6$ at 900 MHz)

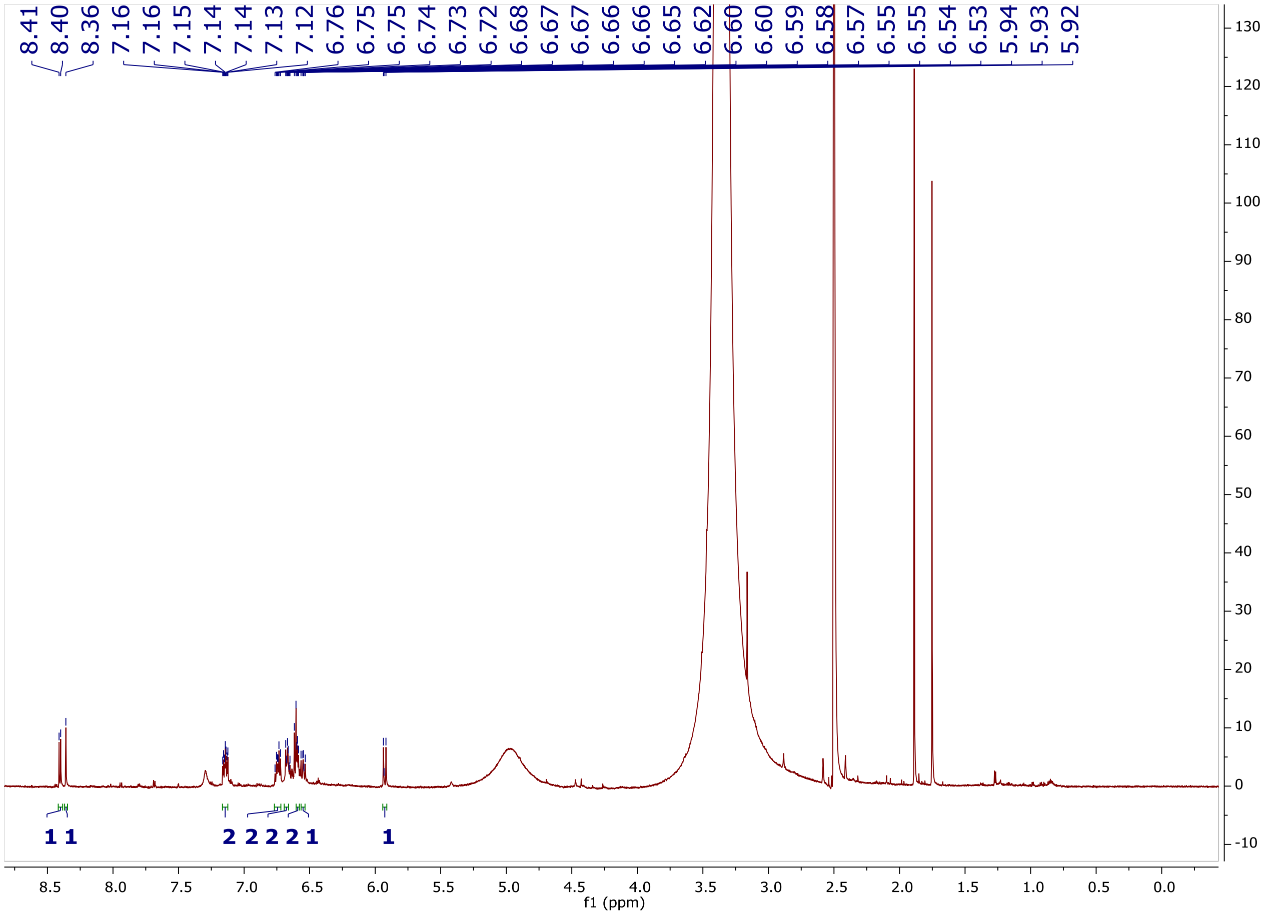

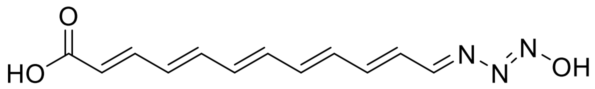


**5** $^{1}$H-$^{1}$H COSY, DMSO-$d_6$ at 900 MHz

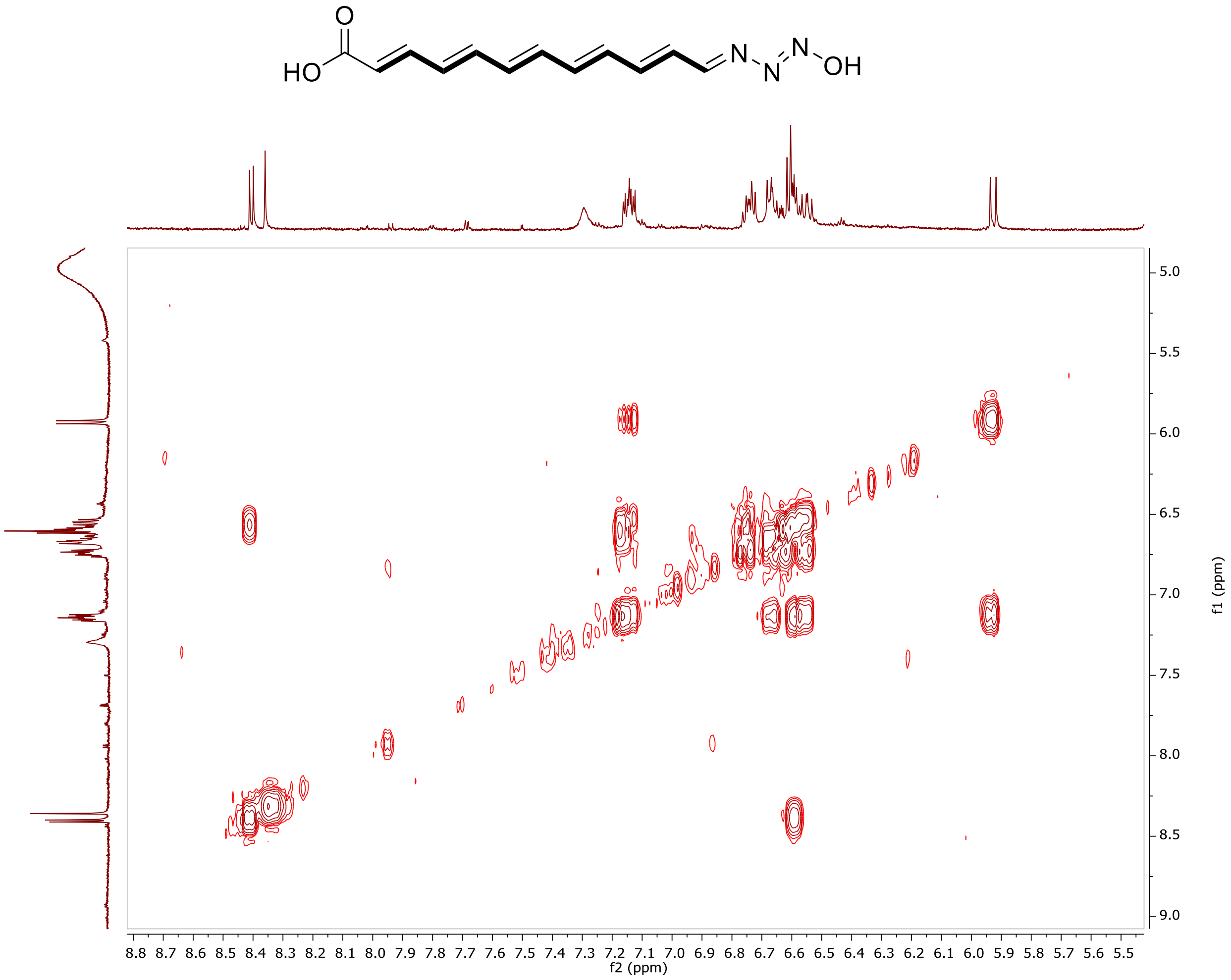

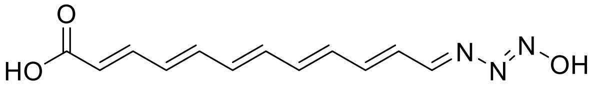

**5** ($^{1}$H-$^{13}$C HSQC, DMSO-$d_6$ at 900 MHz)

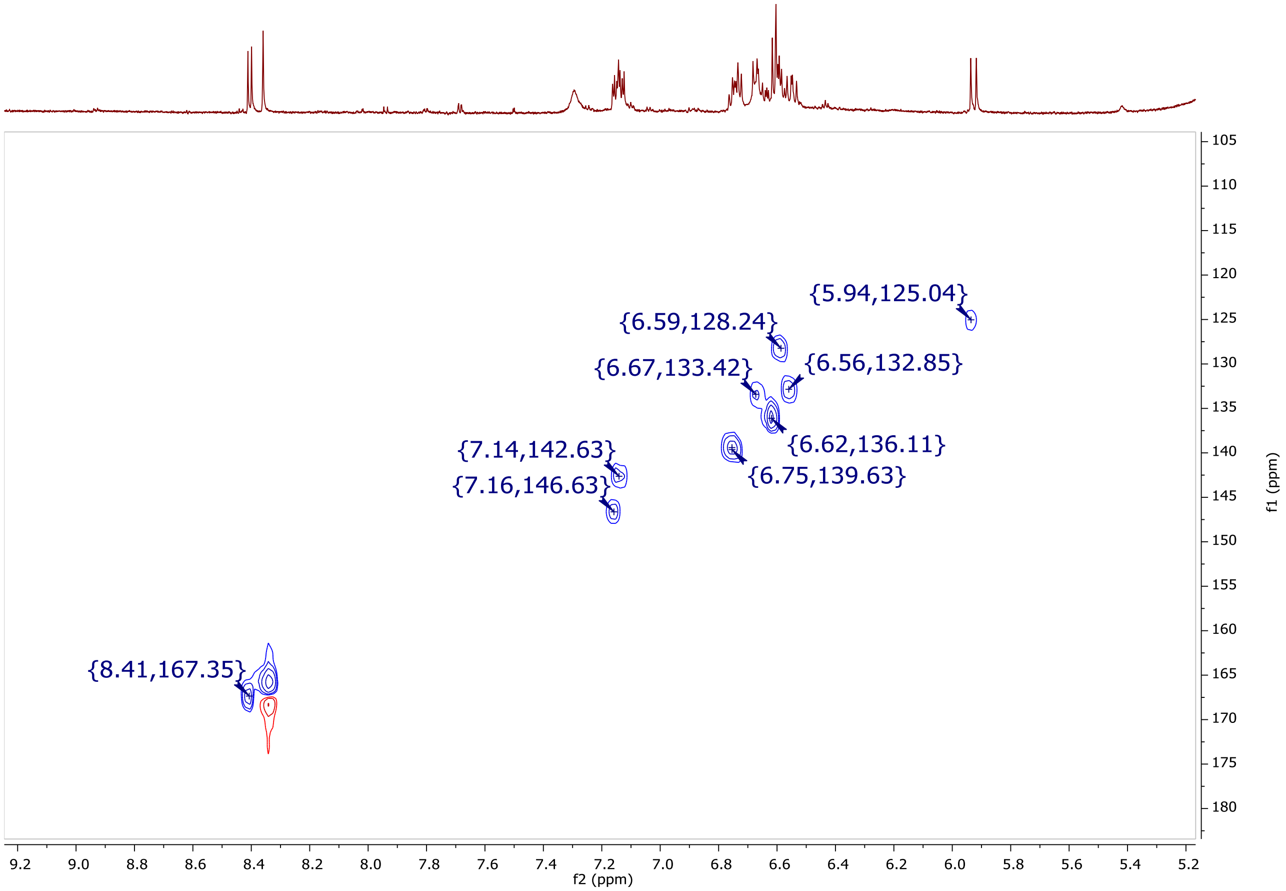

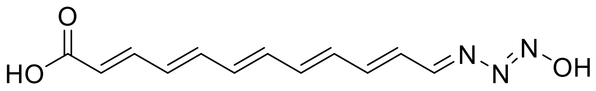

**5** ($^{1}$H-$^{13}$C HMBC, DMSO-$d_6$ at 900 MHz)

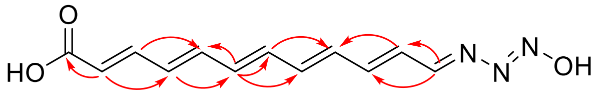

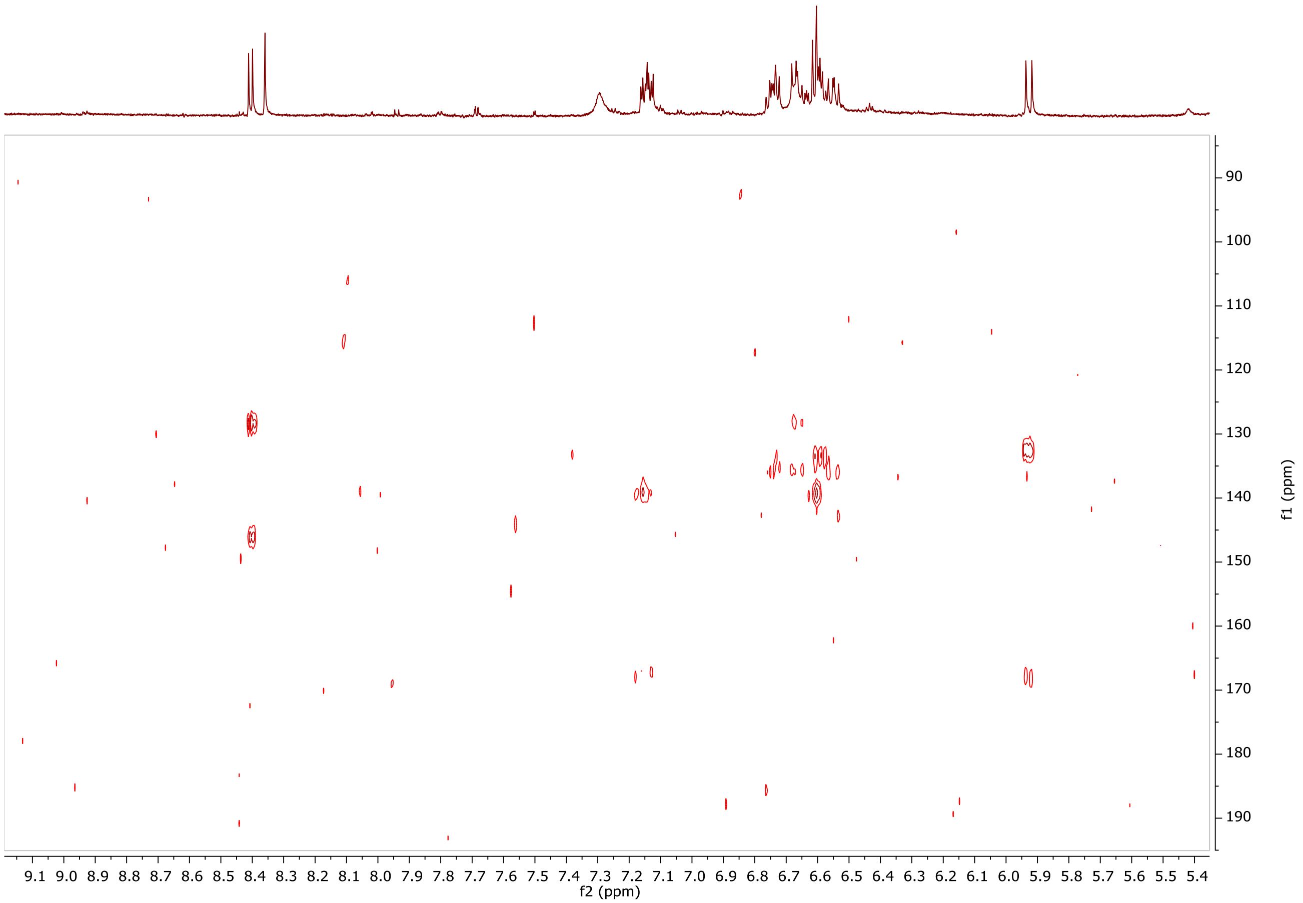

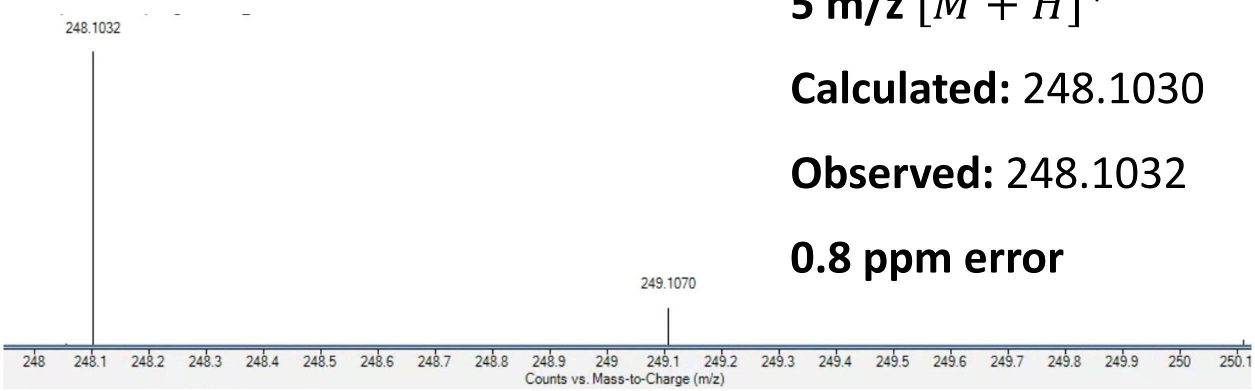


**5 m/z** $[M + H]^+$

**Calculated:** 248.1030

**Observed:** 248.1032

**0.8 ppm error**

**MS2 Fragment** $[M + H]^+$

**Calculated:** 158.0965

**Observed:** 158.0962

**1.9 ppm error**

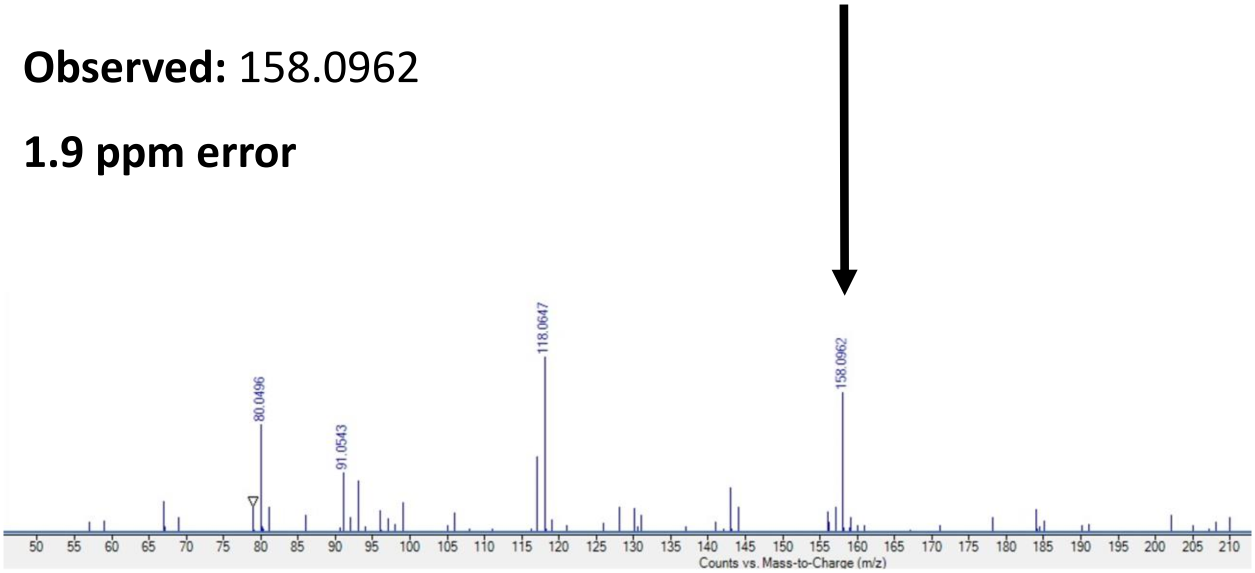


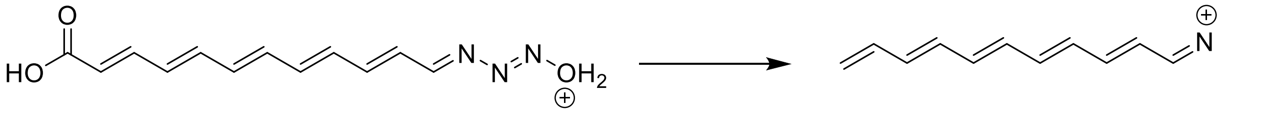


Chemical Formula: $C_{12}H_{14}N_3O_3^+$
m/z: 248.1030 (100.0%), 249.1064 (13.0%), 249.1001 (1.1%)

Chemical Formula: $C_{11}H_{12}N^+$
m/z: 158.0965 (100.0%), 159.0998 (11.9%)

**Supplementary Note 2.** NMR characterization of Tri31 reaction**.**

**Structural elucidation of hexanoyl-HAA.** The hexanoyl-HAA was generated as a yellow amorphous solid by the Tri31 assay described in the Methods section. The positive ion HRESIMS data revealed a peak at m/z 189.1233, corresponding to a molecular formula of $C_8H_{16}N_2O_3$. Inspection of its $^1$H NMR and $^1$H-$^{15}$N HMBC spectroscopic data in comparison with those of the known starting material HAA revealed that the product from the Tri31 enzymatic reaction was a long chain analog of HAA with an amide linkage. This assignment was based on the shift of the $^1$H-$^{15}$N HMBC of nitrogen bearing methylene from the signal between $\delta_H$ 3.65 and $\delta_N$ 66.6 (a primary amine) in HAA to the signal between $\delta_H$ 3.69 and $\delta_N$ 144.7 (an amide) in hexanoyl-HAA. Moreover, the other nitrogen of hydrazine didn't react with hexanoyl-CoA, which could be confirmed via the $^2J$ $^1$H-$^{13}$N HMBC cross-peak between -NH-CO- ($\delta_H$ 9.68) and -$CH_2$-NH- ($\delta_N$ 61.7). Therefore, we validated that the Tri31 reaction only occurred at the terminal -$NH_2$ group. $^1$H NMR (900 MHz, DMSO-$d_6$): $\delta_H$ 9.68 (s, 1H), 3.69 (s, 2H), 2.09 (t, $J$ = 7.4 Hz, 2H), 1.48 (m, 2H), 1.22 (m, 4H), 0.85 (t, $J$ = 7.1 Hz, 3H); $^{15}$N NMR ($^1$H-$^{15}$N HMBC at 900 MHz, DMSO-$d_6$): $\delta_N$ 61.7, 144.8.

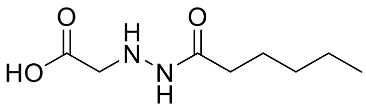


**hexanoyl-HAA**

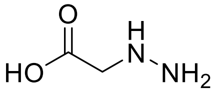


**HAA**

[1]H NMR (900 MHz, DMSO-$d_6$): $\delta_H$ 3.65 (s, 2H).

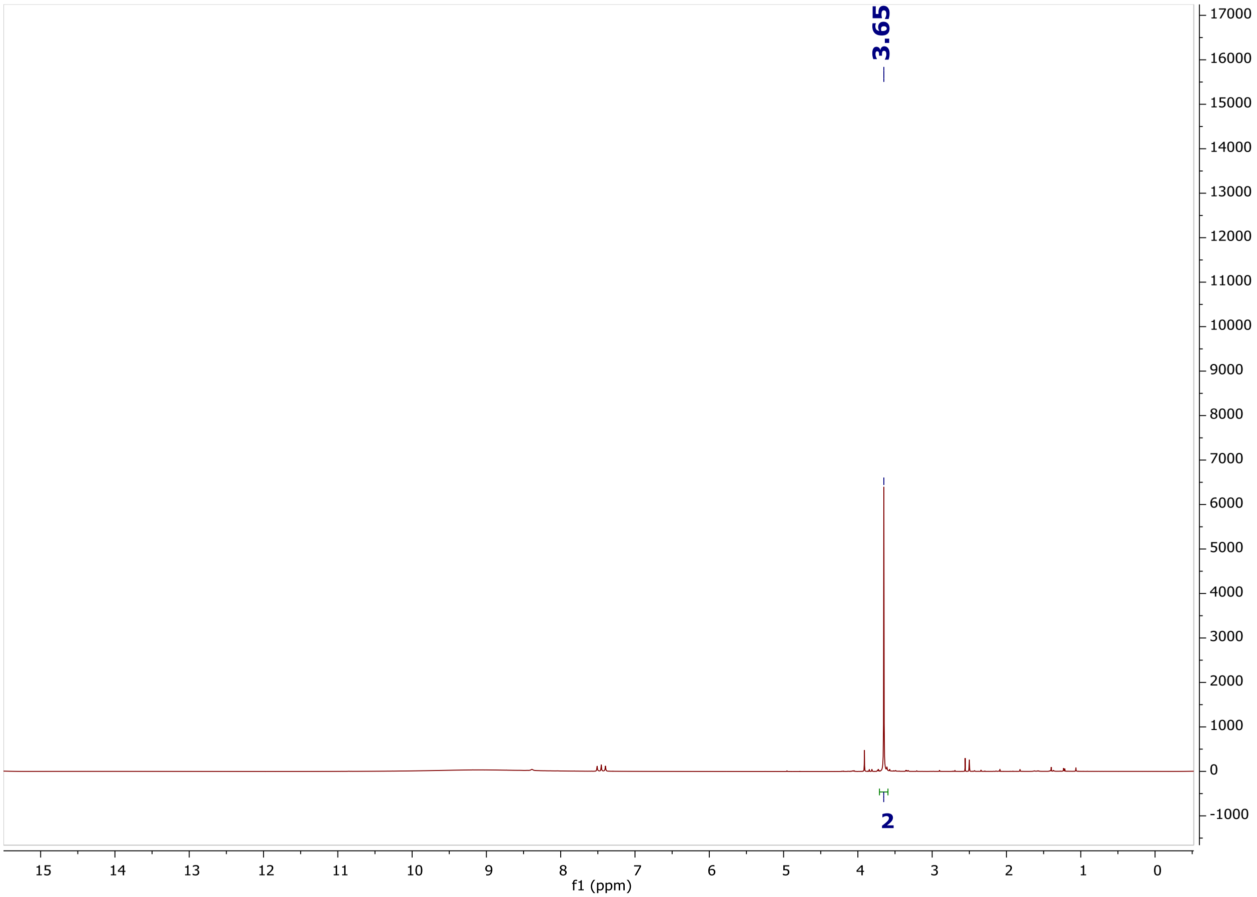

$^{15}N$ NMR ($^{1}H$-$^{15}N$ HMBC at 900 MHz, DMSO-$d_6$): $\delta_N$ 62.7, 66.6.

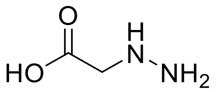


**HAA**

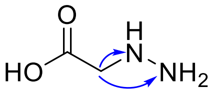


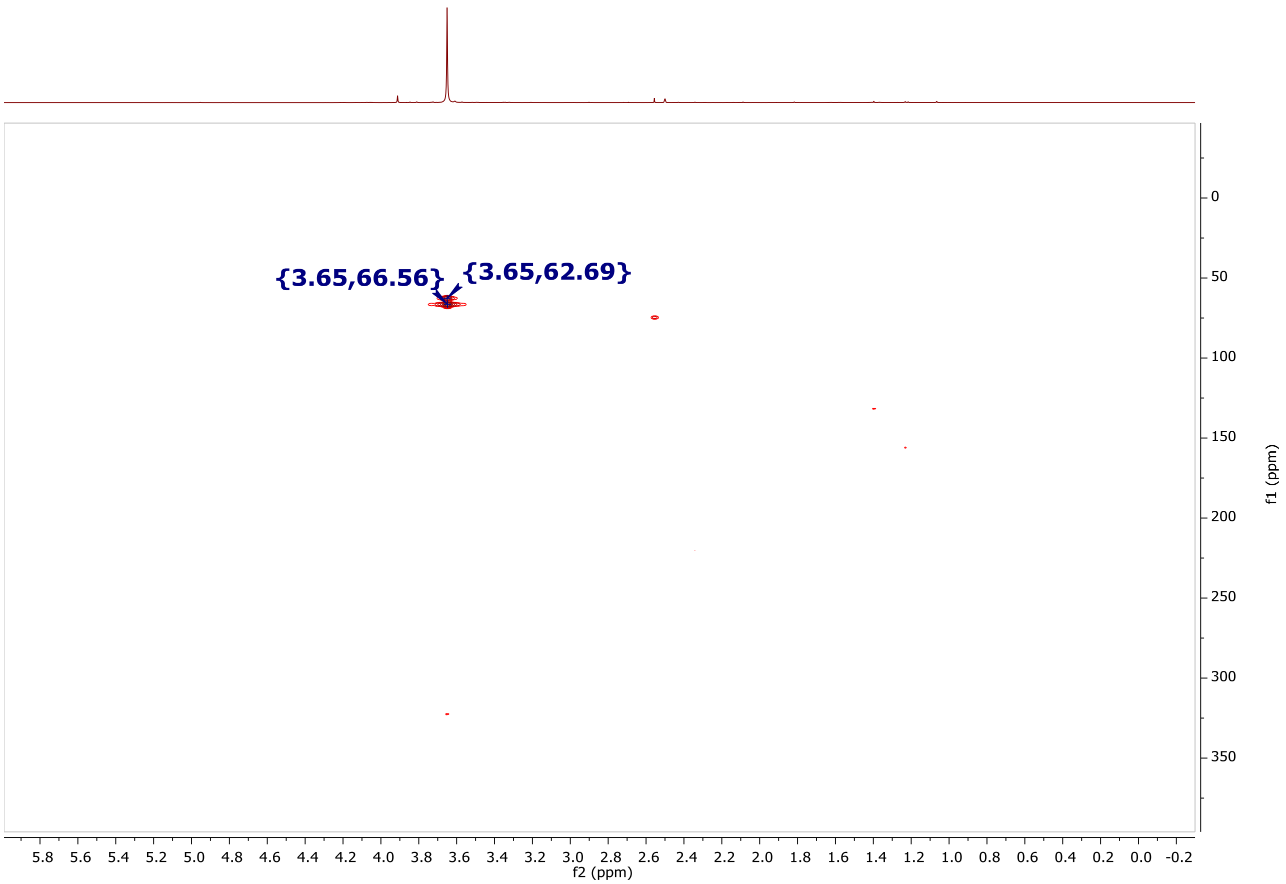

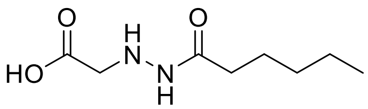


**hexanoyl-HAA** ($^1$H NMR, DMSO-$d_6$ at 900 MHz)

$^1$H NMR (900 MHz, DMSO-$d_6$): $\delta_H$ 9.68 (s, 1H), 3.69 (s, 2H), 2.09 (t, $J$ = 7.4 Hz, 2H), 1.48 (m, 2H), 1.22 (m, 4H), 0.85 (t, $J$ = 7.1 Hz, 3H); $^{15}$N NMR ($^1$H-$^{15}$N HMBC at 900 MHz, DMSO-$d_6$): $\delta_N$ 61.7, 144.8.

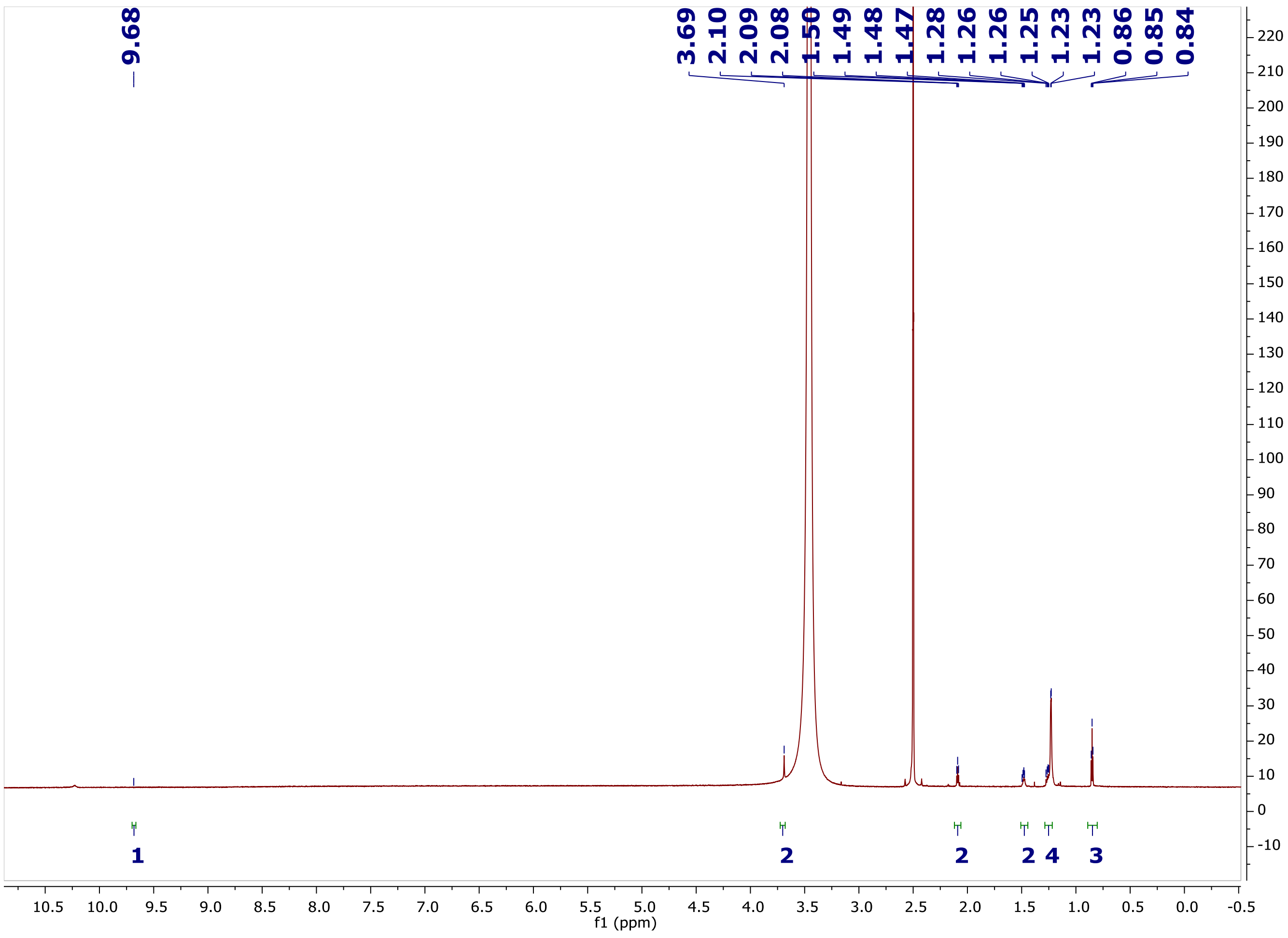

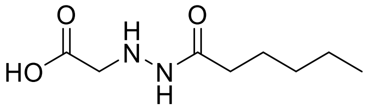

**hexanoyl-HAA**($^1$H-$^{15}$N HMBC, DMSO-$d_6$ at 900 MHz)

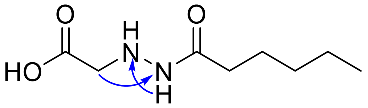

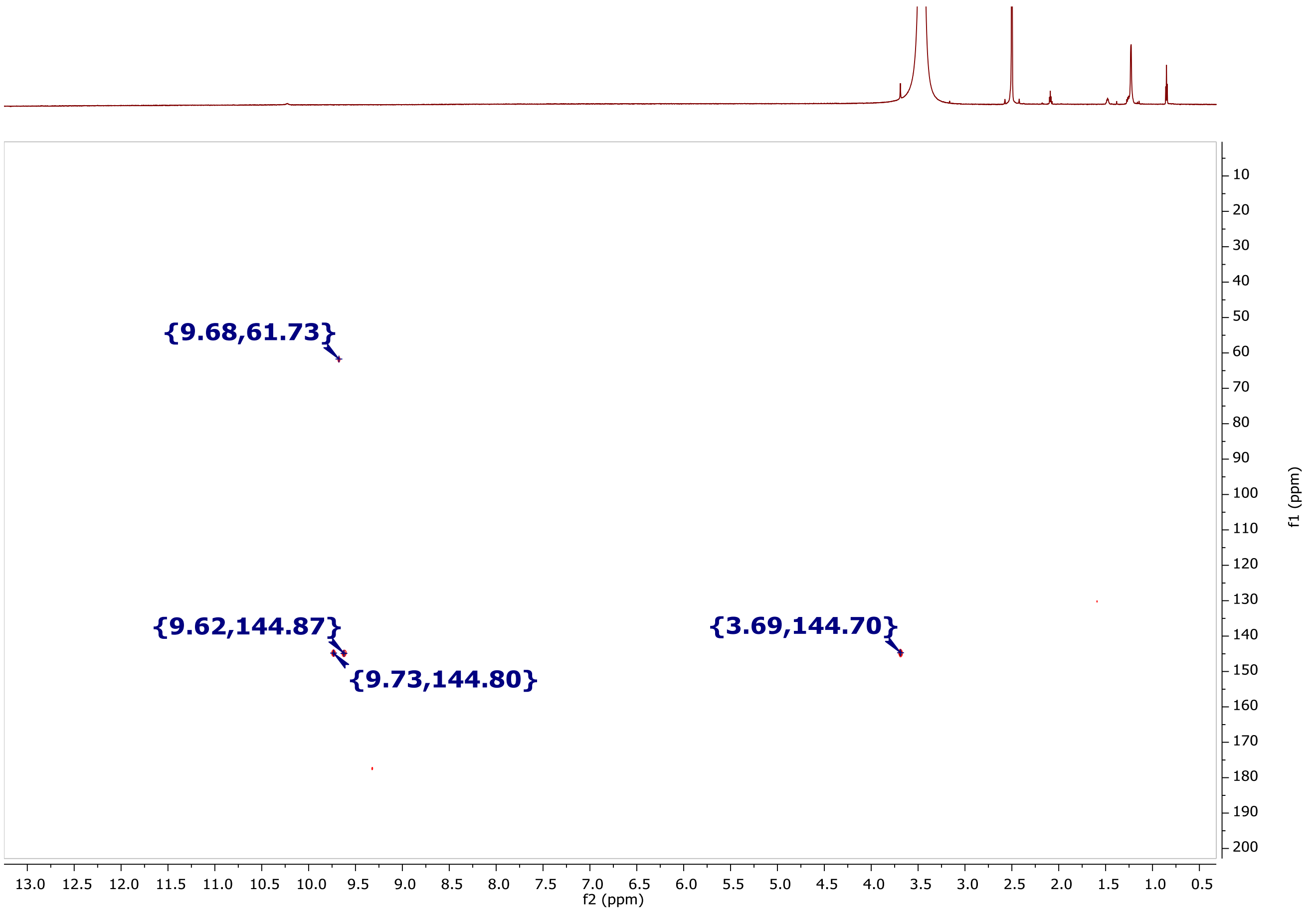

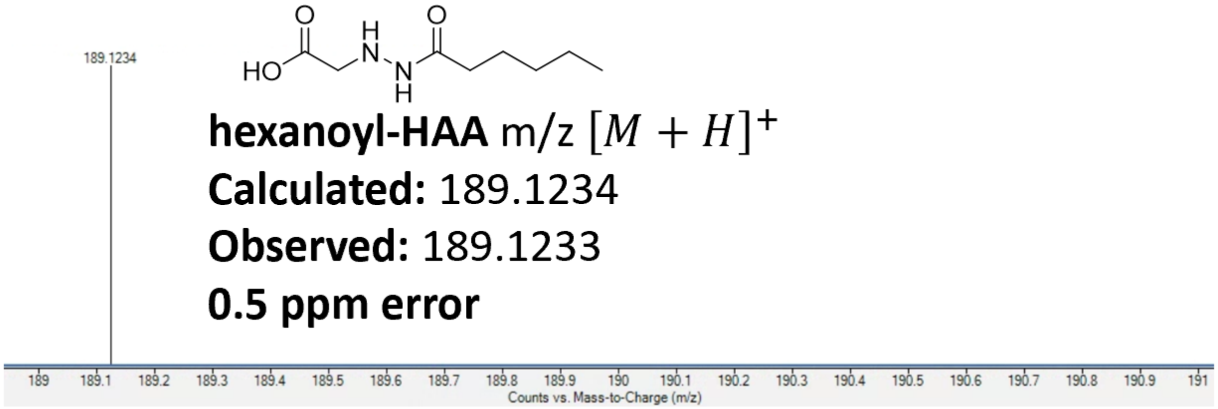
189.1234
HO
O
H
N
N
H
O
hexanoyl-HAA m/z $[M + H]^+$
Calculated: 189.1234
Observed: 189.1233
0.5 ppm error
189
189.1
189.2
189.3
189.4
189.5
189.6
189.7
189.8
189.9
190
190.1
190.2
190.3
190.4
190.5
190.6
190.7
190.8
190.9
191
Counts vs. Mass-to-Charge (m/z)

**Supplementary Note 3.** Structural elucidation of synthetic 2-HYAA.

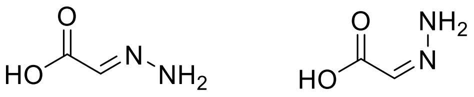

**2-HYAA**

**Structural elucidation of *cis*-2-HYAA and *trans*-2-HYAA.** 2-HYAA was synthesized as a mixture of two geometric isomers. Its $^{1}H$ NMR and HSQC spectra simply displayed signals for two olefinic methines ($\delta_H$ 6.96, $\delta_C$ 129.1 and $\delta_H$ 6.62, $\delta_C$ 132.4). Furtherly, the presences of two 2-HYAA isomers were indicated via $^{3}J$ $^{1}H$-$^{13}C$ HMBC correlations between -CH=N- ($\delta_H$ 6.96) and -COOH ($\delta_C$ 168.1), and -CH=N- ($\delta_H$ 6.62) and -COOH ($\delta_C$ 169.0). Normally, the chemical shift of the carbon at the *trans* position was more deshielded than that at the *cis* position, so we assigned the methine at $\delta_H$ 6.96 in *cis*-2-HYAA, while the methine at $\delta_H$ 6.62 in *trans*-2-HYAA. From their integrations of both signals in $^{1}H$ NMR spectrum, we calculated the ratio of *cis:trans* was 1:1.

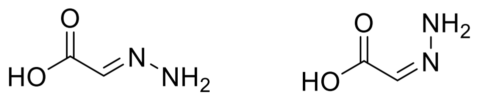


**2-HYAA** ($^1$H NMR, DMSO-$d_6$ at 900 MHz)

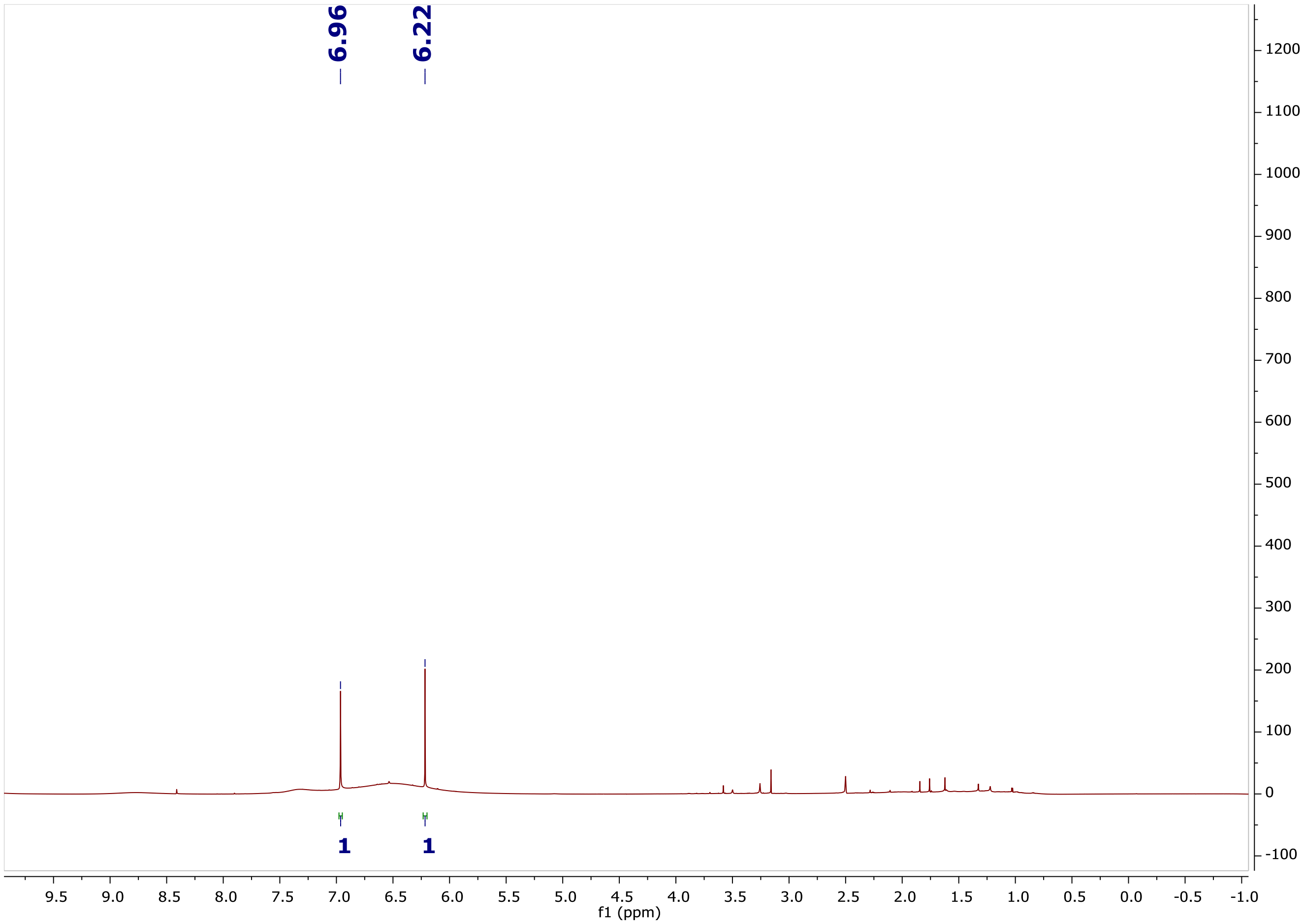

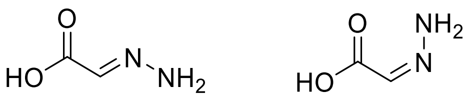

**2-HYAA** ($^1$H-$^{13}$C HSQC, DMSO-$d_6$ at 900 MHz)

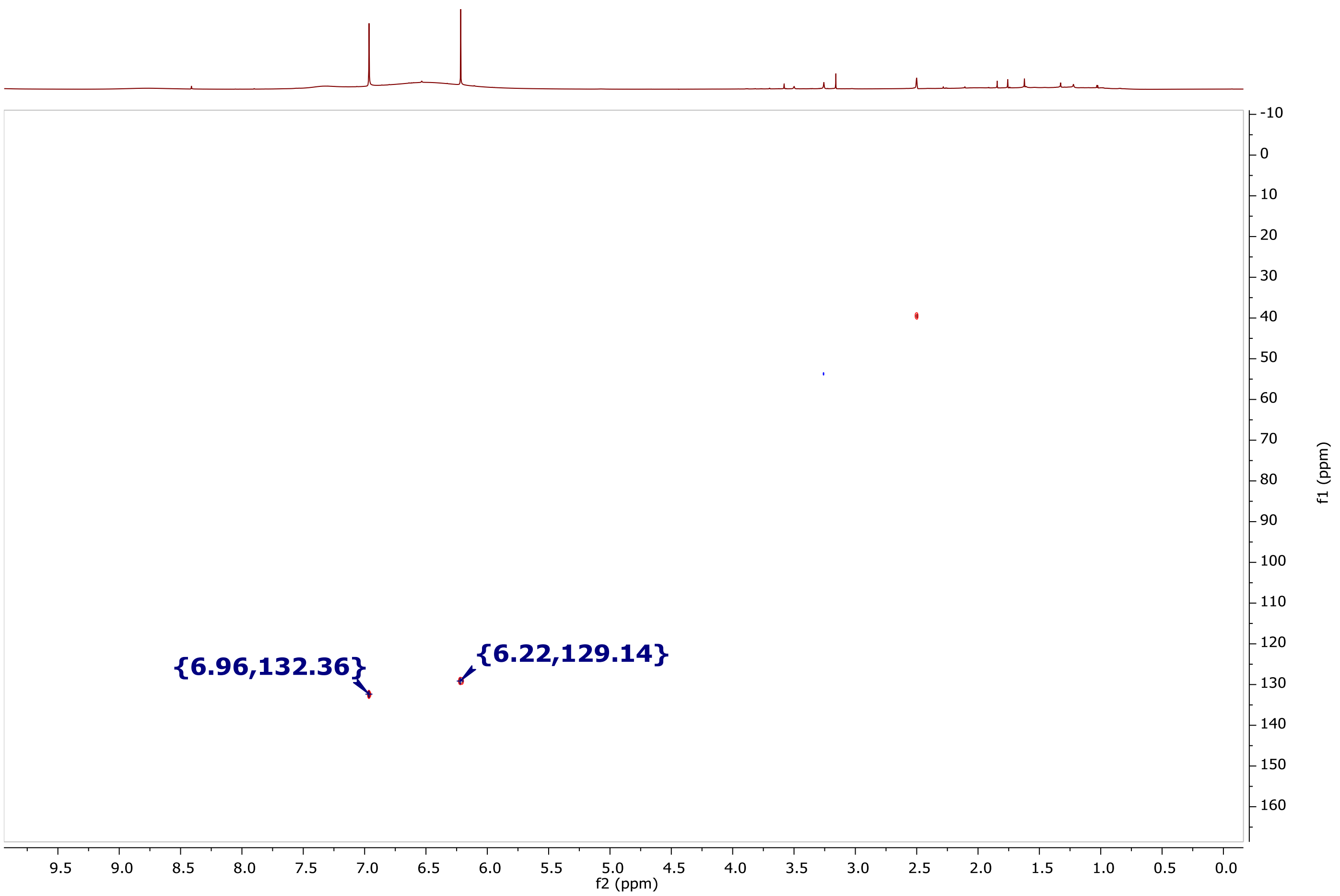

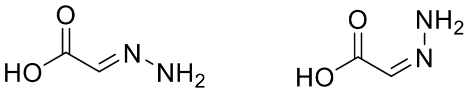

**2-HYAA** ($^1$H-$^{13}$C HMBC, DMSO-$d_6$ at 900 MHz)

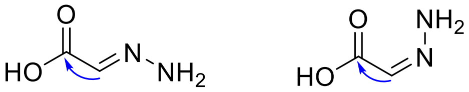

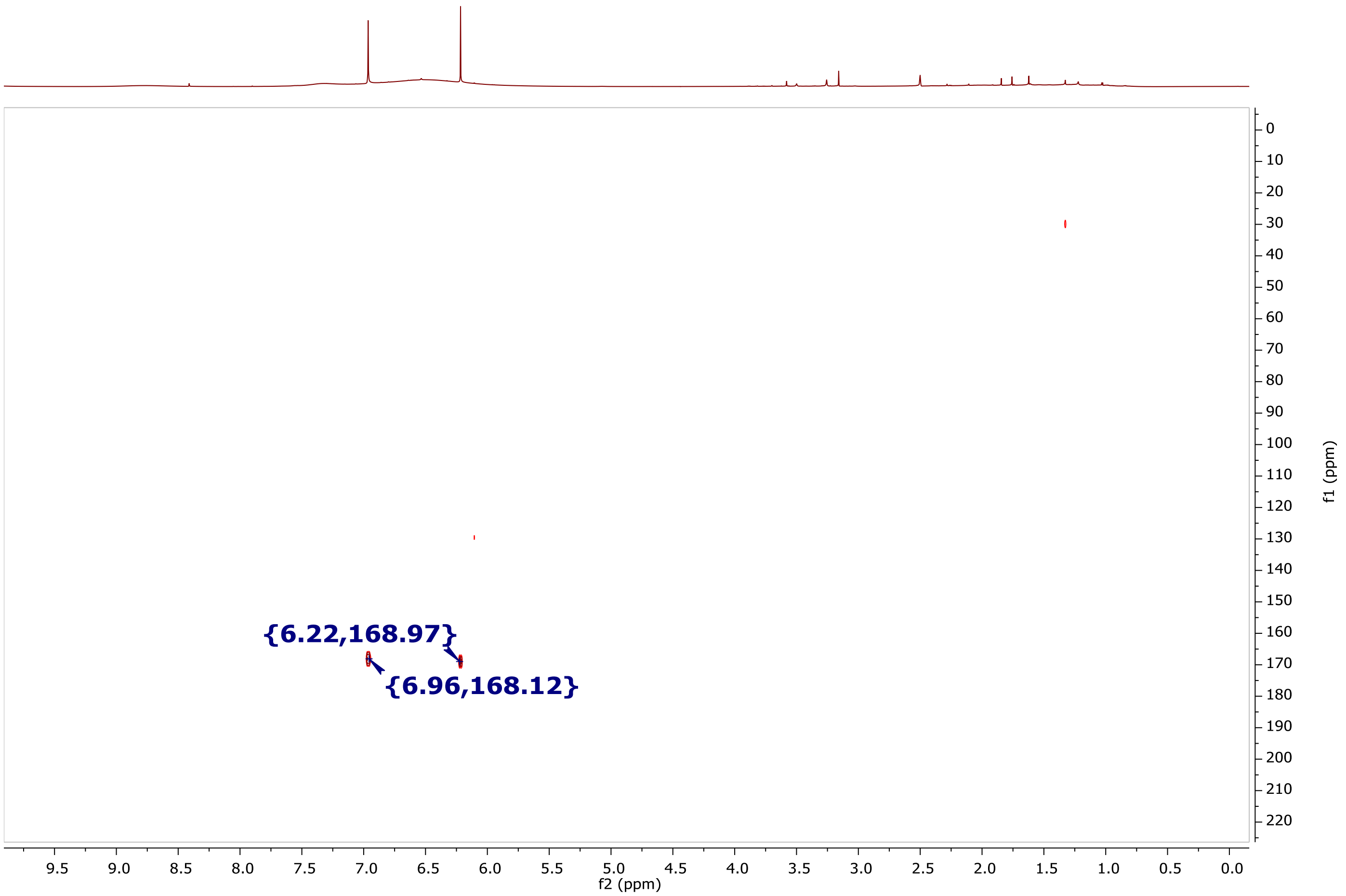

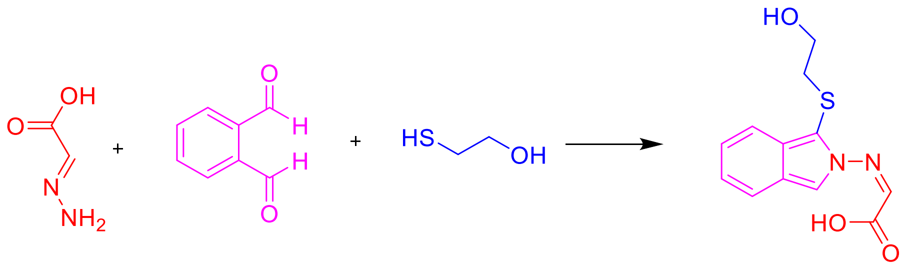


**2-HYAA** **OPTA** **2-ME** **2-HYAA-D**

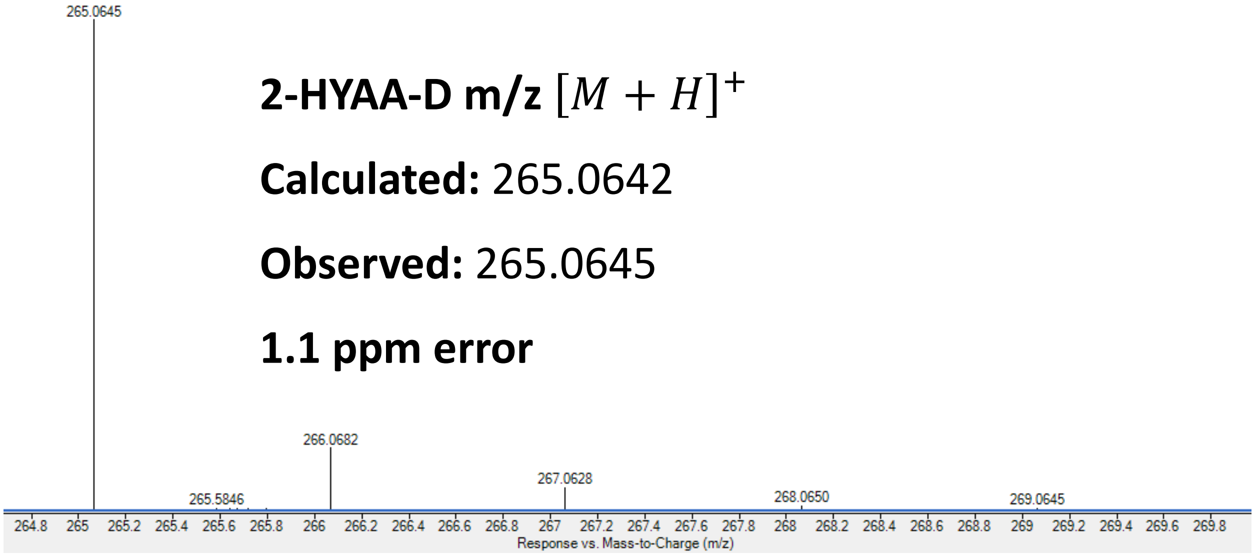


**2-HYAA-D m/z** $[M + H]^{+}$

**Calculated:** 265.0642

**Observed:** 265.0645

**1.1 ppm error**

**2-HYAA-D Std. UV**

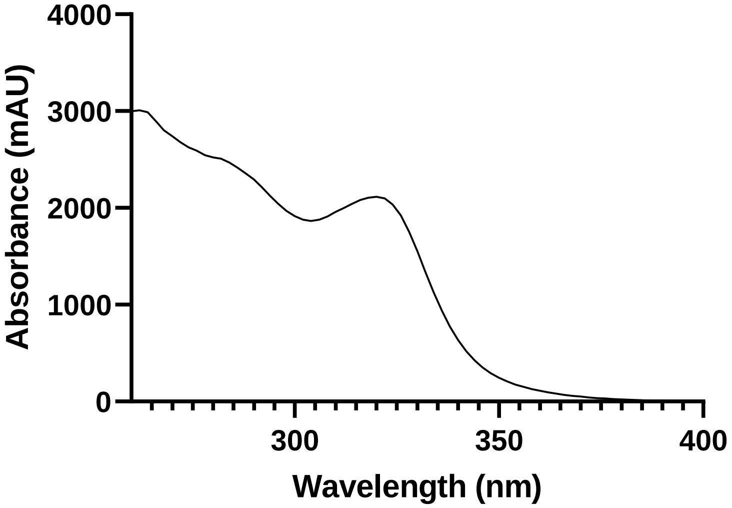

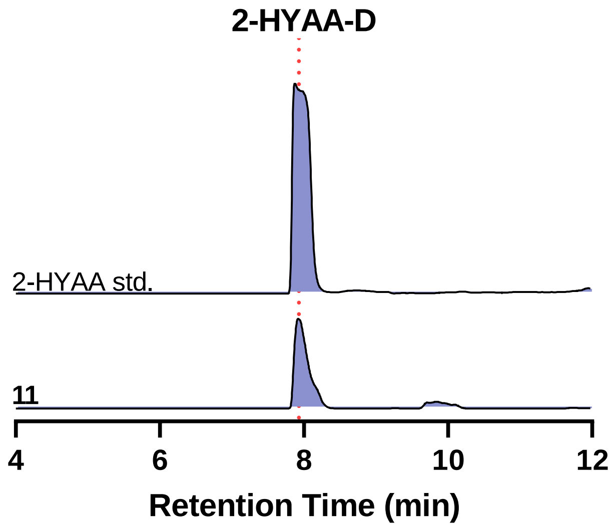


Our NMR characterization indicates a 1:1 *cis* to *trans* 2-HYAA ratio. However, from our LC-UV-HRMS analyses of 2-HYAA-D, we only observed one peak despite various methods utilized to separate putative multiple species. We thus do not have direct evidence for the configuration of the HYAA moiety of **11**. Based on the final structure of the triacsins, we propose **11** is likely in the *E* (*trans*) configuration as all congeners were reported to be *trans* at that position.

**Supplementary Note 4.** Structural confirmation of synthetic **15**.

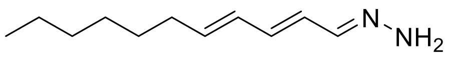


**15**

**Structural elucidation of compound 15.** Compound **15** was purified as a yellow amorphous solid via organic synthesis. Its positive ion HRESIMS revealed a peak for a protonated molecular ion at m/z 181.1699, corresponding to the molecular formula $C_{11}H_{20}N_2$. Its $^1$H NMR and HSQC spectra displayed signals for five olefinic methines ($\delta_H$ 7.34, 6.57, 6.21, 6.11 and 5.72), five methylenes ($\delta_H$ 2.06, 2H, 1.35, 2H, and 1.26, 6H), and one methyl groups ($\delta_H$ 0.85). Consequently, the skeleton of long chain was indicated by COSY correlations, while the hydrazone moiety was indicated by the $^3J$ HMBC correlations between -CH= ($\delta_H$ 6.11) and =N- ($\delta_N$ 345.5), as well as $^2J$ HMBC correlations between -CH= ($\delta_H$ 7.34) and =N- ($\delta_N$ 345.5), and -NH$_2$ ($\delta_H$ 6.57) and =N- ($\delta_N$ 345.5).

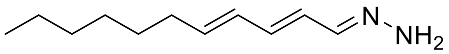


**15** ($^1$H NMR, DMSO-$d_6$ at 900 MHz)

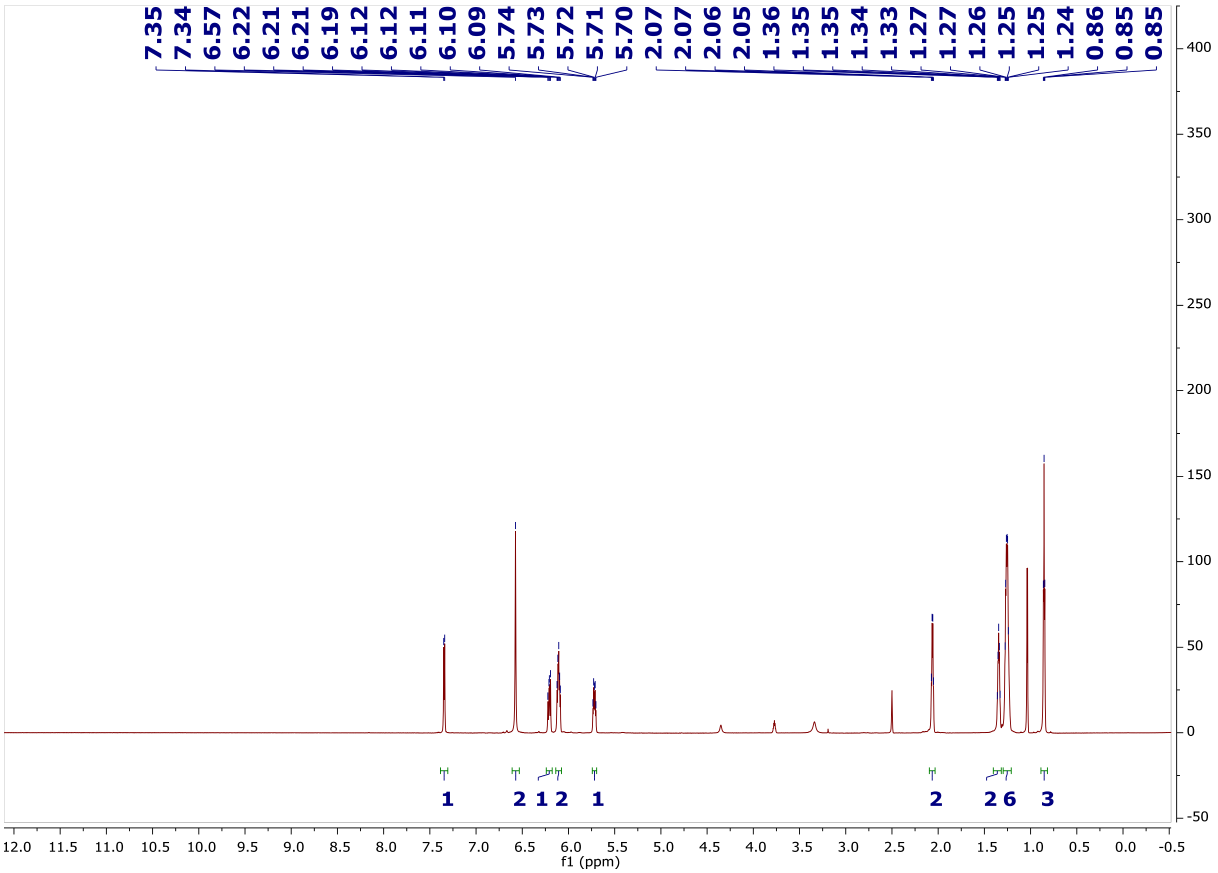

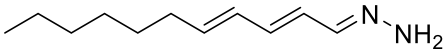


**15** ($^{13}$C NMR, DMSO-$d_6$ at 225 MHz)

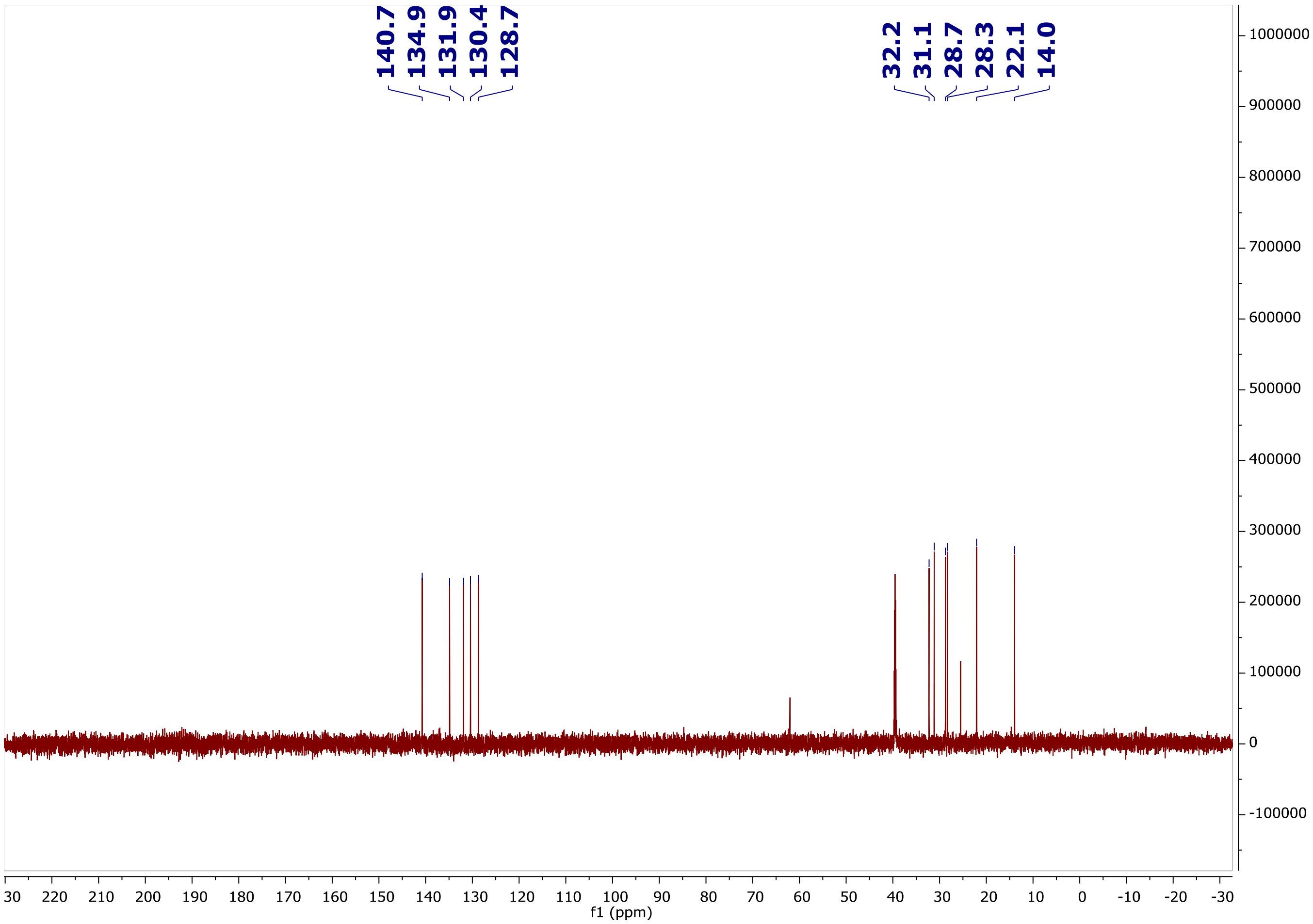

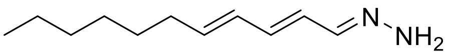


**15** ($^1$H-$^1$H COSY, DMSO-$d_6$ at 900 MHz)

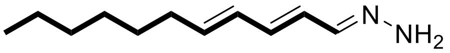


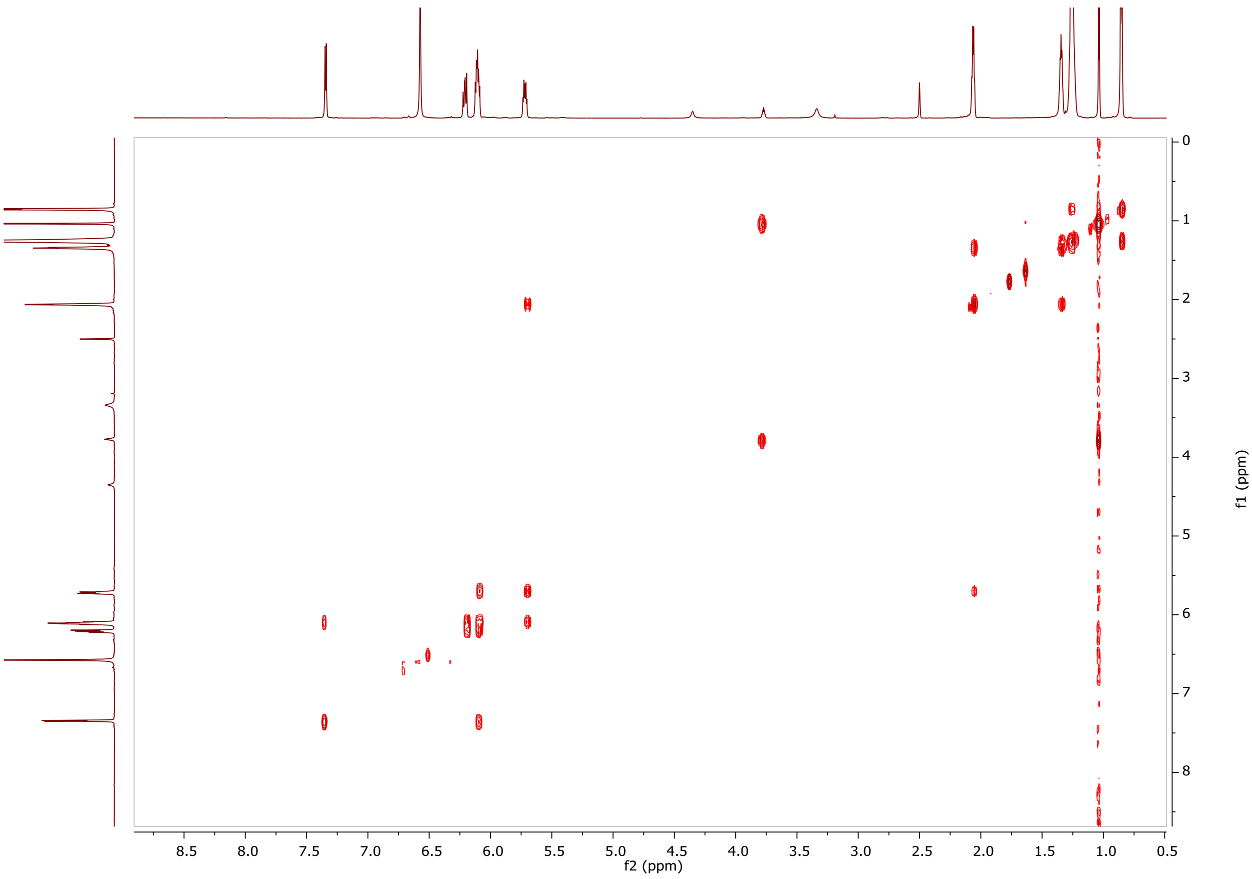

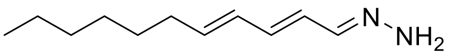


**15** ($^1$H-$^{13}$C HSQC, DMSO-$d_6$ at 900 MHz)

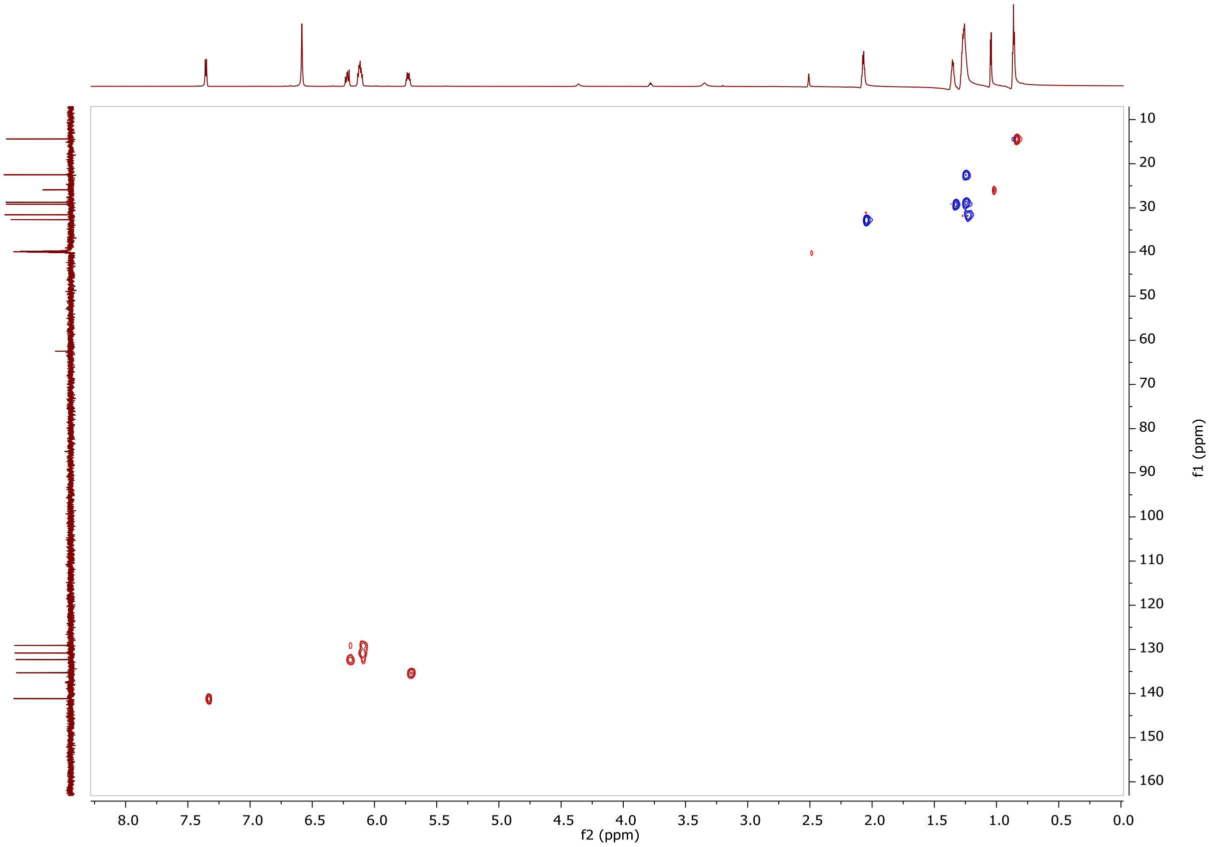

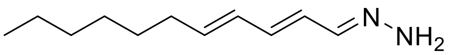


**15** ($^1$H-$^{15}$N HMBC, DMSO-$d_6$ at 900 MHz)

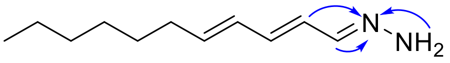


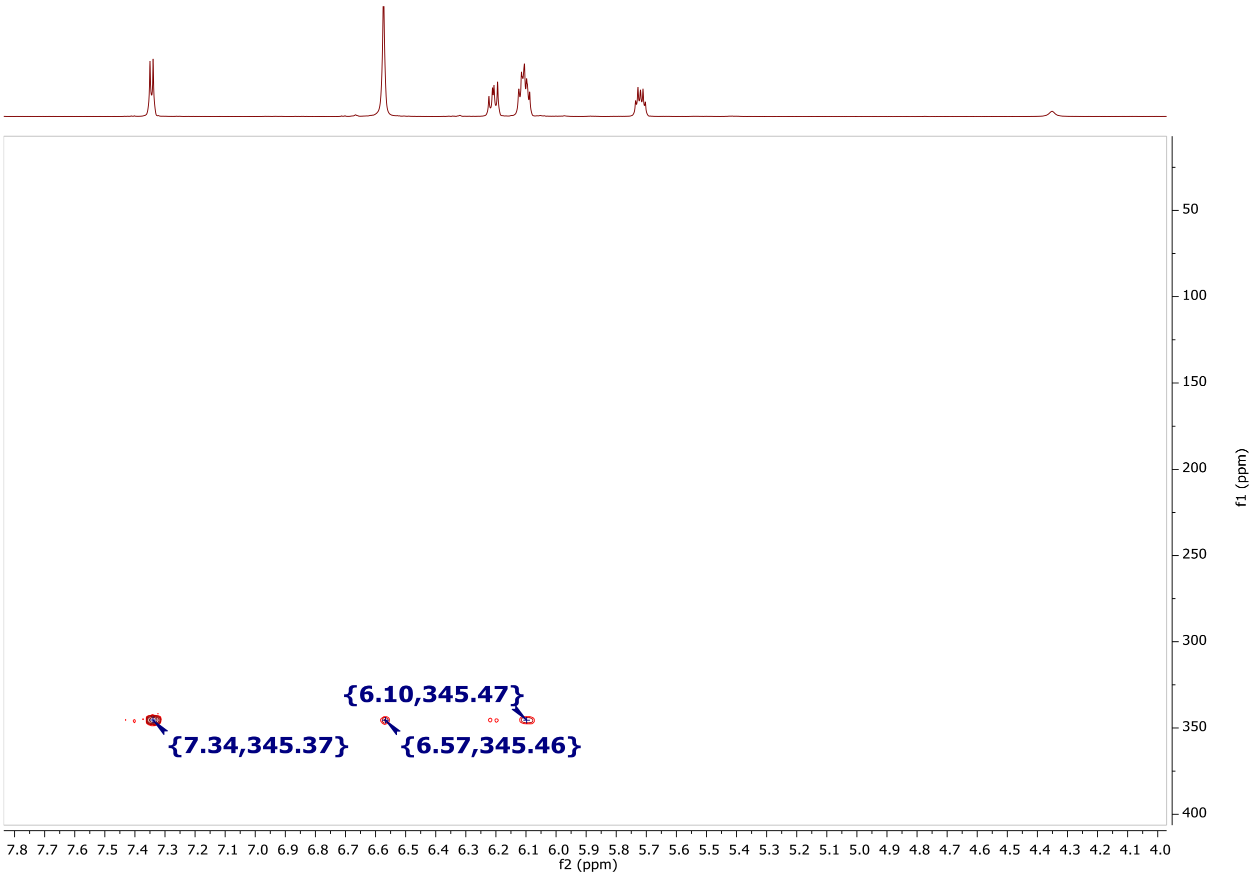


**15 m/z** $[M + H]^+$

**Calculated:** 181.1700

**Observed:** 181.1699

**0.6 ppm error**

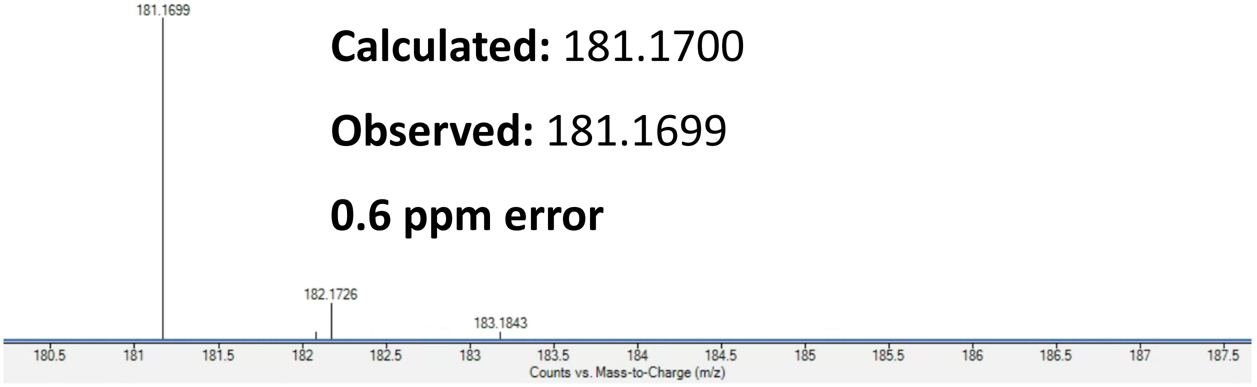

**Supplementary Note 5.** Structural confirmation of purified triacsin A (**1**).

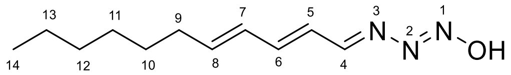


**1**

**Structural elucidation of 1.** The known compound, triacsin A, was isolated as a yellow amorphous solid. The positive ion HRESIMS data displayed a peak for a protonated molecular ion at m/z 210.1597, corresponding to a molecular formula of $C_{11}H_{19}N_3O$. The structure of triacsin A was verified by comparison with previously reported NMR data[1].

Table 1. The NMR Data of **1** in DMSO-$d_6$

| Position | $\delta_H$ ($J$ in Hz) | $\delta_C$ (C type) |
|---|---|---|
| 4 | 8.37 d 9.9 | 167.5, CH |
| 5 | 6.44 dd 15.2, 9.9 | 125.4, CH |
| 6 | 7.05 dd 15.2, 10.9 | 146.9, CH |
| 7 | 6.37 dd 15.0, 10.9 | 129.5, CH |
| 8 | 6.16 dt 15.0, 6.9 | 143.2, CH |
| 9 | 2.17 m | 32.3, $CH_2$ |
| 10 | 1.40 m; 1.28 m | 28.0, $CH_2$ |
| 11 | 1.23 m | 28.6, $CH_2$ |
| 12 | 1.25 m | 31.0, $CH_2$ |
| 13 | 1.27 m | 21.8, $CH_2$ |
| 14 | 0.86 t 6.9 | 14.0, $CH_3$ |
| -OH | 13.89 s | |

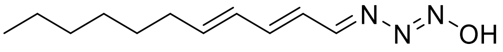


**1** ($^1$H NMR, DMSO-$d_6$ at 900 MHz)

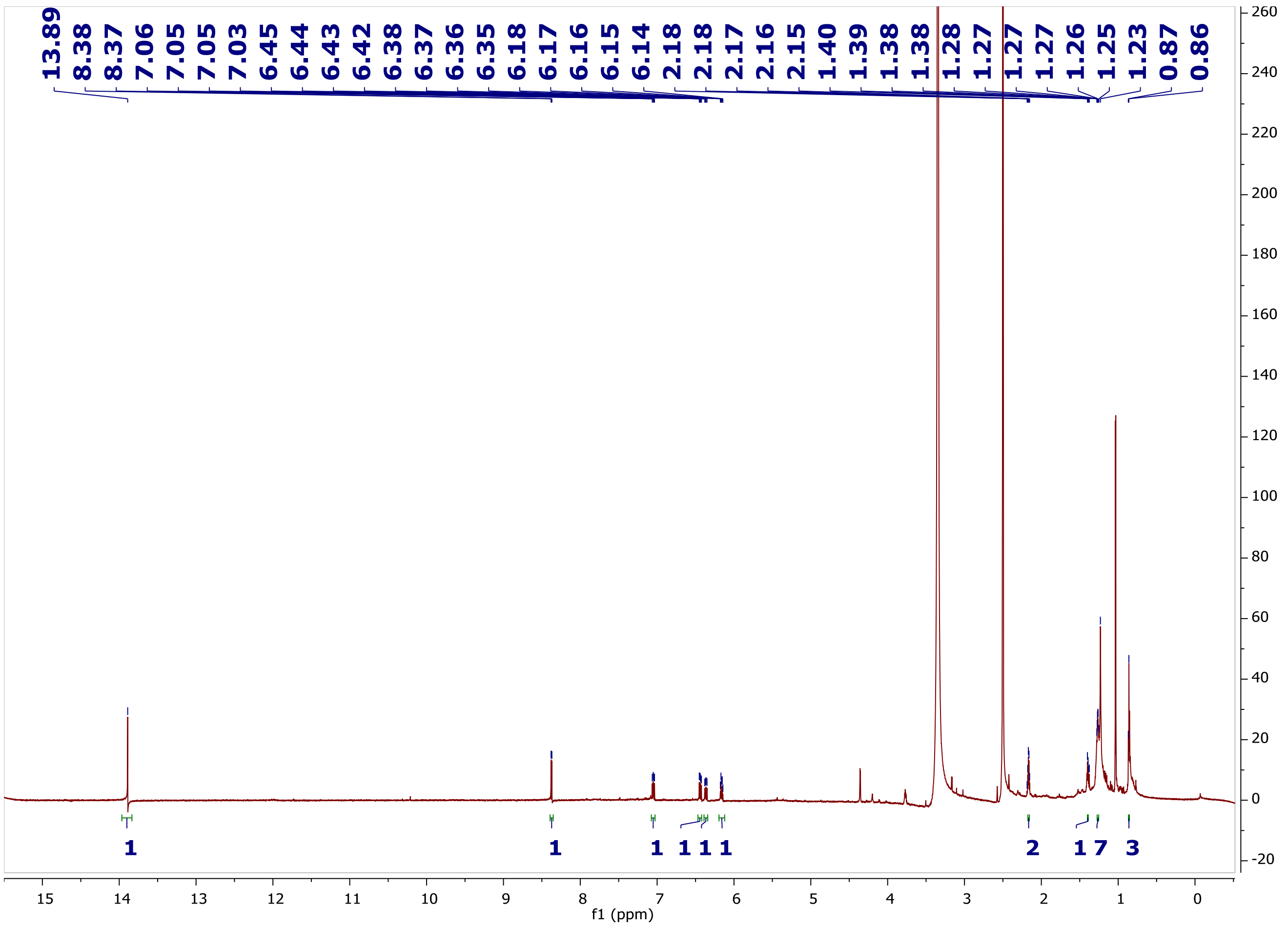

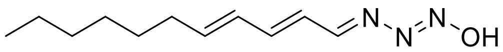

**1** ($^1$H-$^1$H COSY, DMSO-$d_6$ at 900 MHz)

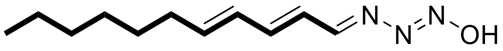

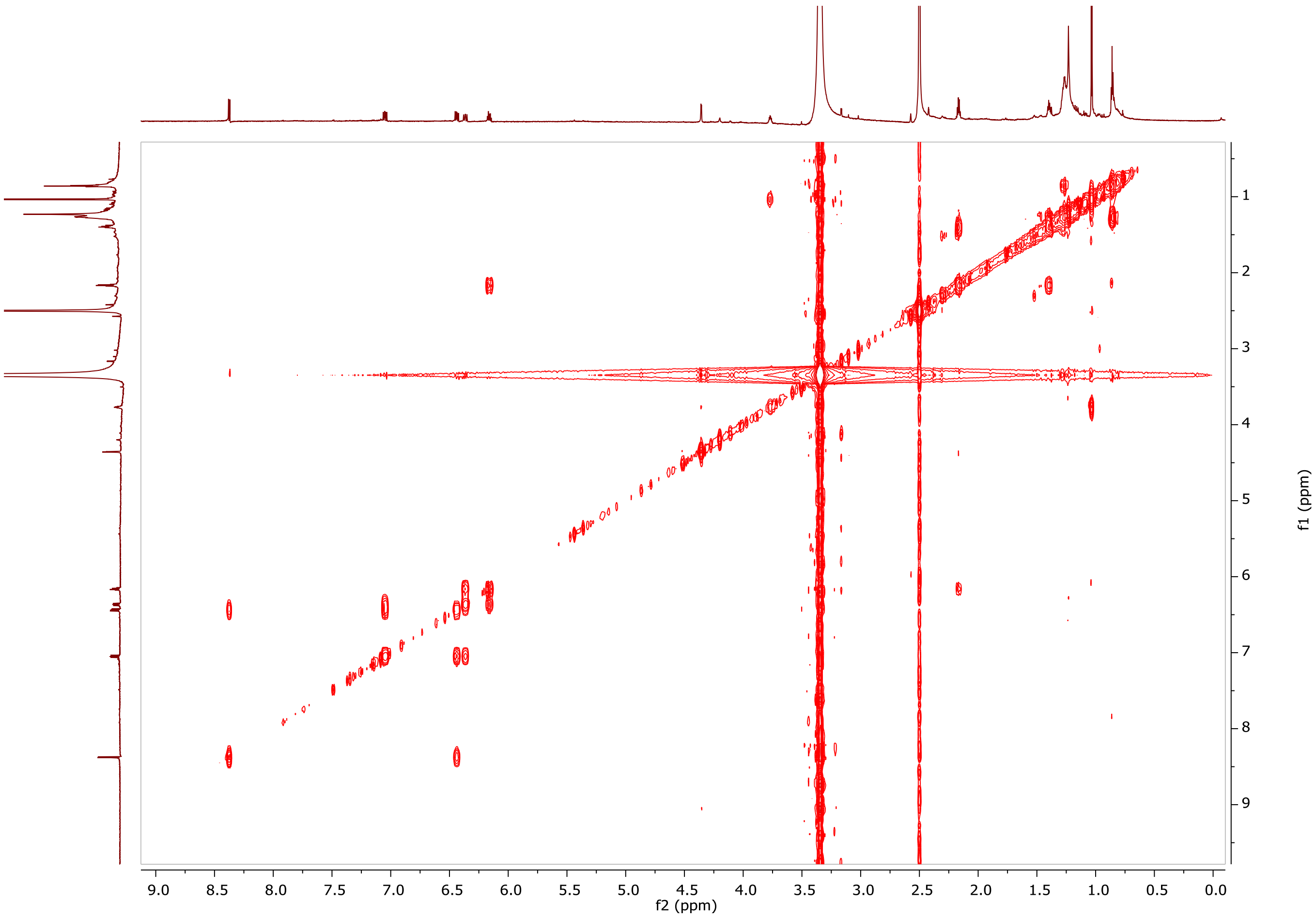

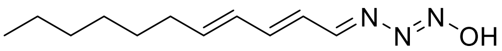


**1** ($^1$H-$^{13}$C HSQC, DMSO-$d_6$ at 900 MHz)

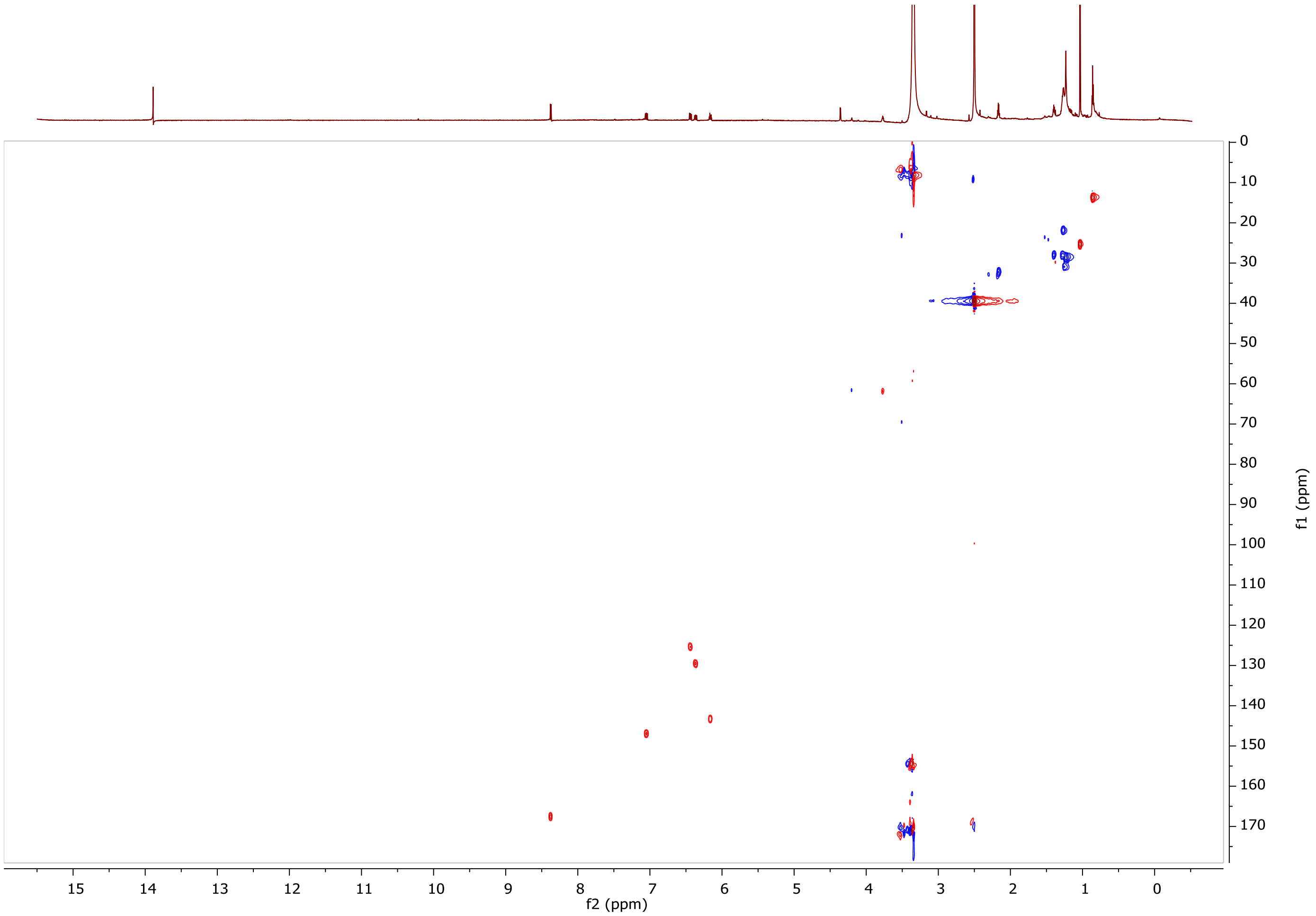

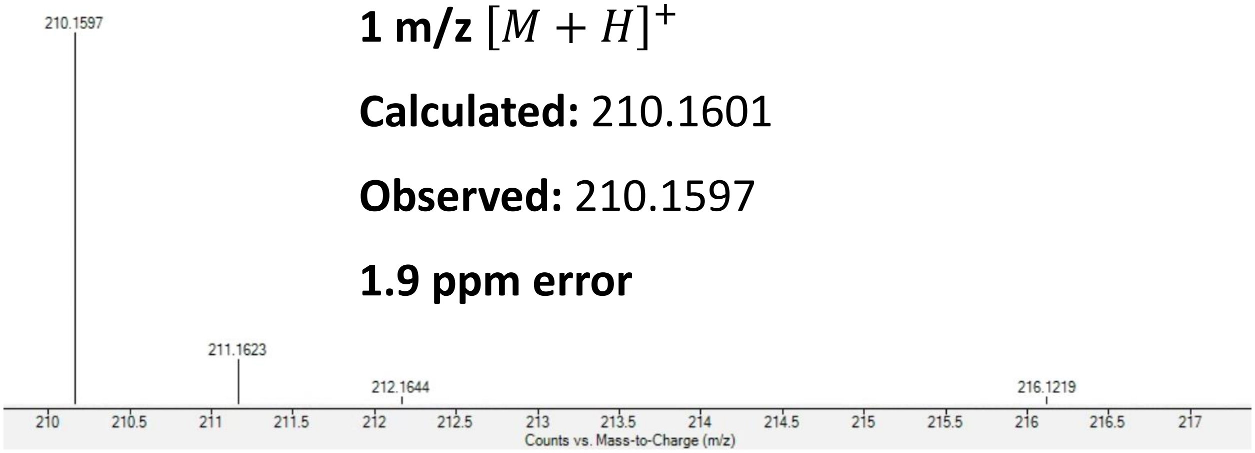


**1 m/z** $[M+H]^+$

**Calculated:** 210.1601

**Observed:** 210.1597

**1.9 ppm error**

**1 UV**

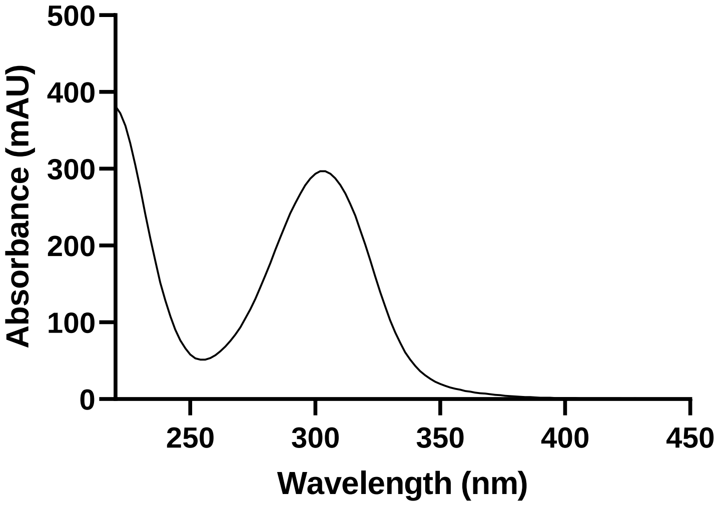
